# Quantifying year-round nocturnal bird migration with a fluid dynamics model

**DOI:** 10.1101/2020.10.13.321844

**Authors:** Raphaël Nussbaumer, Silke Bauer, Lionel Benoit, Grégoire Mariethoz, Felix Liechti, Baptiste Schmid

## Abstract

The movements of migratory birds constitute huge biomass flows that influence ecosystems and human economy, agriculture and health through the transport of energy, nutrients, seeds, and parasites. To better understand the influence on ecosystems and the corresponding services and disservices, we need to characterize and quantify the migratory movements at various spatial and temporal scales.

Representing the flow of birds in the air as a fluid, we applied a flow model to interpolated maps of bird density and velocity retrieved from the European weather radar network, covering almost a full year. Using this model, we quantified how many birds take-off, fly, and land across Western Europe, (1) to track waves of bird migration between nights, (2) cumulate the number of bird on the ground and (3) quantify the seasonal flow into and out of the study area through several regional transects.

Our results show that up to 188 million (M) birds take-off over a single night. Exemplarily, we tracked a migration wave in spring, in which birds crossed the study area in 4 days with nocturnal flights of approximately 300 km. Over the course of a season, we estimated that 494 million (M) birds entered through the southern transects and, at the same time, 251 M left in the northern transects, creating a surplus of 243 M birds within the study area. Similarly, in autumn, 544 M more birds departed than arrived: 314 M birds entered through the northern transects while 858 M left through the southern transects.

Our study show-cases the potential of combining interdisciplinary data and methods to elucidate the dynamics of avian migration from nightly to seasonal and yearly time-scales and from regional to continental spatial scales.

## 1 Background

The sheer numbers of migratory birds create huge biomass flows (Alerstam, 1993; Dokter et al., 2018; Hahn, Bauer, & Liechti, 2009) that impact ecosystem functions and human economy, agriculture and health through the transport of energy, nutrients, seeds, and parasites (Bauer & Hoye, 2014). To understand these influences on ecosystems and make use of, or avoid, the resulting services and disservices, we need year-round and continental-wide monitoring of migratory fluxes and their quantification at various spatial and temporal scales. Continental networks of weather radars are increasingly becoming essential tools for monitoring large-scale migratory movements (Bauer et al., 2019). However, most studies so far have focused on specific stages of the migration journey: migratory flights (e.g., Dokter et al., 2018; Horton et al., 2020; Nilsson et al., 2019; Nussbaumer et al., 2019; Van Doren & Horton, 2018), or stop-overs (e.g., Buler et al., 2017; Cohen et al., 2020; McLaren et al., 2018). Yet, none have explicitly considered and differentiated between the three successive stages of take-off, flight and landing, and we therefore lack a comprehensive model of the entire migratory journey.

To integrate migratory take-off, flight and landing into a single framework, we adopted a methodology from fluid mechanics. While novel in aeroecology, fluid mechanics methods have been applied in ecology before, for instance, the concept of permeability from Darcy’s Law to calculate species movement rates (Jones, Watts, & Whytock, 2018) or a hydrological residence time model to estimate the stop-over duration of migratory birds (Drever & Hrachowitz, 2017).

Here, we treat the nocturnal broad-fronted migration as a fluid and model bird density as a conservative quantity using the continuity equation (e.g., Pedlosky, 1987). More specifically, we combine interpolated maps of bird density and velocity into a discretised flow model (Figure 1). Since we assume that the biomass of birds moving from one grid cell to another is conserved, any change of bird density (in the air) must be explained by movements to and from the ground. Thus, we can quantify how many birds take-off, fly, and land at any given time and location. In subsequent steps, we use the resulting maps of take-off and landing to (1) track waves of bird migration between nights across Europe, (2) estimate the accumulation (i.e. changes in numbers) of birds on the ground throughout the year and (3) quantify the seasonal flow in and out of the study domain through several transects.

**Figure 1.**
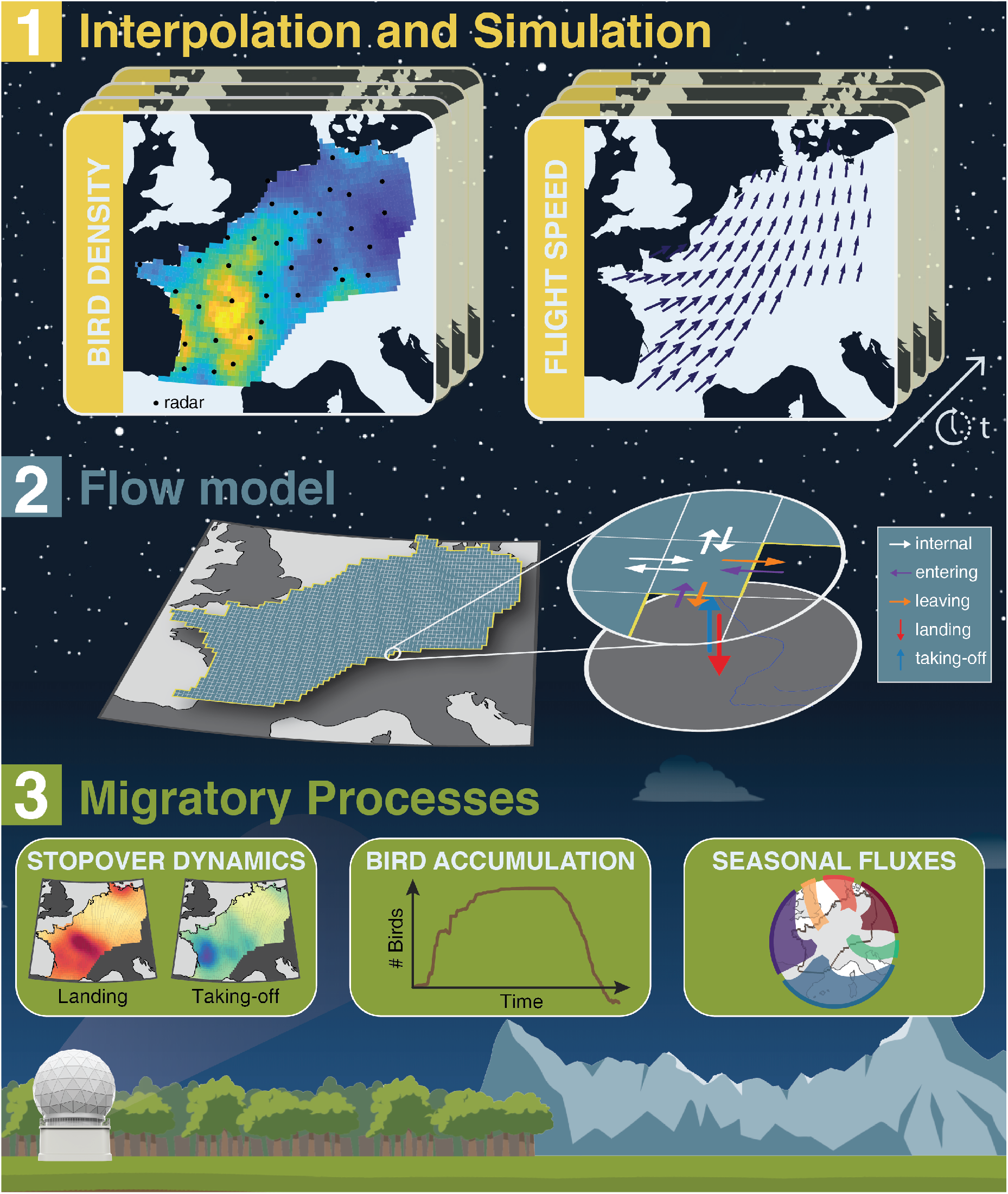
Overview of the methodology for modeling nocturnal bird migration as a fluid flow at the continental scale. **1. Interpolation and Simulation** (section 2.2). First, we interpolate vertical profile time series of bird density and velocity field measured by weather radar data into continuous spatio-temporal maps following Nussbaumer et al. (2019). **2. Flow model** (section 2.3) Then, using the interpolated data in a flow model allows us to estimate the number of birds entering, leaving, taking off from and landing in each grid cell at each time step. **3. Migration Processes** (section 2.4) The resulting maps of take-off and landing birds allow us to investigate the spatio-temporal variation of stop-over, the accumulation of birds on the ground, and the geographic variation in the seasonal fluxes of migrating birds.

## 2 Methodology

### 2.1 Data

We used the data from 37 weather radars in France, Germany, the Netherlands and Belgium operating between 13 February 2018 and 1 January 2019. This dataset is currently the longest available time series over a large part of Western Europe. It consists of vertical profiles of bird density [birds/km^3^], flight speed [m/s] and flight direction [°] which were generated with the vol2bird software (Dokter et al., 2011) and are available on the ENRAM repository (ENRAM, 2020) at a 5 min x 200 m (0-5000m a.s.l.) resolution. Similar to previous studies (Nilsson et al., 2019; Nussbaumer et al., 2019), the vertical profiles were cleaned as follows (supplementary material 1.2). First, we eliminated high-reflectivity contamination (e.g. rain and ground scatter) using a dedicated graphical user interface. Then, we removed contamination from slow moving targets with low-reflectivitiy such as insects or snow based on standard deviation of radial velocity and air speed (Nussbaumer, Schmid, Bauer, & Liechti, 2021). Finally, we vertically integrated bird density and flight speed (i.e. volumetric to areal) while (1) accounting for the impact of local topography on the surveyed volume, and (2) simulating bird density in the volume of air below the altitude surveyed (supplementary material 1.3).

### 2.2 Interpolation and Simulation

Since the radars provide point observations (averaged over a 5-25 km radius around the radar location), we interpolated bird density [birds/km^2^] into a spatio-temporal grid using the methodology developed in (Nussbaumer et al., 2019). The bird velocity field (i.e. the vector field of birds’ flight speed and direction) was interpolated for the two N-S and E-W components separately using a similar methodology. Adjustments of the interpolation method to a year-round dataset and to a velocity field are detailed in supplementary material 2.

The interpolation grid was defined between latitudes 43° and 55° and longitudes -5° and 16°, with a resolution of 0.25° and between 13 February 2018 and 1 January 2019 with a resolution of 15 min in time. Similarly to Nussbaumer et al. (2019), grid cells were excluded if (1) they were located over a water body or above 2000 m a.s.l, (2) they were more than 150 km away from a weather radar, (3) they spanned over day time (i.e., from sunrise to sunset), or (4) rain intensity exceeded 1mm/hr (interpolated from ERA5 dataset from Copernicus Climate Change Service (C3S) (2017)). Nights without data were excluded from the interpolation (5 nights in early April, and 34 nights in July-August). The resulting interpolation maps can be visually explored at www.birdmigrationmap.vogelwarte.ch/2018.

To correctly calculate aggregated measures (e.g. average bird density or sum of birds take-off) and their uncertainties, we generated 500 geostatistical simulations of bird density (Lantuéjoul, 2002; Nussbaumer et al., 2019).

### 2.3 Flow Model

Based on the principle of mass conservation, the continuity equation (Equation 1) describes the transport of a conserved quantity (e.g., bird density): the rate of change of this quantity is equal to its flux into and out of a given volume (e.g. sky). The equation can also include a source/sink term, which accounts for the appearance (and disappearance) of the quantity (e.g., take-off and landing). The differential form of the continuity equation for bird density *ρ* [birds/km^2^] is

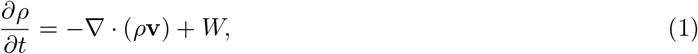

where **v** = [*v*_*lon*_, *v*_*lat*_] is the bird’s velocity field [km/hr] along latitude and longitude and *W* is the source/sink term [birds/hr/km^2^] and ∇ denotes the vector differential operator. The continuity equation is discretised with a Forward Time Centered Space (FTCS) scheme (Roache, 1972). The source/sink term can be computed for each cell (*i, j, t*) with

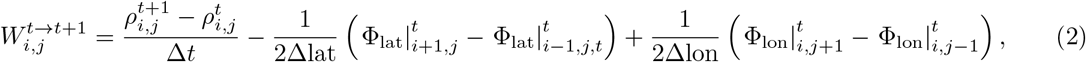

where **Φ** = *ρ***v** = [Φ_lon_, Φ_lat_] is the flux term expressed in [birds/km/h] and discretised in longitude, latitude and time with the indexes *i, j, t* respectively. Δlon, Δlat and Δt are the grid resolution in time, longitude and latitude respectively.

We applied this model to bird migration by using the spatio-temporal maps of bird density (*ρ*) and flight speed vector (**v**) derived from geostatistical simulations (Section 2.2). The local fluxes were computed for each grid-cell by multiplying the density with the flight vector and then linearly interpolated to the grid cells’ boundaries for both the longitudinal and latitudinal components. As the grid was defined in equal latitude and longitude intervals, the resolution Δlon in km varied along the latitude axis. Finally, using Equation 2, the source/sink term was computed for each grid cell at each time step as the change of bird density over time minus the spatial difference of fluxes. The source/sink term *W* was composed of birds taking-off and landing (within the study area) which can be separated according to the sign of *W*. Indeed, as the reference of the mass balance was the sky, positive values of *W* correspond to birds taking-off while negative values correspond to landing. Additionally, the values of the fluxes at the study area’s boundaries were extracted as the number of birds entering (positive) and leaving (negative) the study area. We uses the 500 simulations to produce space-time maps of (1) take-off and landing [birds/ km^2^] and (2) fluxes in lat-lon [birds/hr/km] at the boundaries of the study area.

### 2.4 Migratory Processes

The resulting maps were processed to address specific ecological questions. We were particularly interested in characterizing and quantifying nightly migration pulses and stopovers, the accumulation of birds on the ground, and the seasonal migration flows. To achieve this, we processed each of the 500 realisations as follows:

- **The nightly migratory pulses and stopovers** were calculated by summing the take-off and landing movements separately over each night, and by visually comparing the maps of landing in the morning with those of take-off the following evening.
- **The year-round accumulation of migratory birds on the ground** was quantified by first aggregating the four fluxes (take-off, landing, entering, leaving) over the whole study area and for each night. Then, the nightly change in the number of birds on the ground was computed as the difference between landing and take-off, or equivalently, between entering and leaving. The cumulative sum of these daily changes corresponds to the number of birds that remained on the ground. We arbitrarily set the starting value of the accumulation to zero because the initial number of (resident and/or wintering) birds on the ground is unknown.
- **The seasonal flow of bird migration** is quantified by summing the fluxes of birds entering and leaving the study area over spring (February-June) and autumn (August-December). To capture the variability of movements across Europe, we defined six transects along the boundary of the study area according to the major flyways: United Kingdom, the North, the East, the Alps, Spain and the Atlantic (Figure 4).

## 3 Results

### 3.1 Nightly migratory pulses and stopovers

For illustration purposes, we selected a well-defined migration wave spanning from 6 to 10 April, during which birds moved from south-western France to north-eastern Germany (Figure 2). The nightly averaged bird density and flight speed was highest between the main take-off and landing areas. More importantly, one night’s landing and the following night’s take-off were in good agreement, demonstrating that the data and proposed methodology can accurately track a wave of migration over several days. This agreement was particularly striking in this example because birds did not stopped over, migrating every nights. The crossing of the study area in approximately 4 nights corresponds to nightly migratory bouts of around 300 km. On April 10, one radar in south-west France detected a new wave arriving.

**Figure 2.**
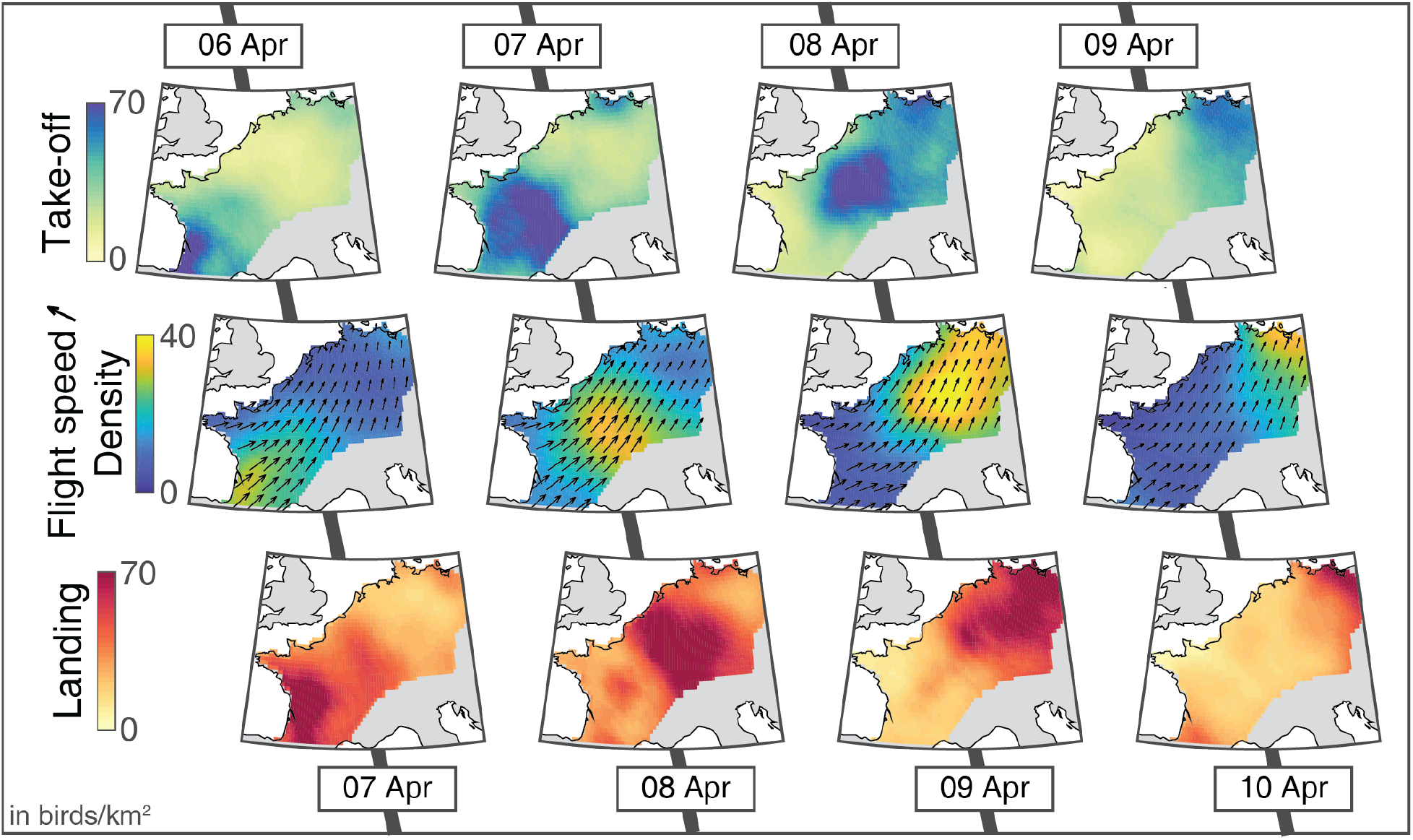
Consecutive phases of take-off (top row), flight (middle row) and landing (bottom row) of bird migration between 6 and 10 April 2018. Take-off and landing maps show the sum of take-off and landing over the entire night, respectively, while the density and flight speed maps show the average over the night.

### 3.2 Year-round accumulation of migratory birds on the ground

Summing the flow at the daily (or nightly) scale allows us to characterize the year-round changes in numbers of migratory birds on the ground (Figure 3, a seasonal sum of the bird movements (take-off, landing, entering, and leaving) is provided in the supplementary material Figure 3.1). The number of birds on the ground rises steeply in March with almost 200 million more birds entering in the study area than leaving it, e.g., to migrate further North or East. Numbers are declining from August onwards, and plummeted in mid-October. The number of bird on the ground became negative in autumn because our methods did not explicitly account for reproduction and mortality, therefore the birds leaving the area in autumn included the new generation. The spring migration period was shorter and more condensed (March - May) than the autumn migration (August to mid-November) (Figure 3), with 50% of all take-offs taking place during 19 nights in spring and 29 nights in autumn. At peak migration, we estimated 118 (Q5-Q95: 99-137) million birds taking off in a single night in spring (30 March – 1 April) and 148 (133-164) million in autumn (17-18 October). Prior to these two peak migration events, we observed that the accumulation curve of bird on the ground flattened, indicating a temporarily reduced migratory traffic possibly due to unfavorable weather conditions (”Zugstau”).

**Figure 3.**
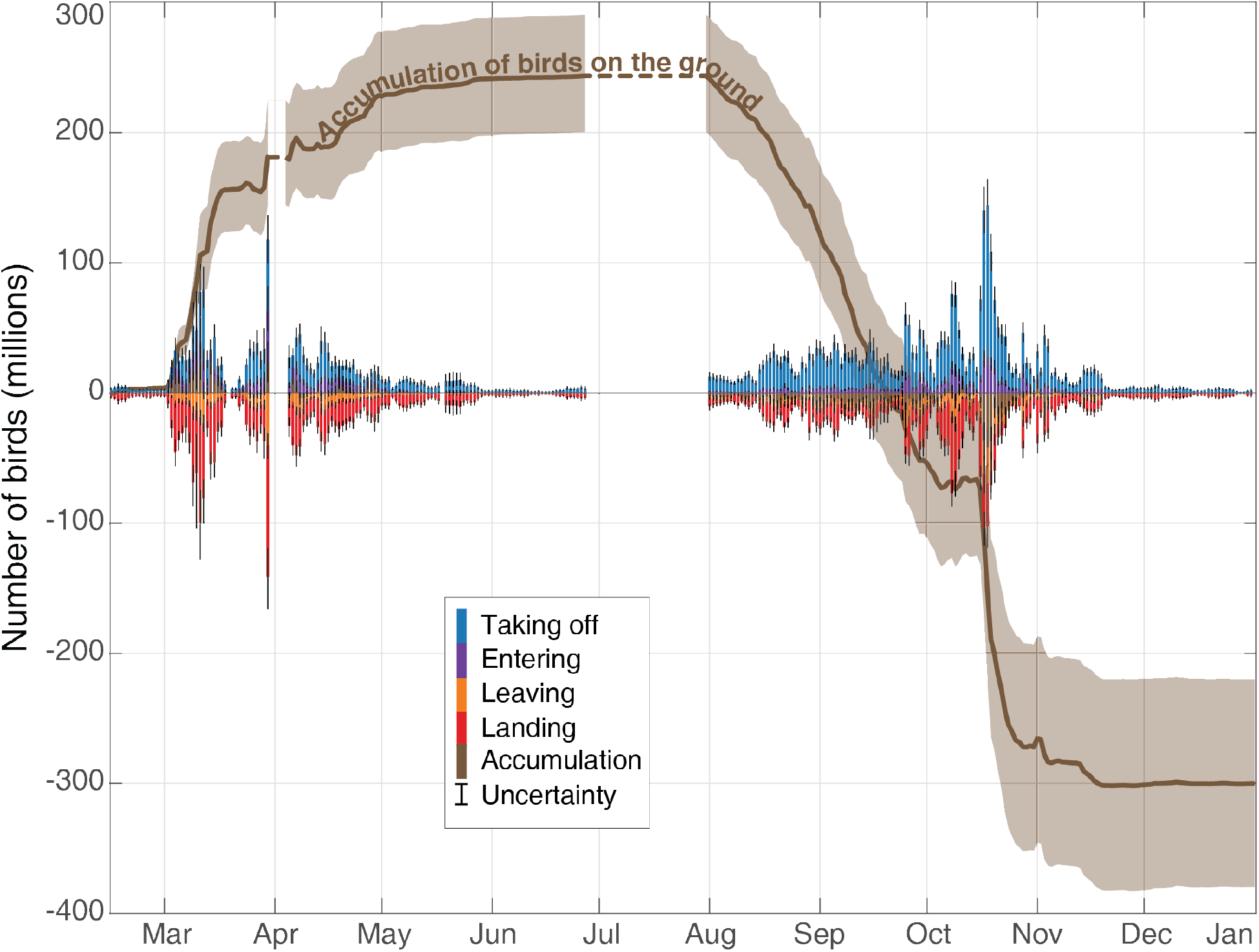
Time series of the daily number of birds taking-off (blue) and landing (red), and entering (purple) and leaving (orange) the study area. The changes in the number of birds on the ground and their cumulative sum (brown line) is calculated as the difference between the number of birds landing in, and taking-off, from the study area. The uncertainties (Q5-Q95) are illustrated with fine black line on the bar plots and with shaded area for the cumulative time series. The dotted lines denote absence of data.

### 3.3 Seasonal Flow

The bird migration in the study area (both in spring and in autumn) was mainly directed between Spain and Eastern Germany (Figure 4). Indeed, even the migration through the Atlantic transect mostly comprises birds crossing the Bay of Biscay from/to Spain.

**Figure 4.**
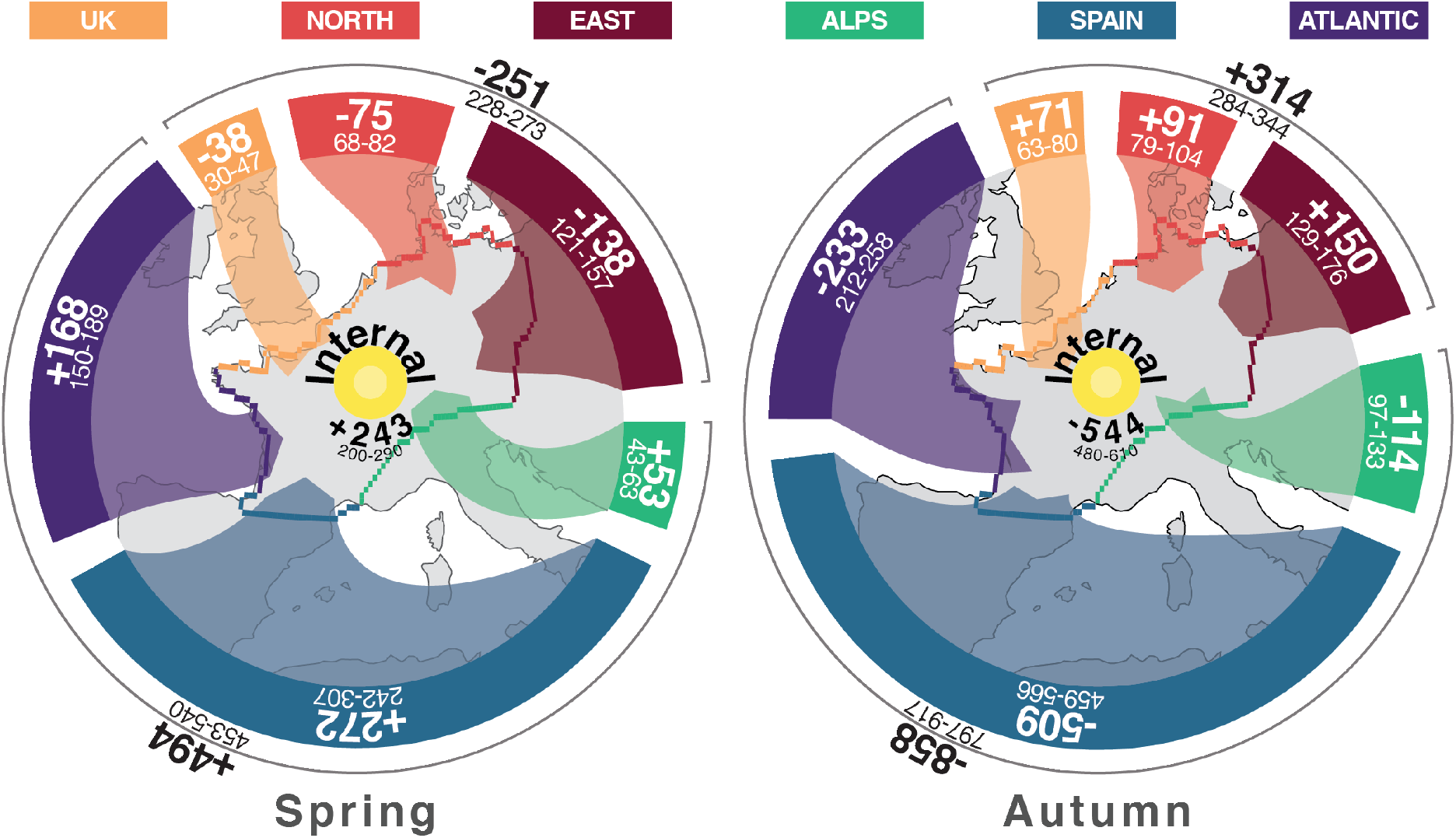
Bird migration flows (in millions of birds) in spring and autumn were aggregated along six transects representing the major flyways. The direction of movement, i.e. into or out of the area, is indicated by the arrows and the sign (+/-) of the mean numbers (bold). The accumulation within the study area results from summing all inward and outward fluxes. Uncertainty for all estimates is provided by their Q5-Q95 ranges.

In spring, 494 (Q5-Q95: 453-540) million birds entered the study area through the southern transects (Alps, Spain and Atlantic) and, at the same time, 251 (Q5-Q95:228-273) million left it in the northern transects (UK, North and East), thus creating a surplus of 243 (Q5-Q95:200-290) million birds that remained within the study area (Figure 4). Similarly, in autumn, 544 (Q5-Q95:480-610) million more birds departed than arrived: 314 (Q5-Q95:284-344) million birds entered through the northern transects while 858 (Q5-Q95:797-917) left through the southern transects. The ratio of the autumn deficit to the spring surplus is 2.2 (Q5-Q95:1.8-2.8), meaning that for one bird staying in spring, two birds left the ground in autumn.

Compared to spring, birds took a more easterly route in autumn, with proportionally more birds flying through the Alps transect (autumn/spring = 114*/*53 = 2.15) than through the Atlantic transect (233*/*168 = 1.4). Moreover, nearly the same number of birds crossed the East transect in autumn and spring (150*/*138 = 1.1). Overall, this pattern could be indicative of a clockwise loop migration where birds migrate to their breeding areas via the Iberian Peninsula in spring and fly to their non-breeding areas further East in autumn.

The seasonal fluxes per transect summarise the number of birds entering and leaving and can therefore cover some fine scale features of migration. Firstly, some transects showed a more unidirectional flow of migrants whereas the entering and leaving fluxes were more balanced for other transects. For instance, 91% of all movements across the Spanish transect are in-movements in spring, i.e. most birds enter the study area rather than leave it, and similarly, 90% of all movements in autumn are out-movements of birds leaving the study area. In contrast, movements across, e.g., the Alps transect were less uni-directional in both seasons with only 63% of all movements in spring being movements into the study area (entering) and similarly, 72% of all movements in autumn were movements out of the study area (leaving) (Figure supplementary material 3.2). Secondly, the timing of migration differs between transects (Figure supplementary material 3.2), with, e.g. Spain and Atlantic transects seeing more than half of its migration before mid-March, while only 20-30% of birds have crossed the East and North transects at that time.

## 4 Discussion

In this study, we presented a novel methodology inspired from fluid dynamics to model the flow of nocturnal migrants, from take-off, during nocturnal flight, to landing. The model produces high-resolution maps that allow investigating the dynamics of migratory movements at various spatial and temporal scales. We used the largest dataset available on the ENRAM data repository to characterize and quantify nightly, seasonal and year-round migration patterns over most of Western Europe.

### 4.1 Model

The model presented in this study builds on the methodology developed in (Nussbaumer et al., 2019), which interpolates point observations of bird densities measured by weather radars into continuous maps. We used these maps as the input for a flow model by considering bird migration as a fluid. This allows us to extract more dynamic information about bird movements and, in particular, their take-off and landing.

The approach used in this study models bird flow (i.e. average bird movement) rather than individual birds. Indeed, the weather radar data consist of bird density and flight speed averaged over a 25km radius. Therefore, the estimated flows cannot capture the properties or behaviour of individual birds. For instance, individual speed are typically be higher than the speed of the flow and the movement directions of individual birds is more variable than the direction of the flow. Similarly, the modeled flow is unable to track separately multiple bird populations simultaneously migrating in different direction.

Throughout the methodology, we identified the sources of errors and tracked the corresponding uncertainties to reliably estimate the ranges of the model outputs. Despite our best efforts to clean the data (supplementary material 1.2), there is an inherent uncertainty in the weather radar data (e.g. ground scattering, measurement errors, radar biases). We partially accounted for the data uncertainty at low altitude when we generated uncertainty range in the vertical integration (see supplementary material 1.3). For more details on the data quality of weather radar, we refers the readers to the assessment and comparison found in Liechti et al. (2019); Nilsson et al. (2018); Nussbaumer et al. (2019). We handle these unknown errors in the geostatical framework (i.e. interpolation) by fitting a nugget effect in the spatio-temporal model (more detailed in supplementary material 2). The nugget effect essentially fits a random noise to the data (e.g., corresponding to the data error), which then, permits the interpolated value to diverge from a datapoint. In addition to the data error (i.e., difference between ‘true’ passage and measured passage), the nugget effect also models small-scale variability/discontinuity in bird density not covered by the dataset (<50-100km) caused by, e.g., geographical features (mountains, rivers, sea), weather conditions (e.g. rain). We percolated these uncertainties throughout our methodology by generating 500 simulation of bird density representing the range of possible values (section 2.2), run in the flow model on each of them, and finally are able to provide for each output (e.g. number of bird on the ground) a distribution of the possible value.

### 4.2 Stopover

In this study, we demonstrated how waves of bird migration at the regional scale can be tracked over multiple nights (Figure 2). Our flow model links birds on the ground with birds in the air and can thus quantify the fluxes of take-off, flight and landing. Looking ahead, this example suggests that a forecast system based on a flow model of bird migration could accurately predict bird landings during the night, and perhaps, on a longer term, take-off and landing over a few days.

Our method can compute the rates of both take-off and landing in higher spatial and temporal resolution than earlier approaches (e.g. 3hrs after sunset in Buler and Diehl (2009), interpolation at civil twilight in Buler and Dawson (2014); Buler et al. (2012) or at maximum density within two hours after sunset in Aurbach, Schmid, Liechti, Chokani, and Abhari (2020)) - a feature that will be particularly useful in follow up studies that link movements and stop-overs to geographical features or short, intense weather events.

Although our model can identify the places and times where birds stop-over, other aspects of stopover dynamics such as stopover duration or survival remain to be tackled in future multidisciplinary studies. The main obstacle to addressing stopover dynamics is the inability to track birds during the day, i.e. which of the birds landing one day are the ones taking-off the following day(s). A similar problem appears at the seasonal scale, where we cannot differentiate birds that are wintering, breeding or passing from the birds landing or taking-off. A potential solution would be to explicitly model stop-over duration with a residence time model (Drever & Hrachowitz, 2017).

### 4.3 Accumulation and seasonal flow

Using our novel methodology and an almost continuous one-year dataset, we assessed the relative changes in the number of birds on the ground. We estimated that in autumn 2018, 858 million birds (Q5-Q95: 797-917) migrated southward through Spain and over the Alps (incl. the Atlantic transect) (Figure 4). The only previous quantification of migrant bird population estimated that between 1.52 and 2.91 billion long-distance migrants leave the entire European continent in autumn (Hahn et al., 2009). Our estimation agrees with these numbers if we consider that our study area (from the British islands to Scandinavia, Finland to Poland, and our study area) corresponds to roughly one third of the European continent as compared to the entire continent in (Hahn et al., 2009). In North America, the number of birds migrating out of the US in autumn was estimated to around 4.72 billion birds (Dokter et al., 2018), which corresponds to an average density of 236 birds/km^2^ (for an area of 19.8 million km^2^). Despite the differences of scale and ecological context, we found a comparable average density of 286 birds/km^2^ when we again assume that a third of the European bird population migrates through the southern transect.

The ratio between autumn and spring fluxes can be used to estimate an index of net recruitment, accounting for both reproduction and mortality (Dokter et al., 2018). For the US, Dokter et al. (2018) estimated a ratio of 1.36 for a transect along the southern border and 1.60 for a transect along the northern border. In our study area in Europe, the resulting indices are 1.74 (Q5-Q95: 1.55-1.94) in the southern transects (Alps, Spain and Atlantic, Figure 4) and 1.26 (1.09-1.42) in the northern transects (UK, North and East, Figure 4). However, such derived values like recruitment critically depend on birds taking similar migration routes in both spring and autumn. If migration routes vary between seasons, e.g. when birds take a more easterly route in autumn, recruitment numbers become distorted. Instead of computing the ratio of migratory birds flying across non-representative transects, we can take advantage of the flow model to estimate a ratio of migratory birds entering and leaving an area of interest, and thereby relate the recruitment index computed over this area to environmental characteristics. For the entire study area, a recruitment index of 2.26 (1.80-2.81) resulted from the ratio between the relative number of birds that have left in autumn (i.e., leaving minus entering) (544 M, Figure 3) and the relative number that have arrived in spring (i.e., entering minus leaving) (243 M, Figure 3). However, as the fluxes of wintering and breeding bird populations cannot be distinguished (see discussion on stopover), this recruitment index also depends on the number of wintering birds that leave the study area in spring and return in autumn with offspring.

Therefore, while this recruitment index can characterize the migratory bird population growth, it cannot separate the influence of breeding and wintering populations. A possible avenue to address this challenge is to combine breeding and/or wintering bird atlas data with our accumulation of birds on the ground. This approach could provide absolute numbers of breeding, passing and wintering birds along with their corresponding recruitment indices.

## Supporting information

Daily Map of Take-off and Landing

## Acknowledgements

We thank Pietro De Anna for initial discussion about applicability of a flow model framework to bird migration, and Mathieu Gravey for the assistance in implementing the Multi-Point Statistics simulation.

This study contains modified Copernicus Climate Change Service Information 2019. Neither the European Commission nor ECMWF is responsible for any use that may be made of the Copernicus Information or Data it contains.

We acknowledge the European Operational Program for Exchange of Weather Radar Information (EUMETNET/OPERA) for providing access to European radar data, facilitated through a research-only license agreement between EUMETNET/OPERA members and ENRAM (European Network for Radar surveillance of Animal Movements).

We acknowledge the financial support from the Globam project funded by BioDIVERSA, including the Swiss National Science Foundation (31BD30 184120), Netherlands Organisation for Scientific Research (NWO E10008), Academy of Finland (aka 326315), Belgian Federal Science Policy Office (BelSPO BR/185/A1/GloBAM-BE) and National Science Foundation (NSF 1927743).

## Authors’ contributions

RN, FL, BS and SB conceived the study, RN, LB, GM designed the flow model, RN implemented the computational framework and performed the analyses, RN, BS and SB wrote the manuscript, with substantial contributions from all authors.

## Data Accessibility

- The Github page of the project (https://rafnuss-postdoc.github.io/BMM/2018/) provides links to the MATLAB files (script and livescript) used for preprocessing, interpolation, flow model and creation of the figures.
- The raw weather radar data were available on the ENRAM repository (ENRAM, 2020) (https://github.com/enram/data-repository).
- The cleaned vertical time series profile are available on Zenodo (Nussbaumer, 2020) (https://doi.org/10.5281/zenodo.3610184)
- The code of the website (https://bmm.raphaelnussbaumer.com/2018) are available on the Github (https://github.com/Rafnuss-PostDoc/BMM-web-2018)

