## Supplementary figures and images for "Quantifying year-round nocturnal bird migration with a fluid dynamics model"

### day_2.png

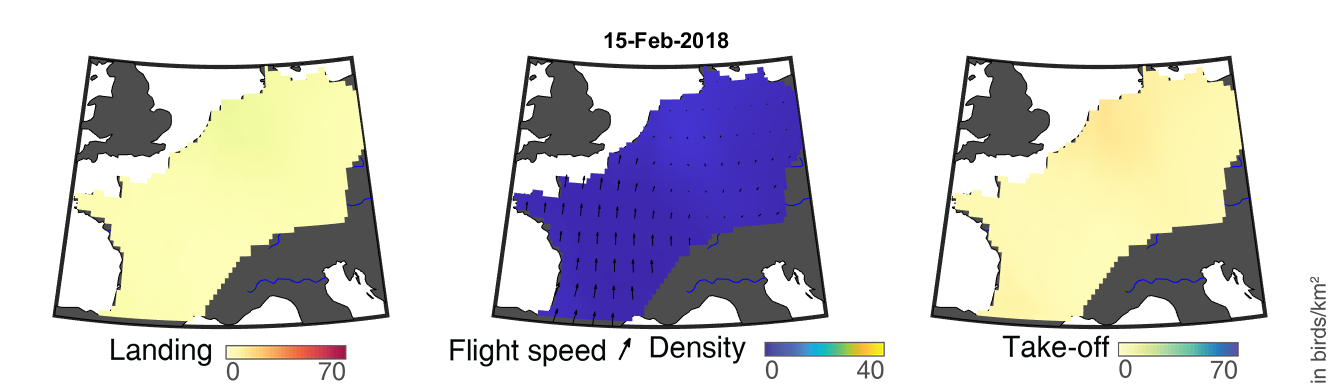

### day_19.png

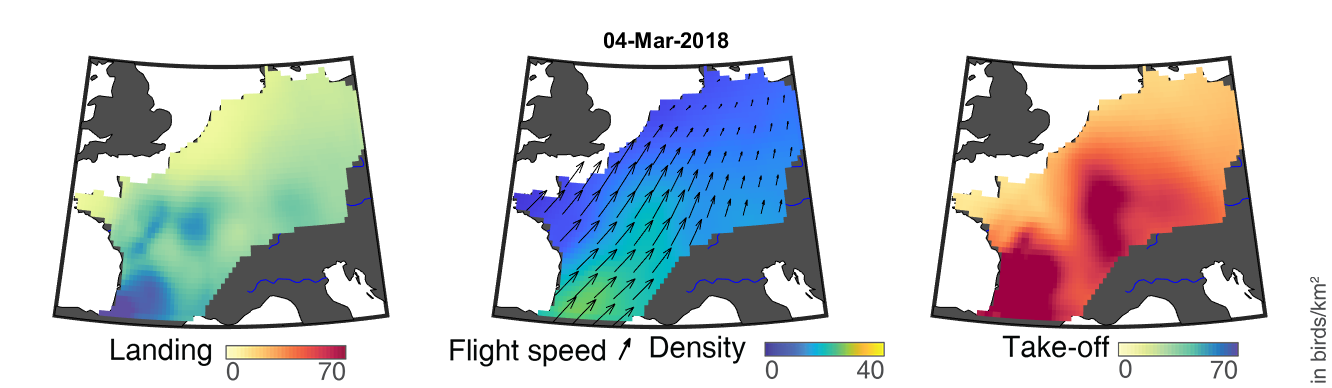

### day_20.png

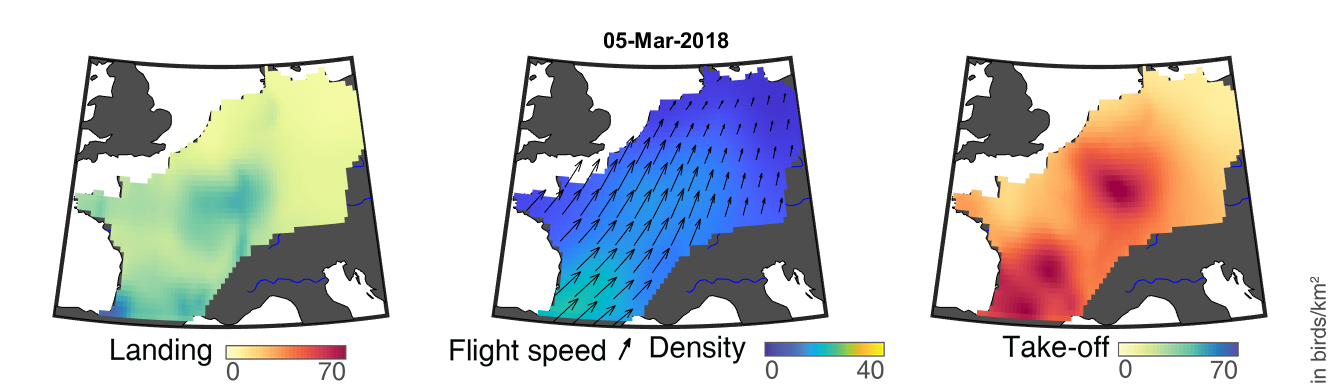

### day_21.png

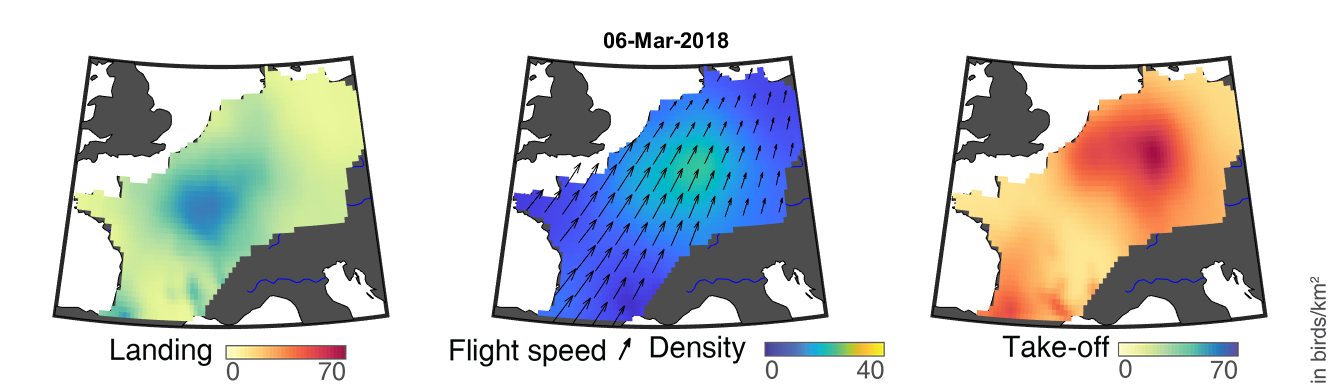

### day_22.png

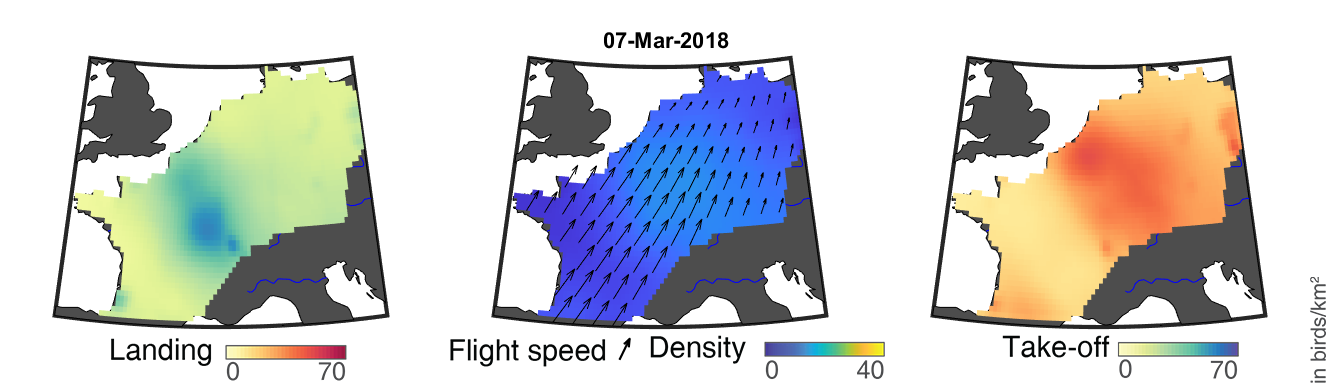

### day_23.png

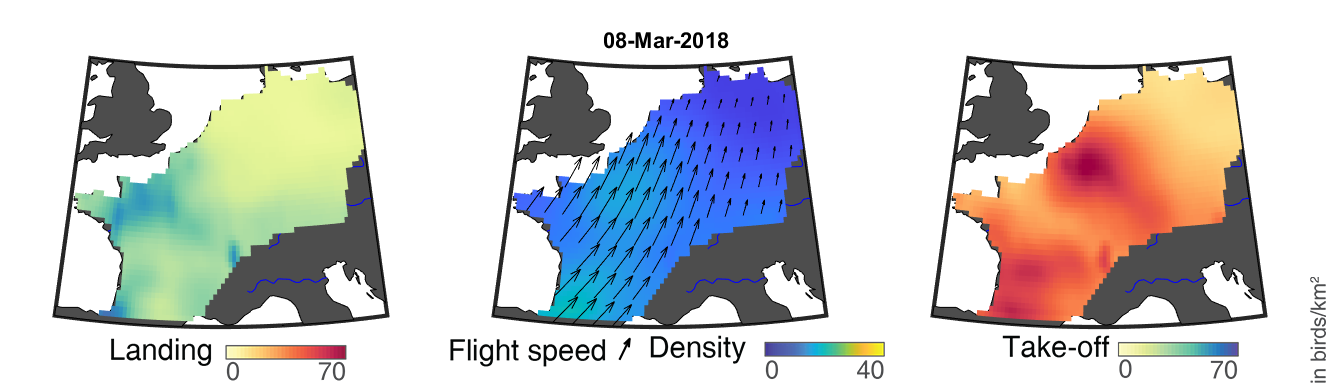

### day_24.png

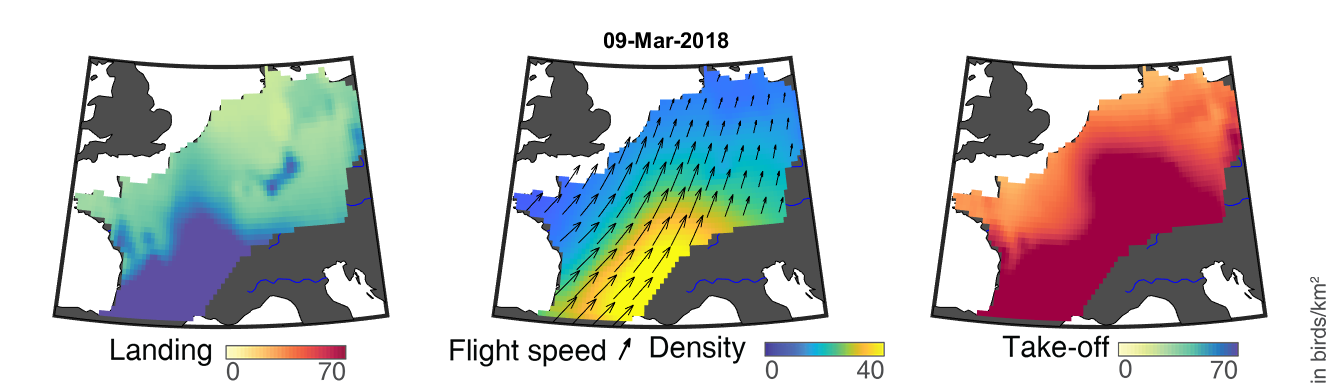

### day_25.png

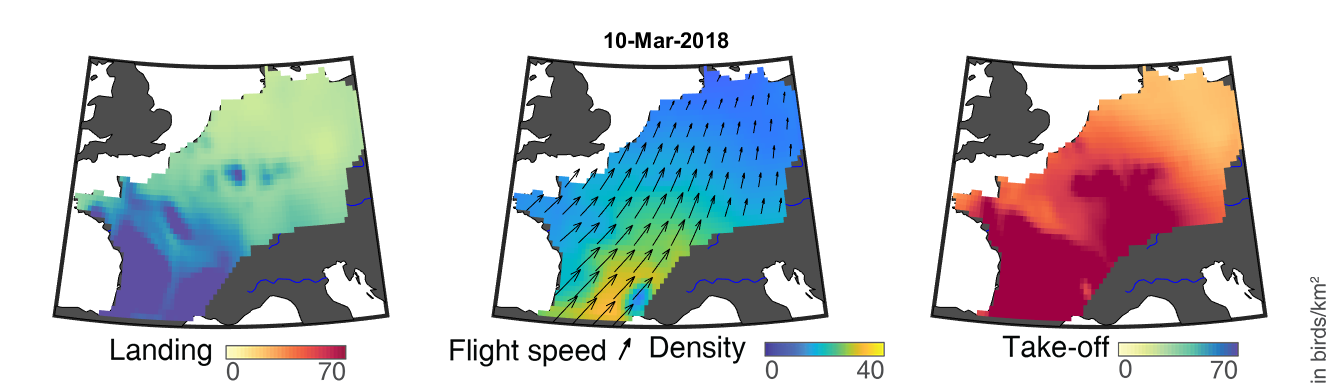

### day_26.png

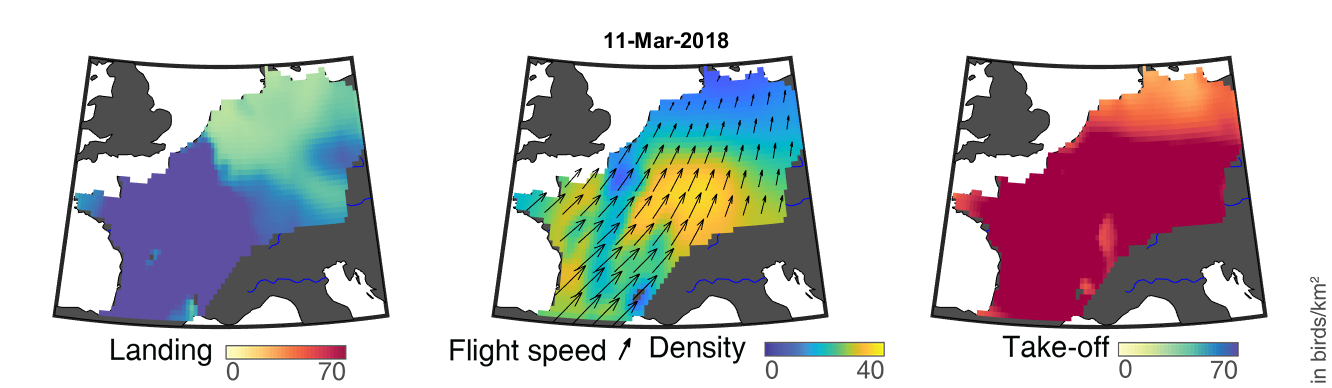

### day_27.png

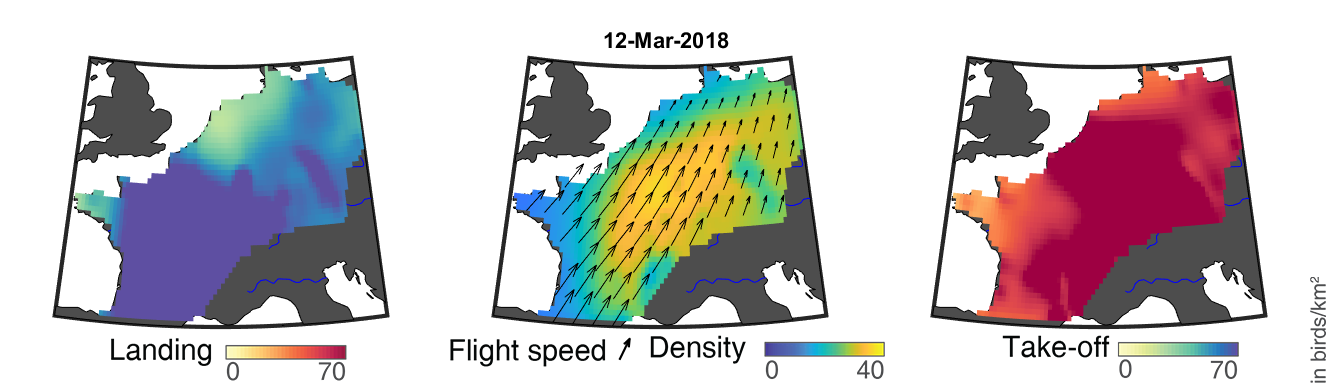

### day_190.png

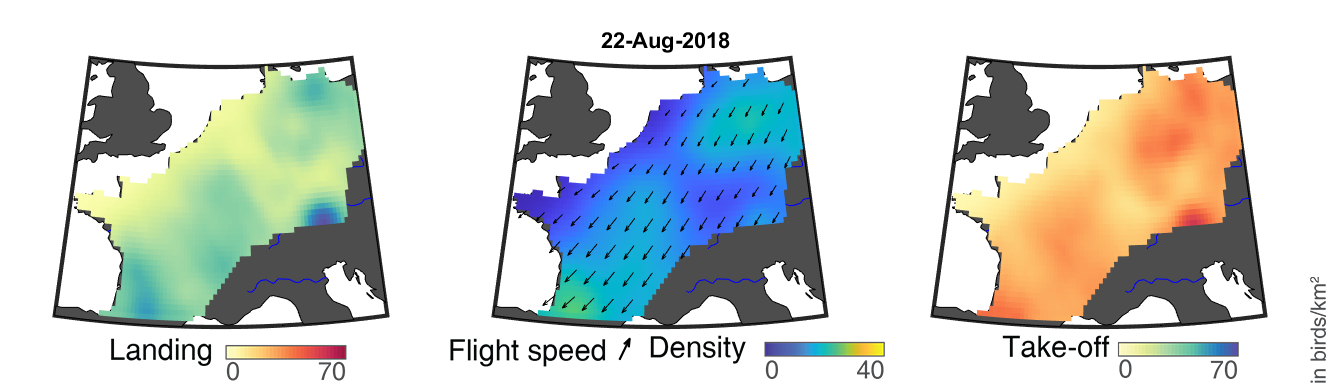

### day_191.png

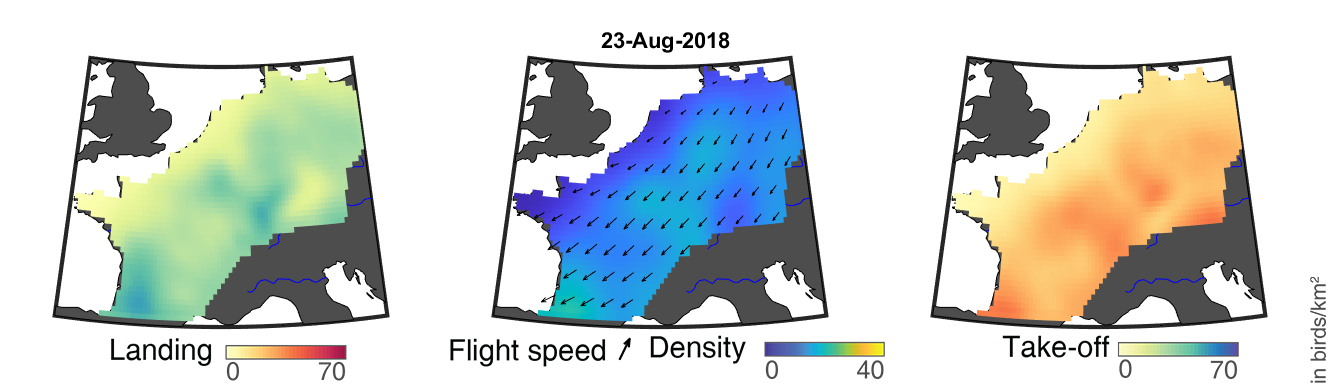

### day_192.png

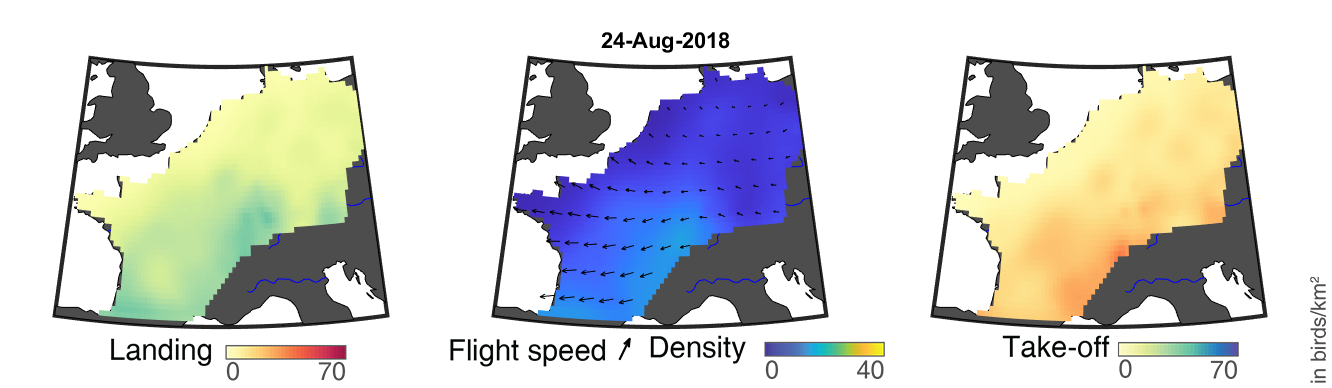

### day_193.png

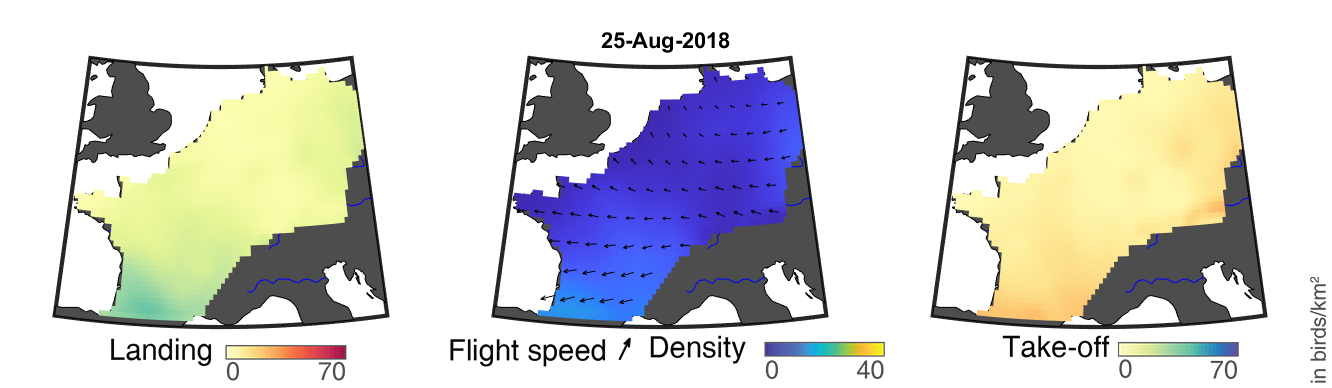

### day_194.png

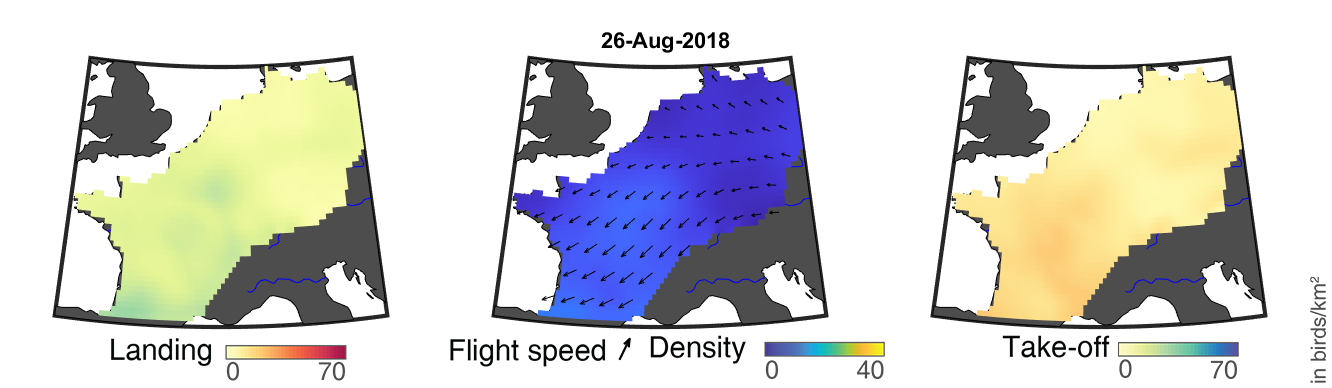

### day_195.png

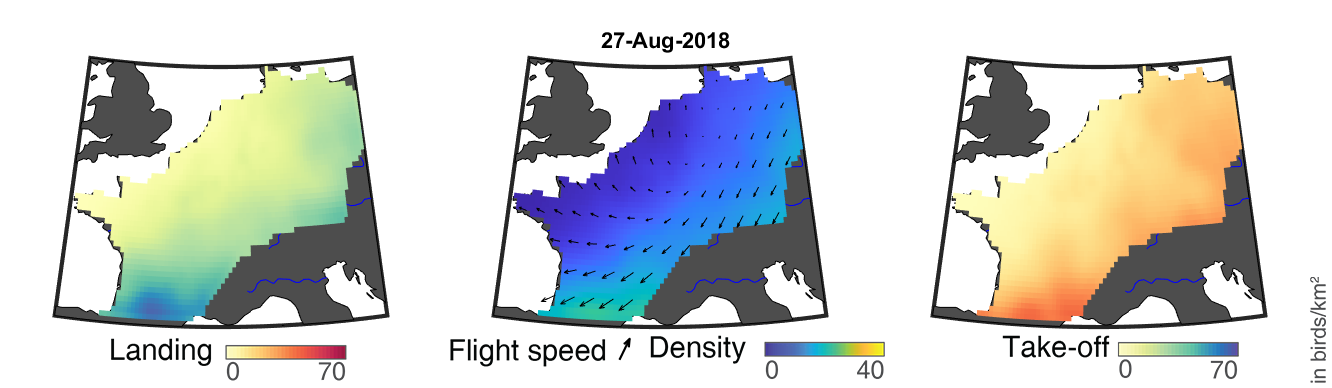

### day_196.png

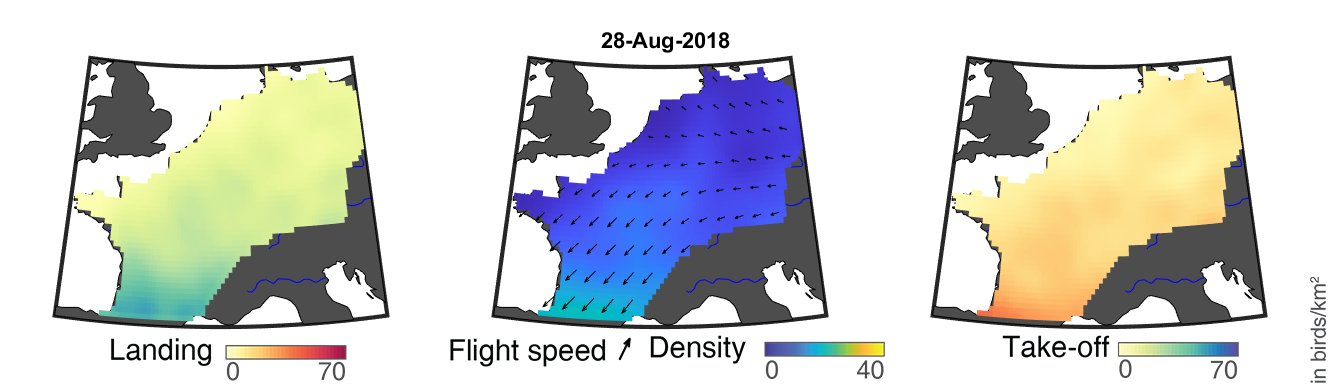

### day_197.png

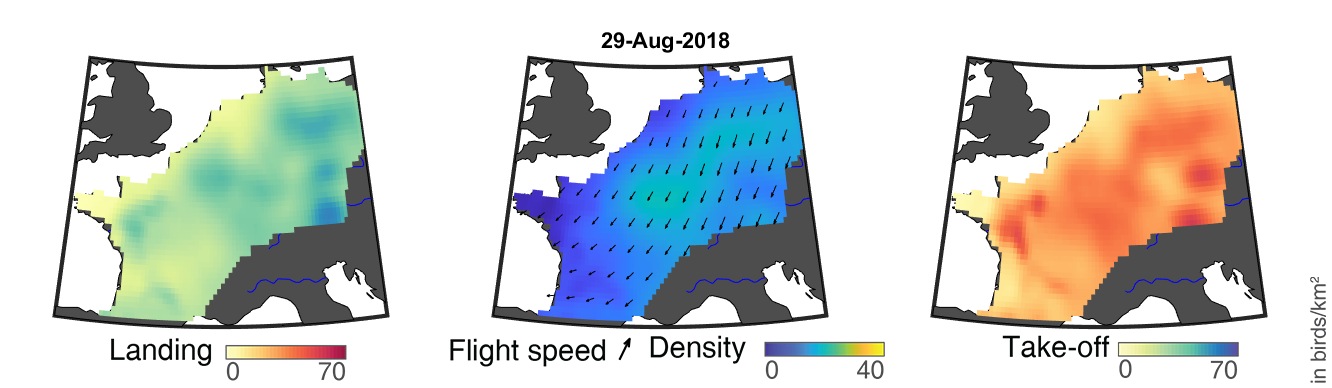

### day_198.png

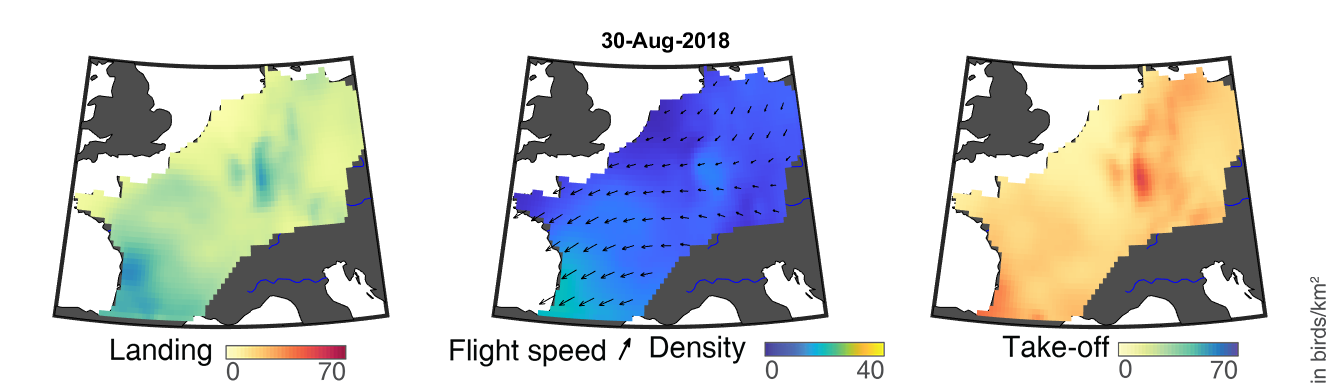

### day_199.png

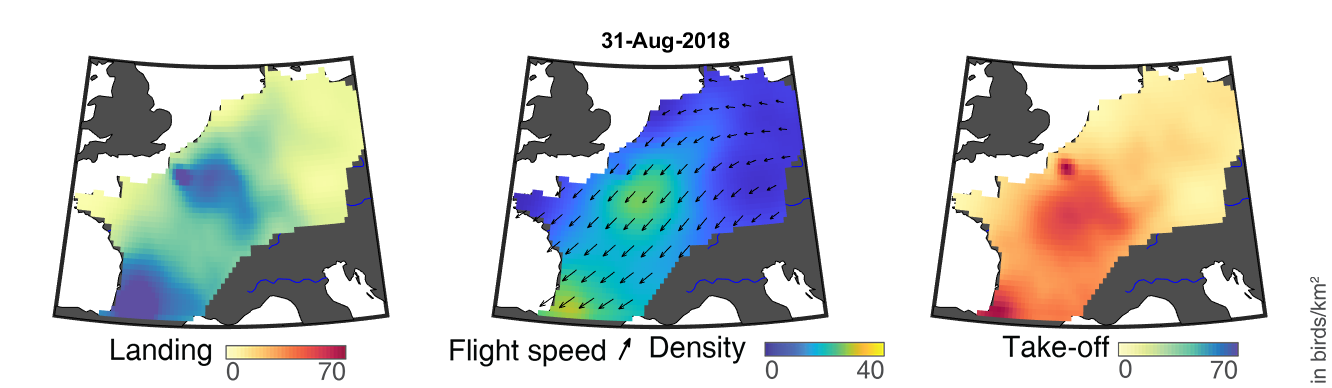

### day_200.png

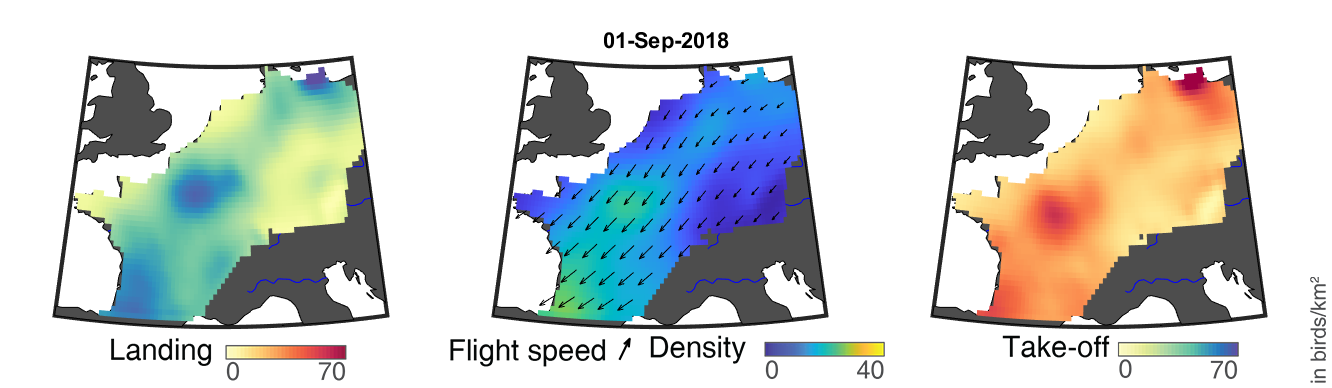

### day_201.png

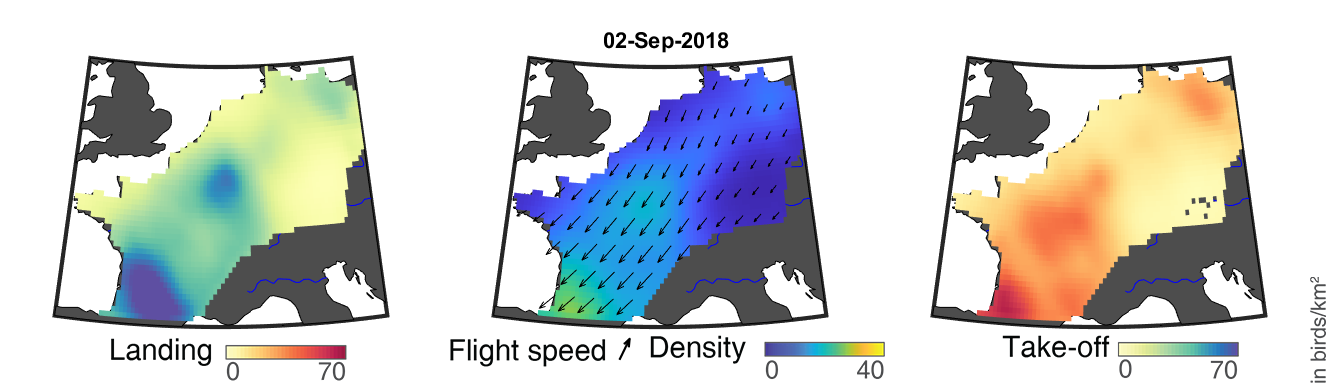

### day_202.png

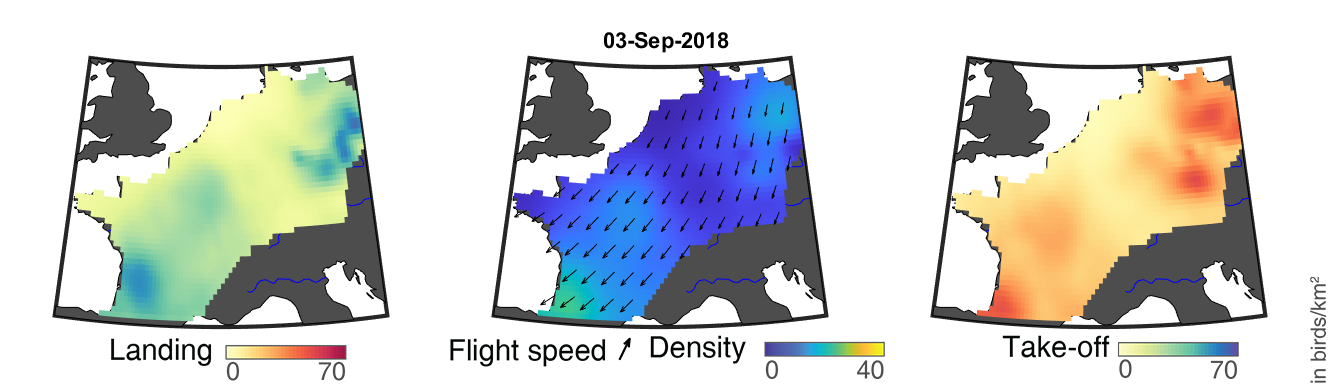

### day_203.png

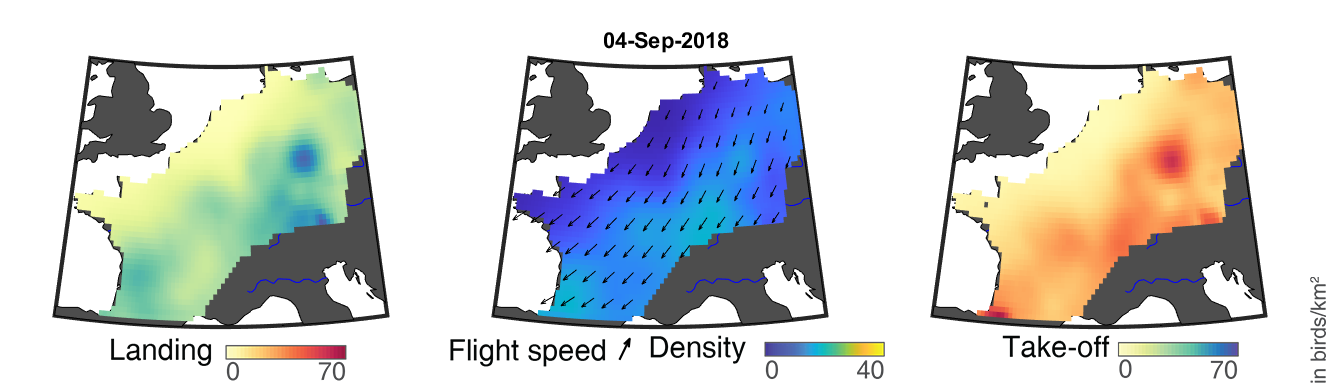

### day_204.png

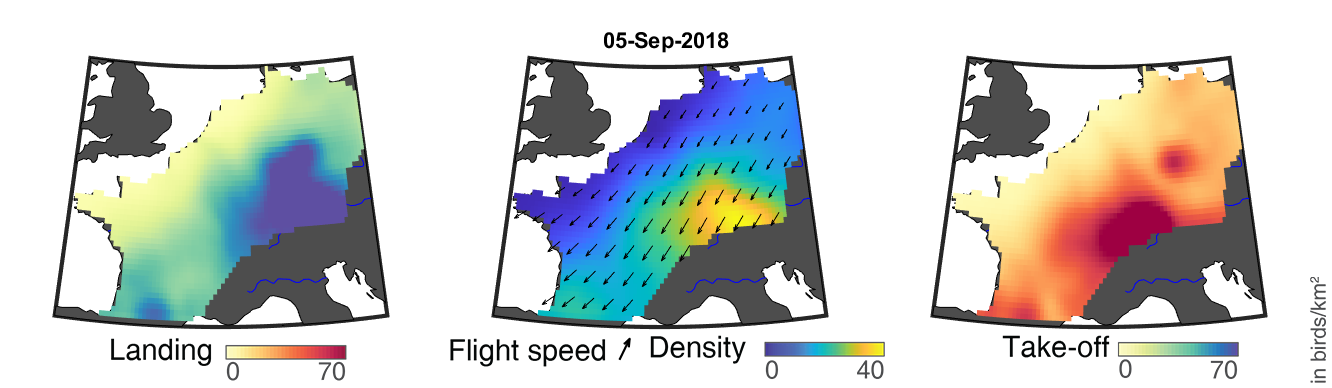

### day_205.png

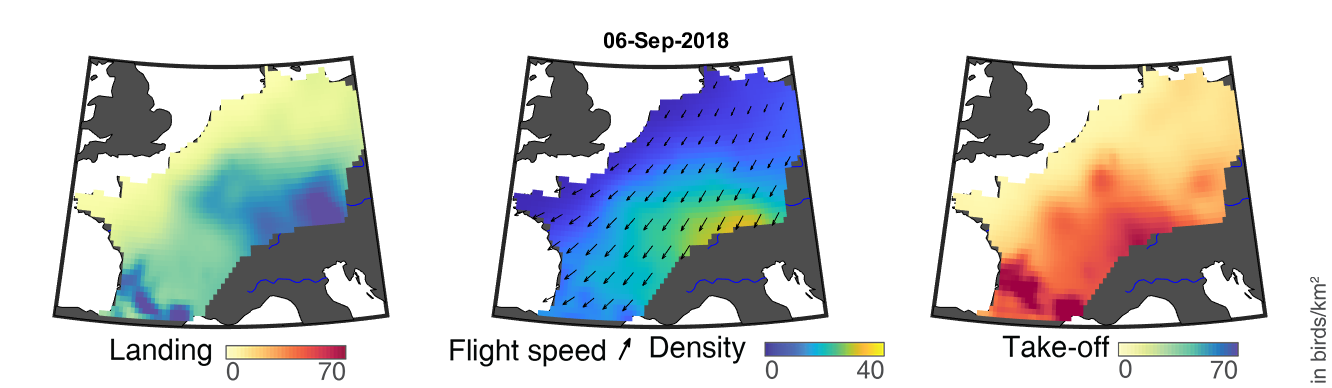

### day_206.png

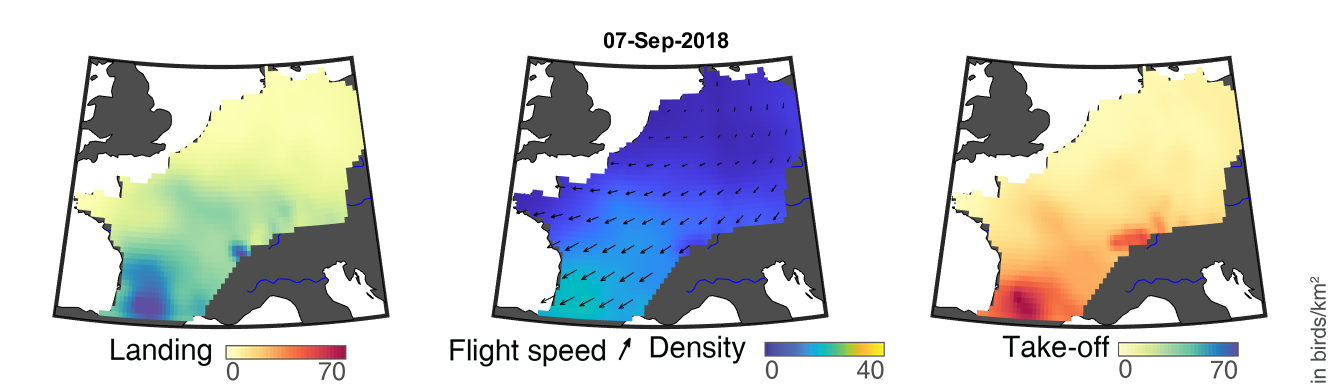

### day_207.png

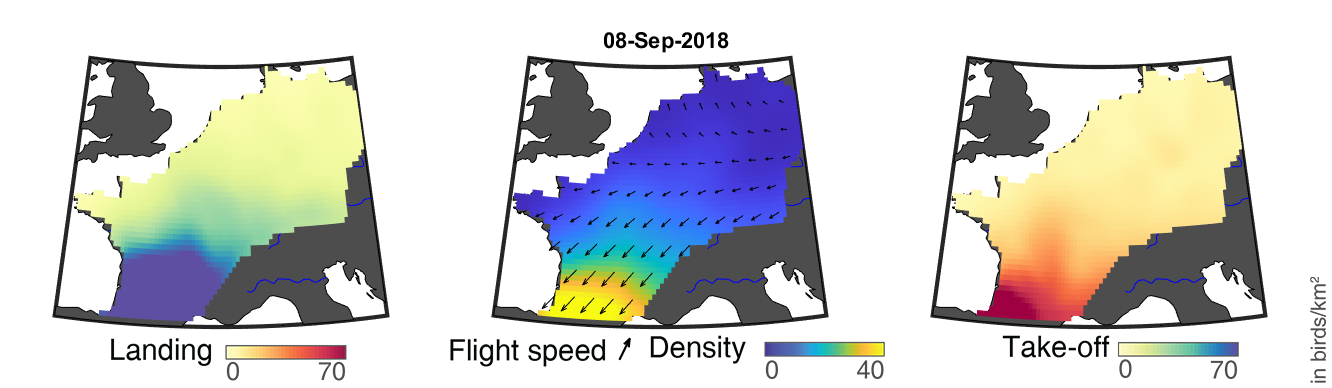

### day_208.png

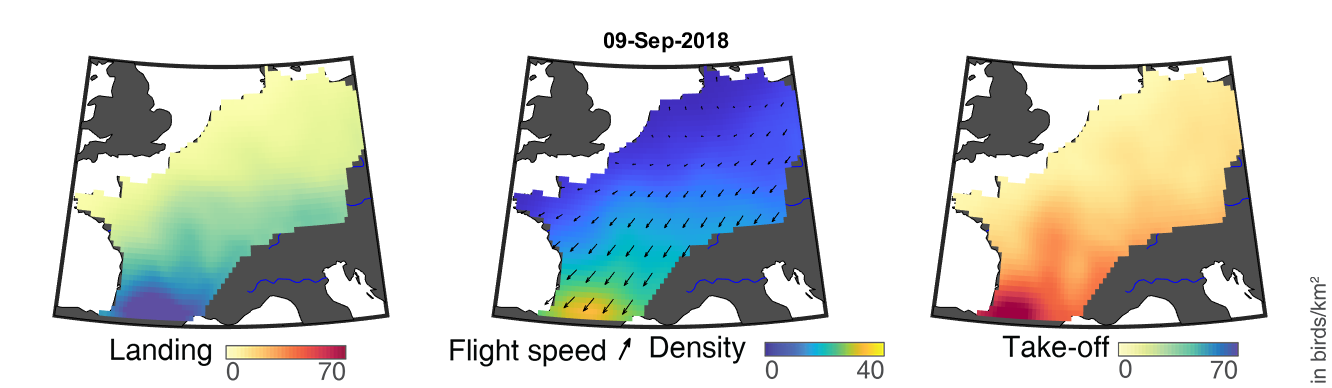

### day_209.png

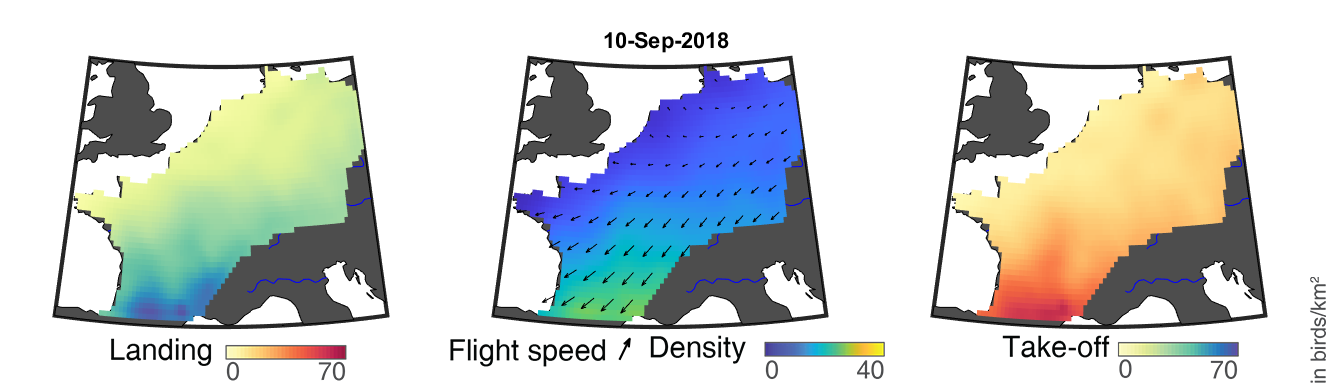

### day_210.png

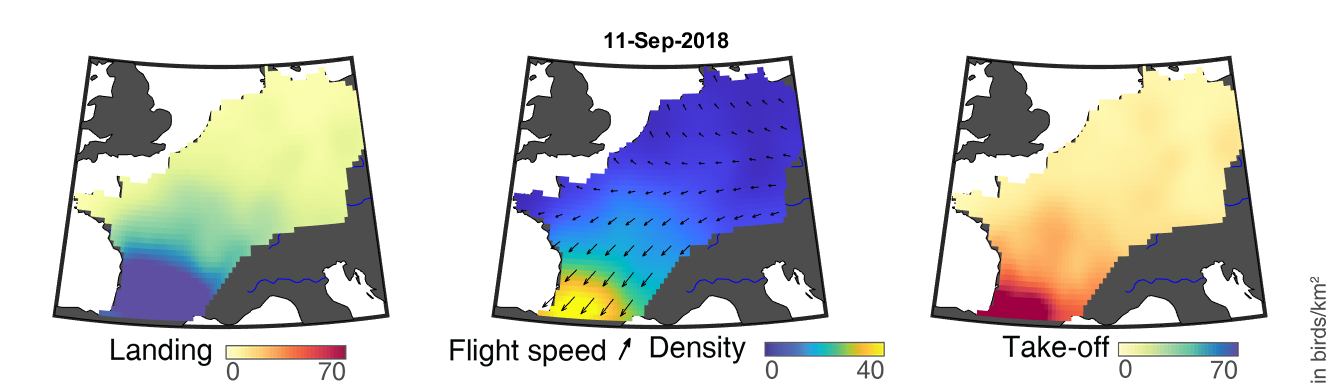

### day_211.png

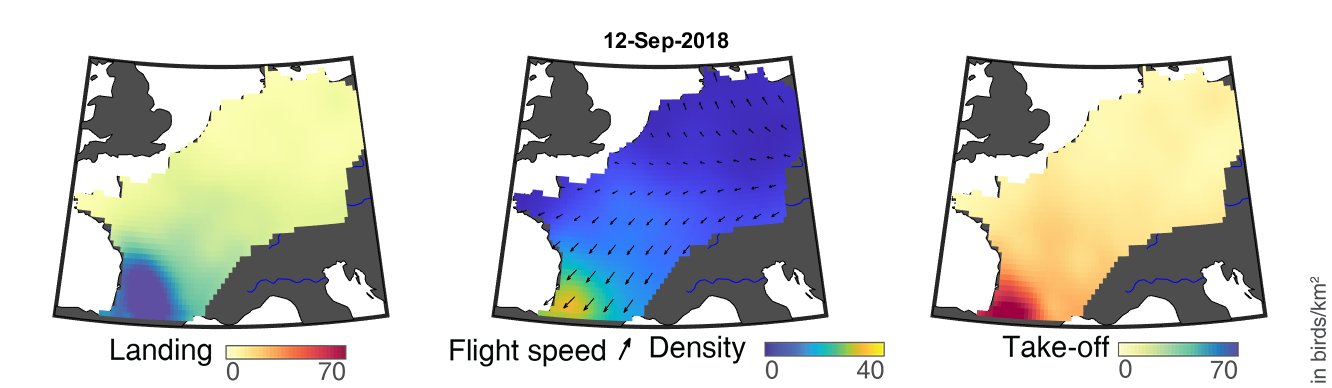

### day_212.png

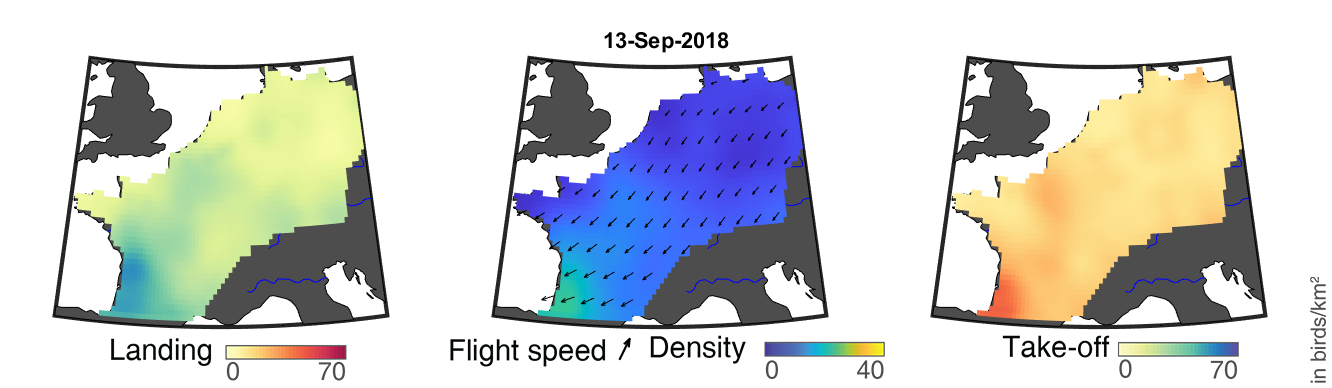

### day_213.png

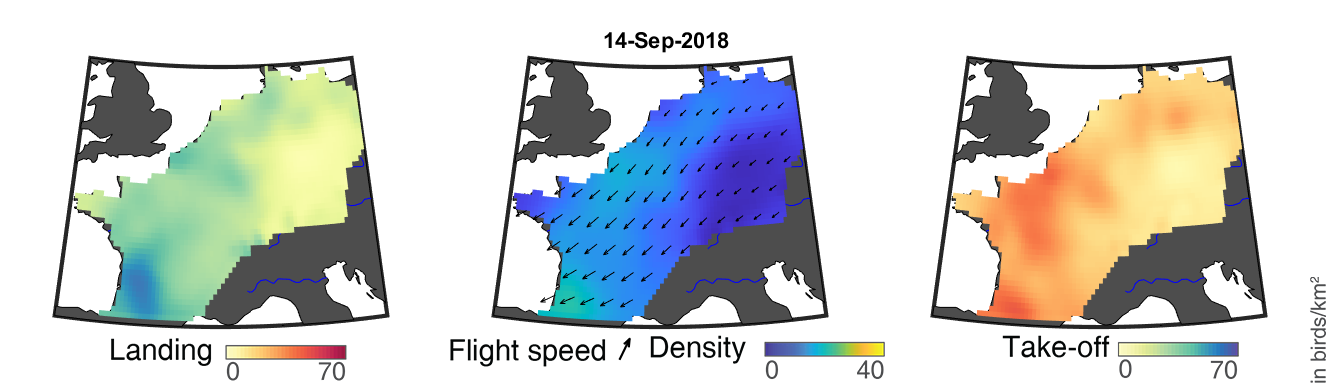

### day_214.png

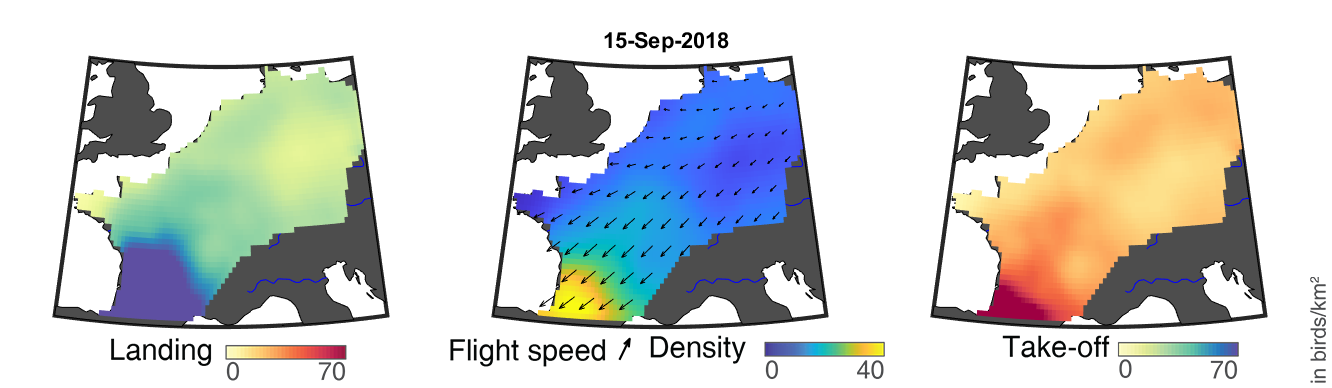

### day_215.png

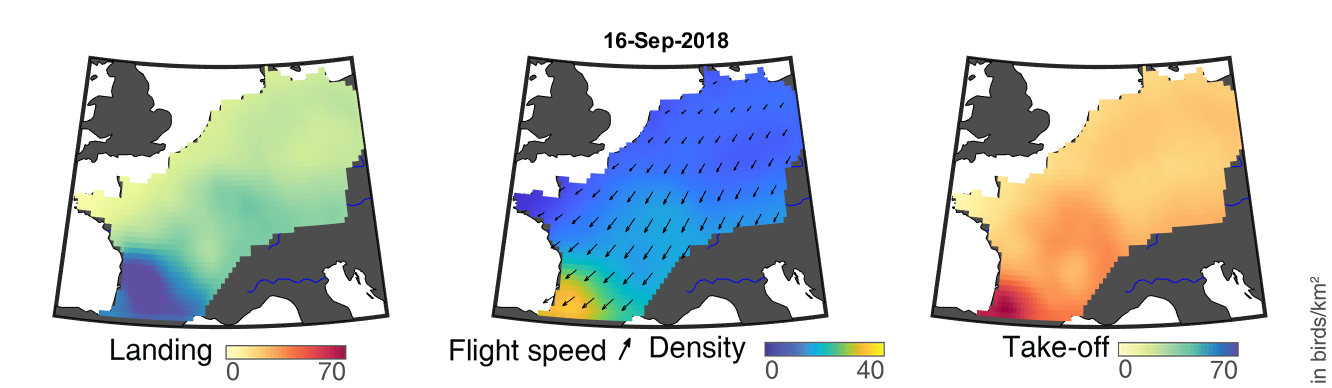

### day_216.png

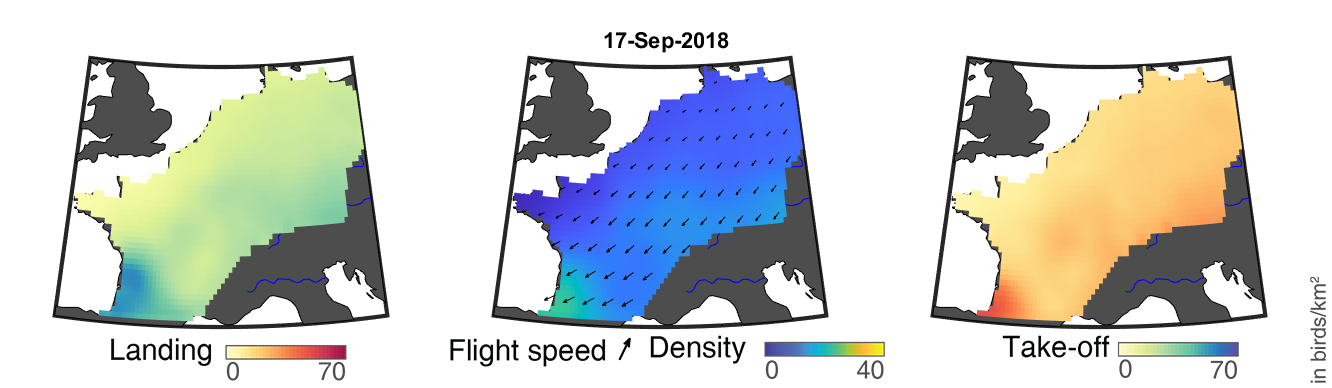

### day_217.png

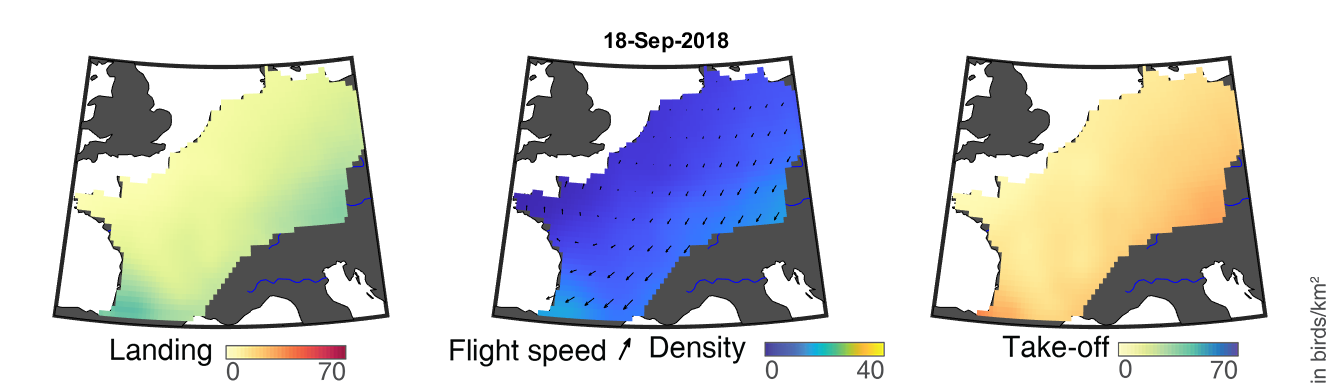

### day_218.png

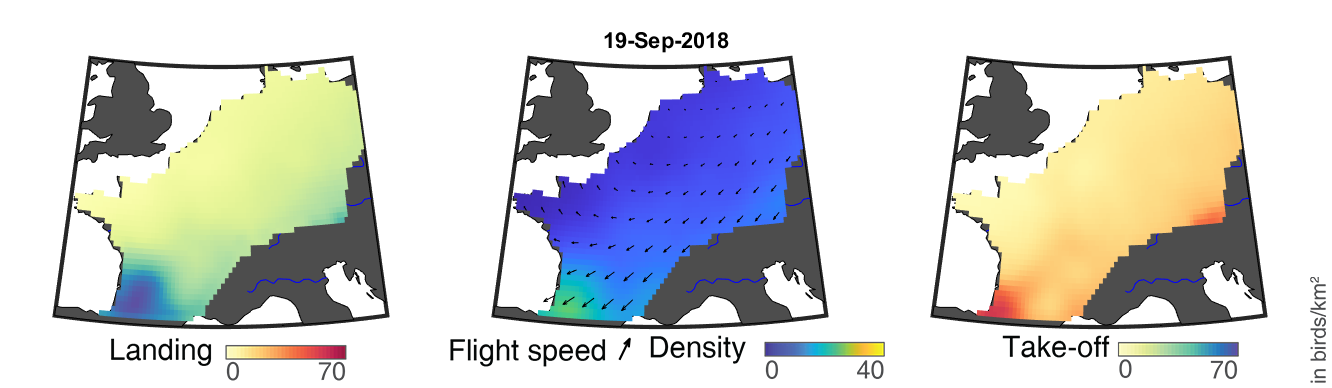

### day_219.png

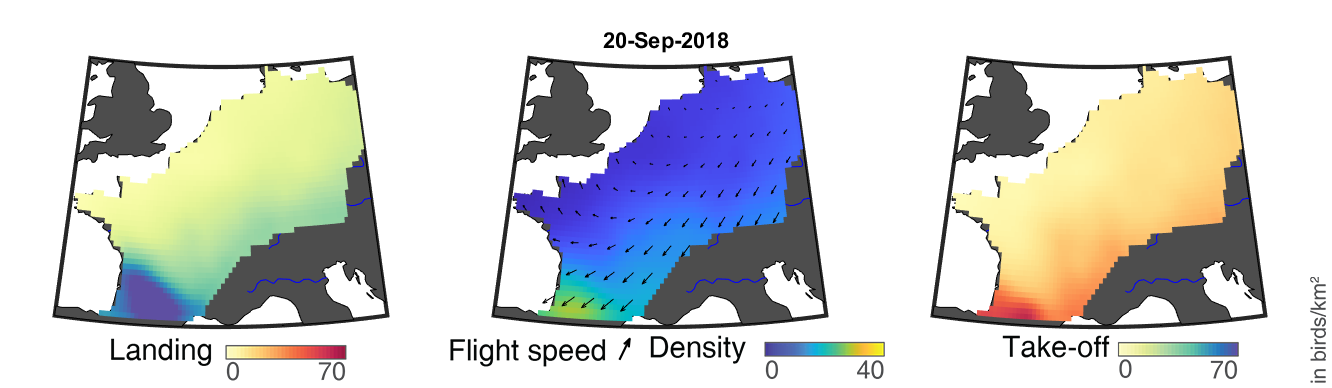

### day_220.png

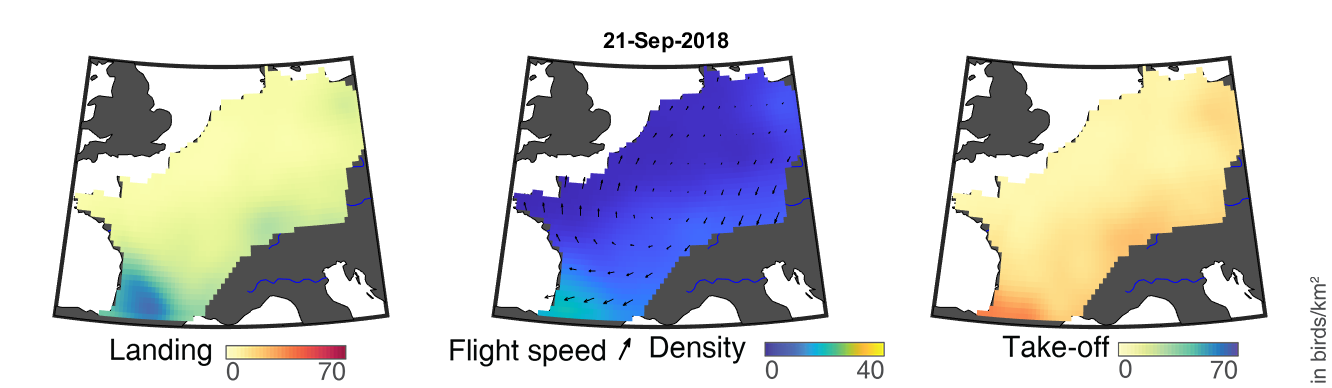

### day_221.png

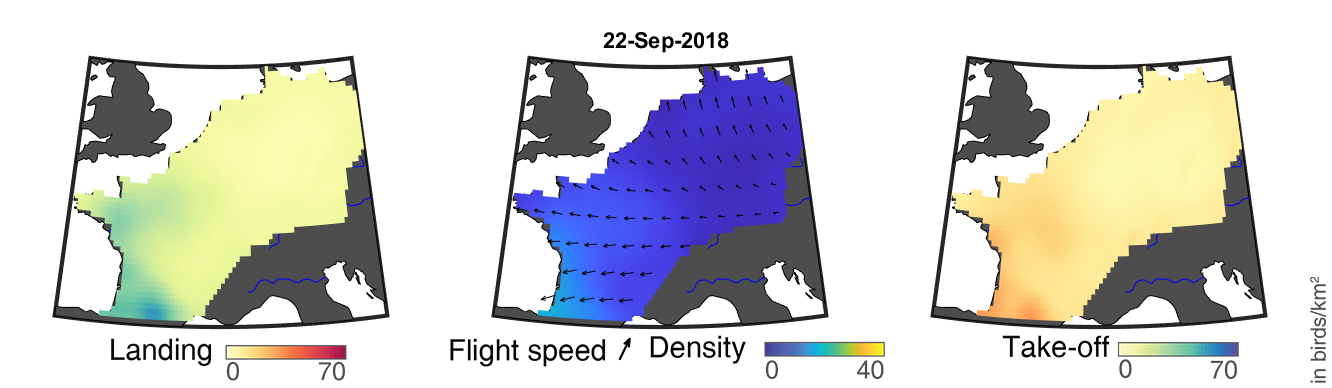

### day_222.png

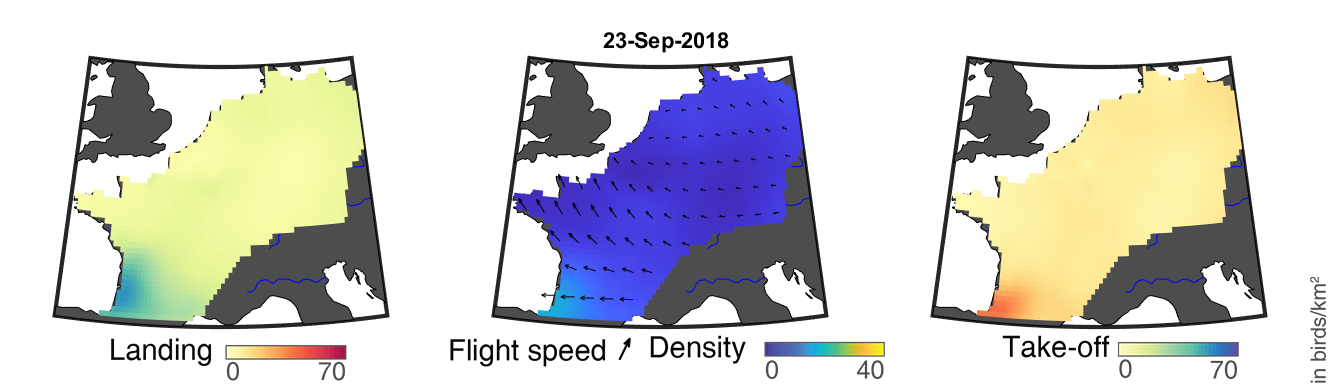

### day_223.png

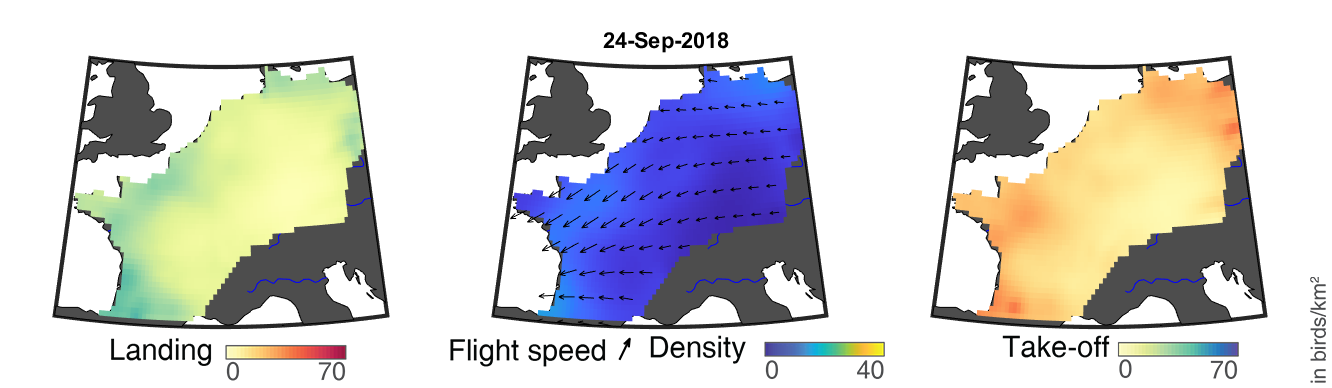

### day_224.png

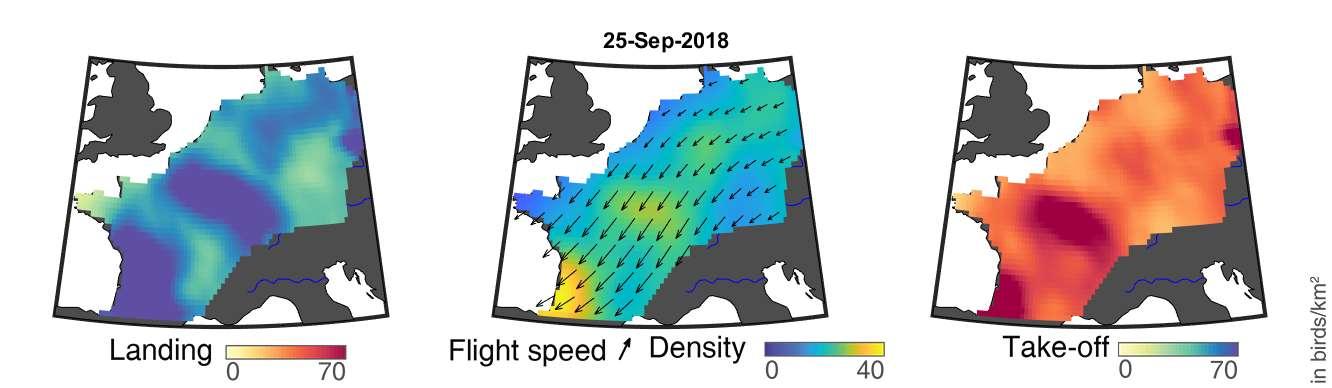

### day_225.png

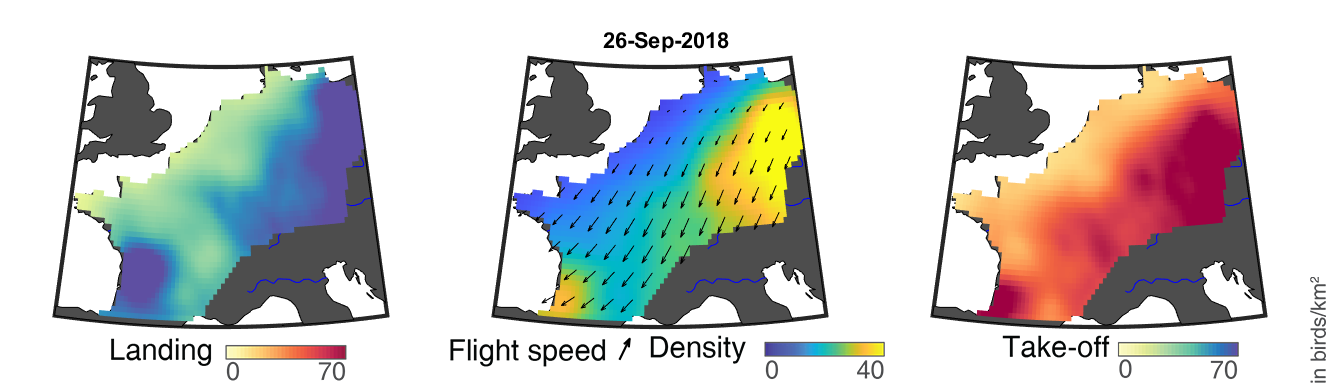

### day_226.png

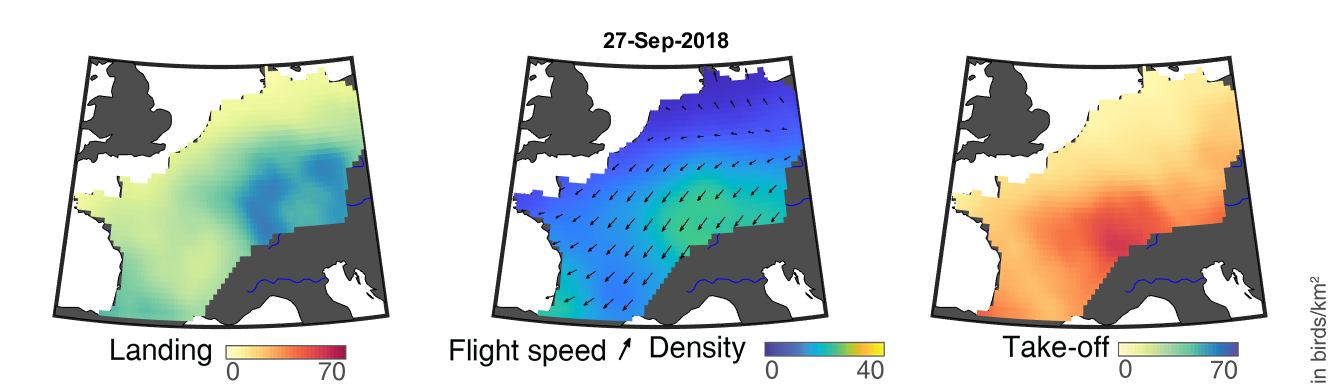

### day_227.png

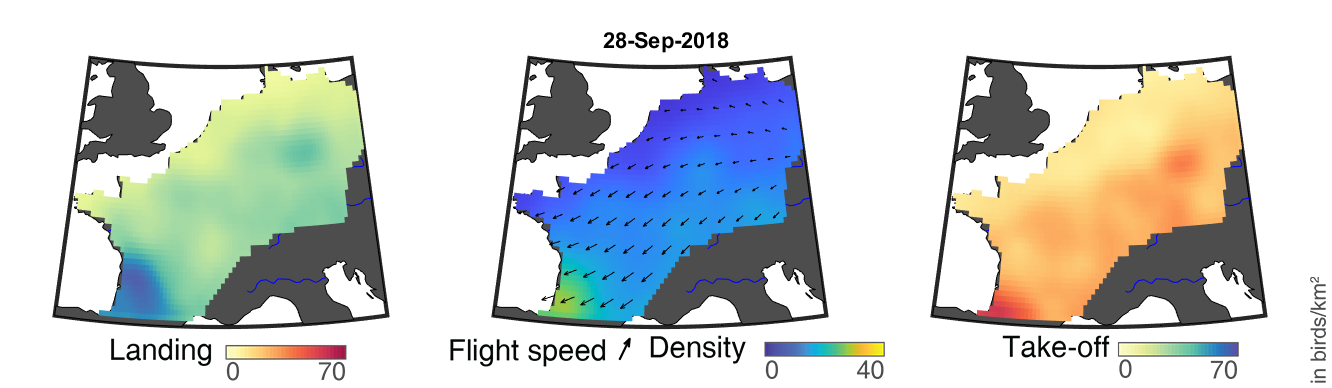

### day_228.png

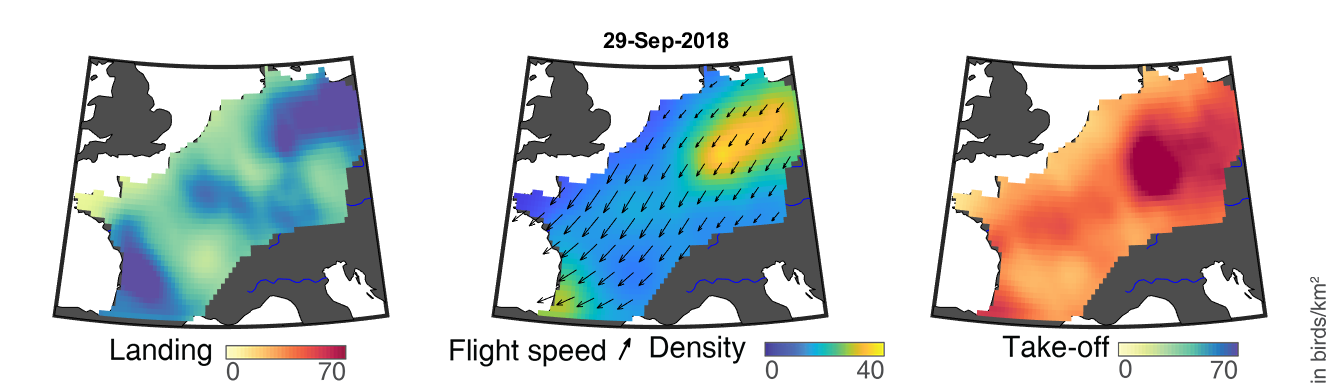

### day_229.png

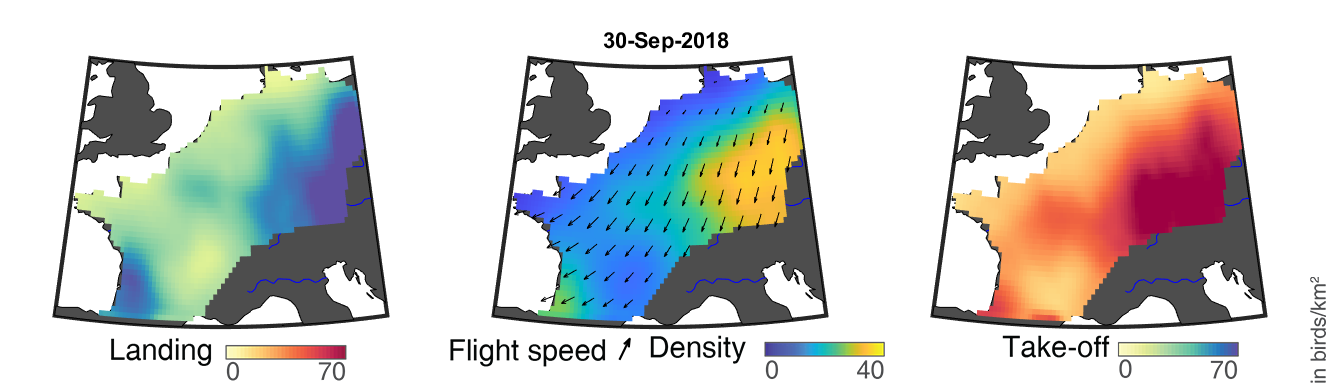

### day_230.png

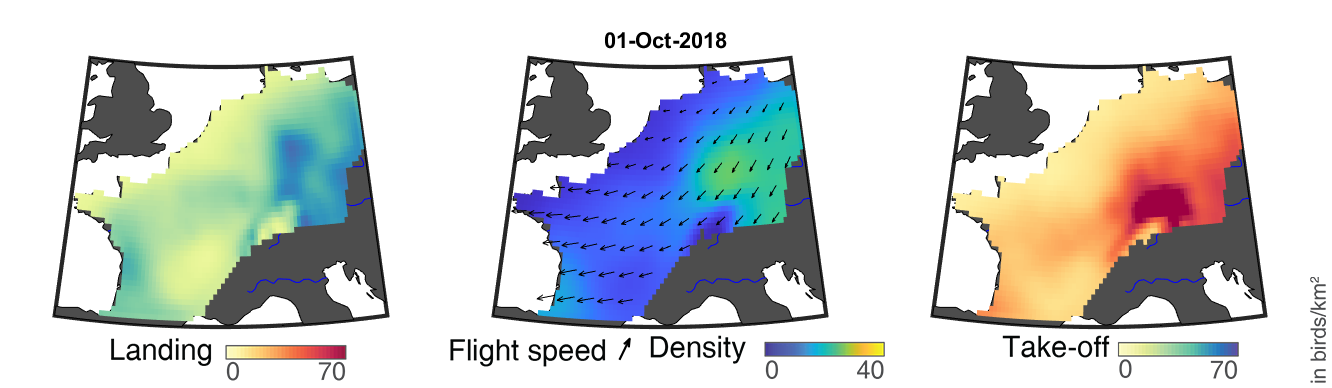

### day_231.png

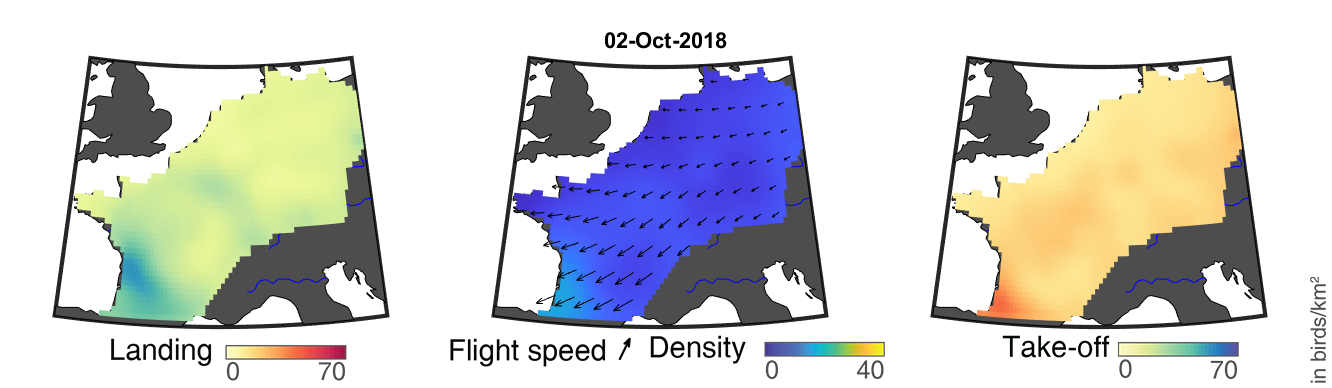

### day_232.png

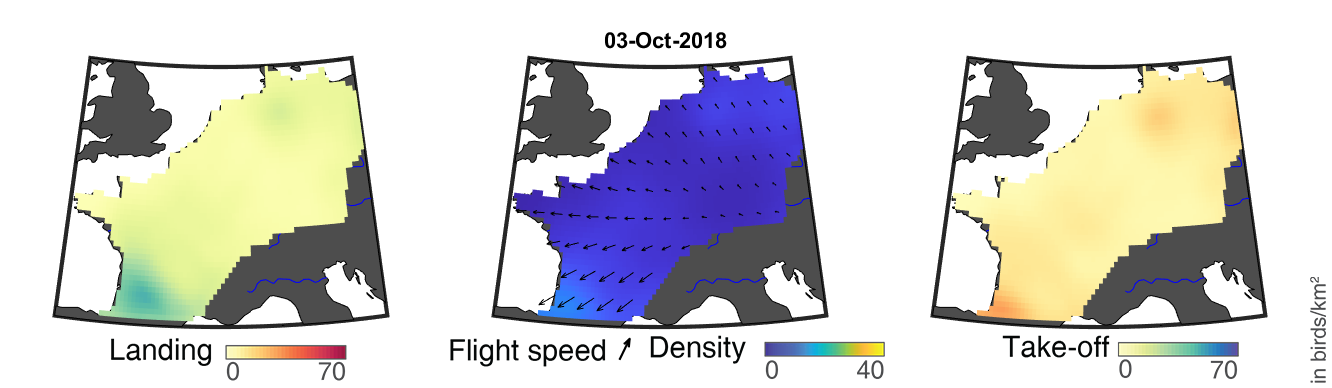

### day_233.png

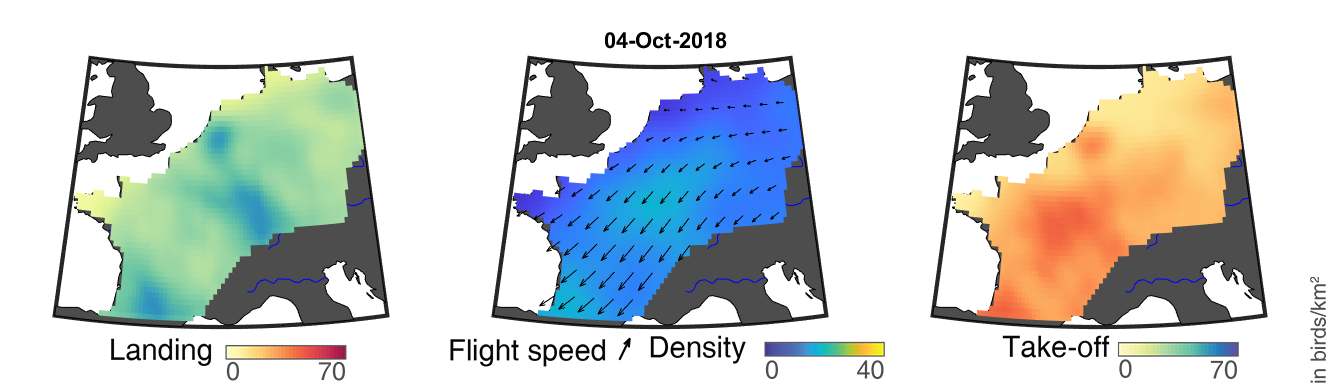

### day_234.png

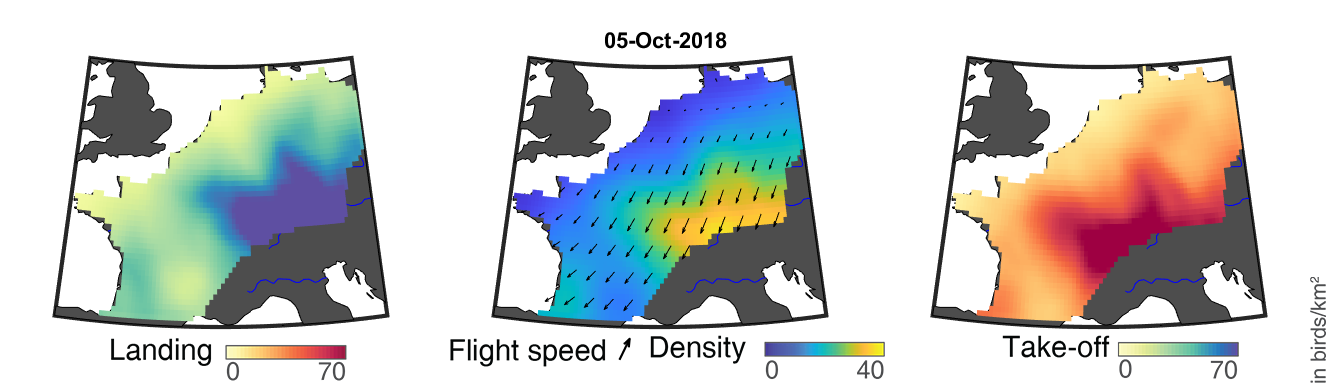

### day_235.png

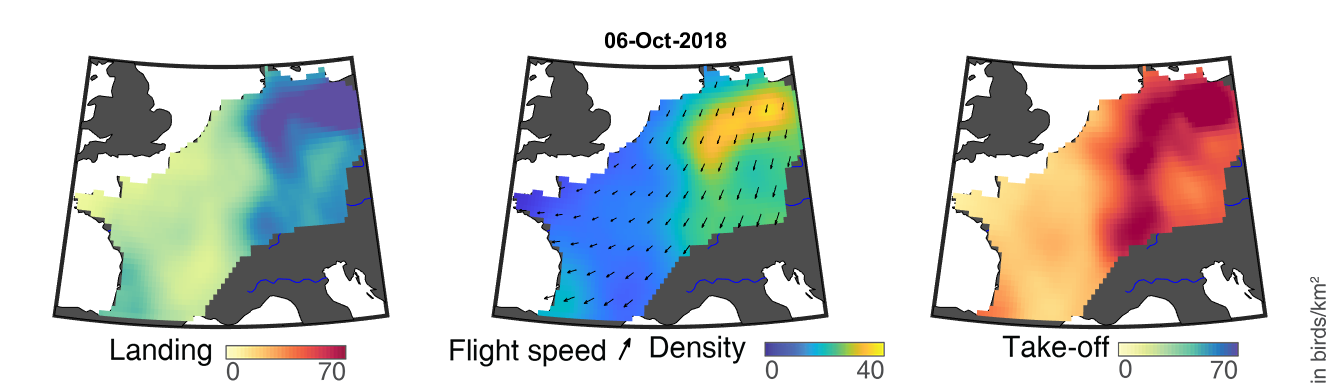

### day_236.png

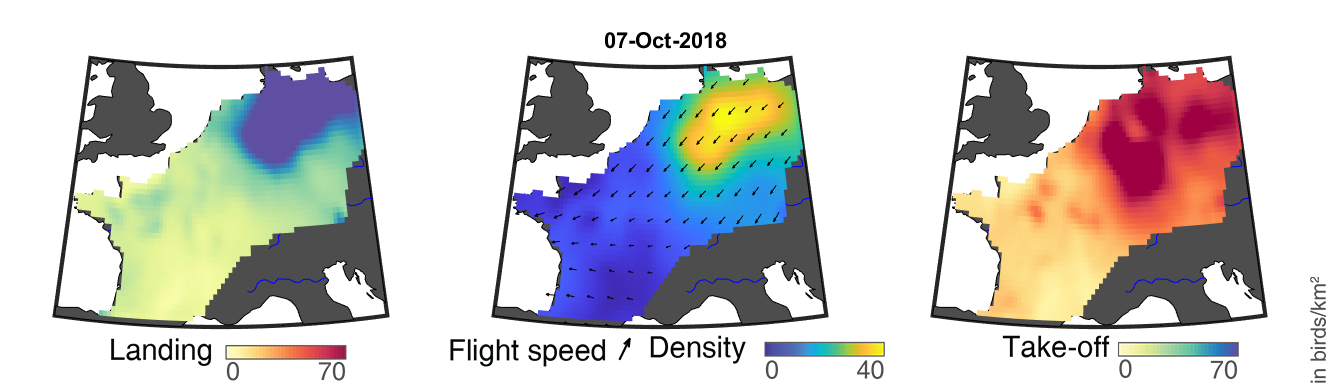

### day_237.png

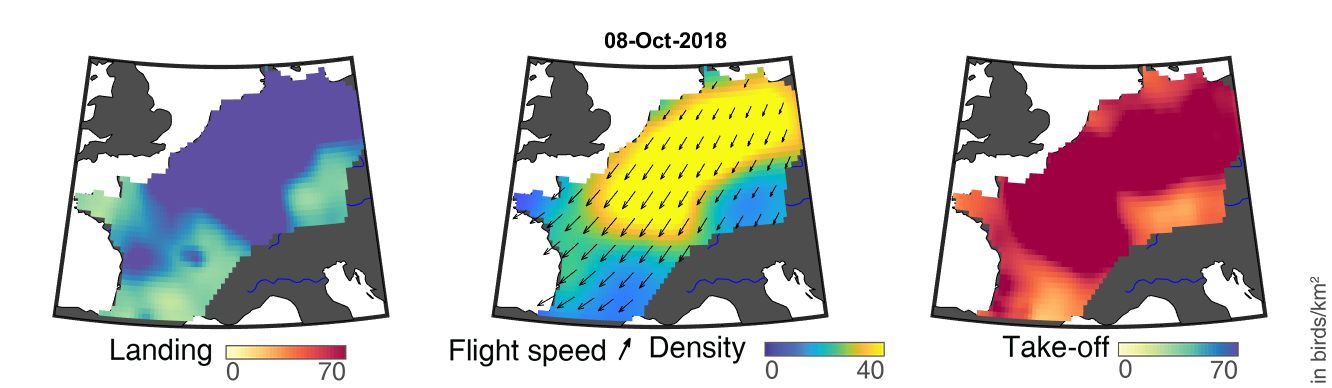

### day_238.png

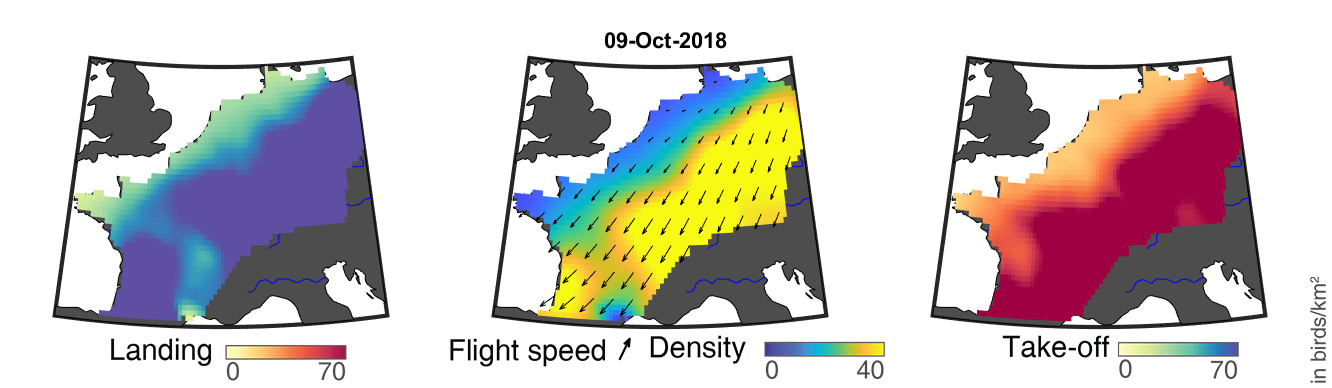

### day_239.png

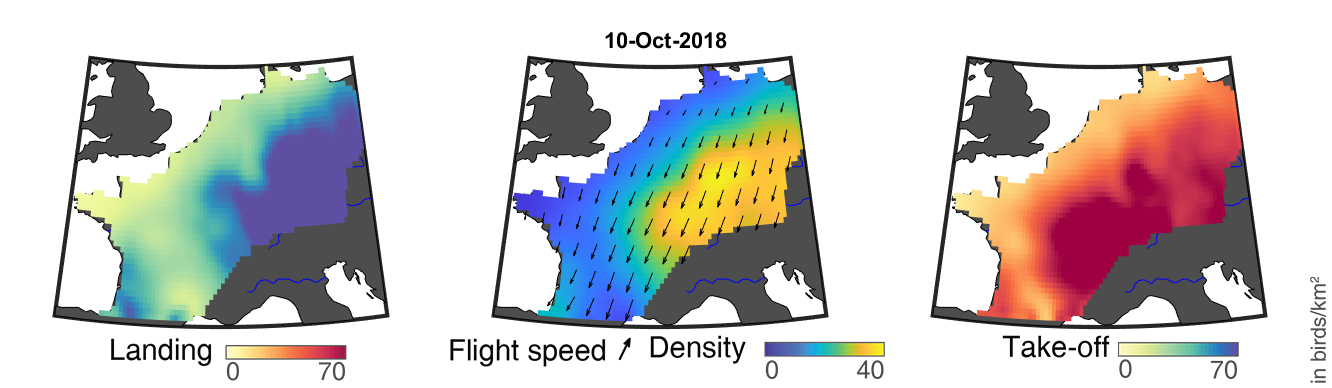

### day_240.png

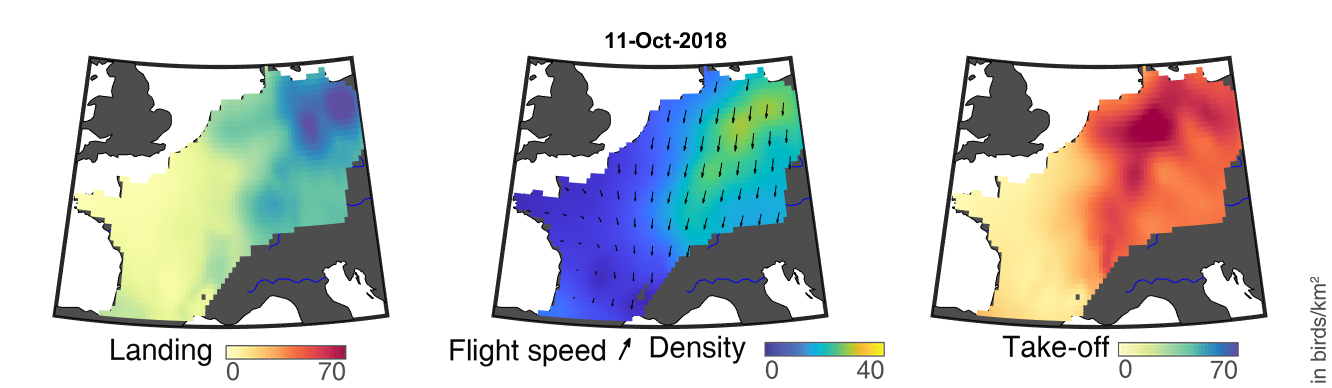

### day_241.png

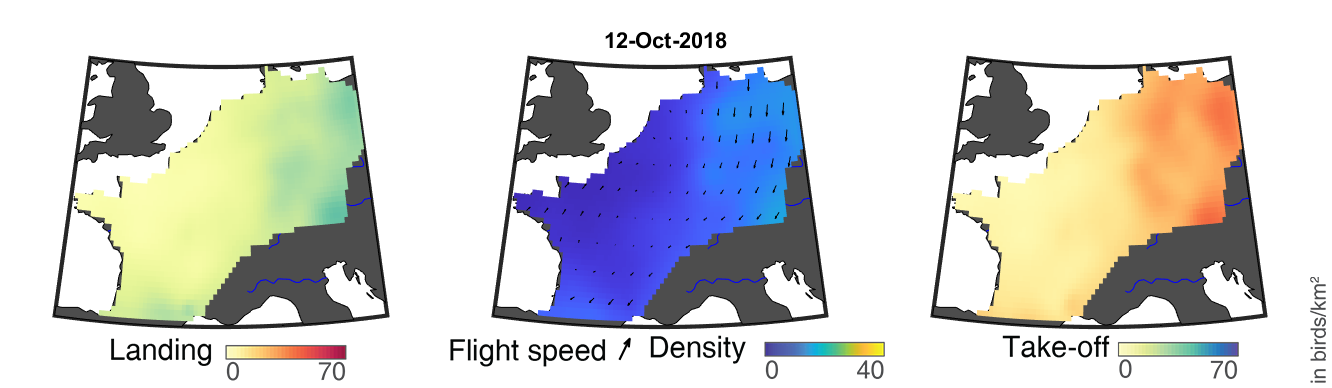

### day_242.png

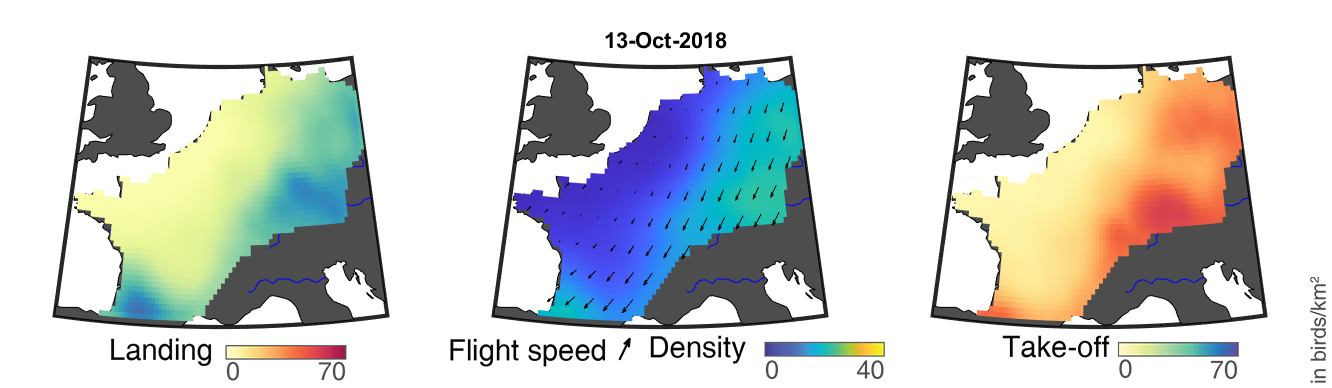

### day_243.png

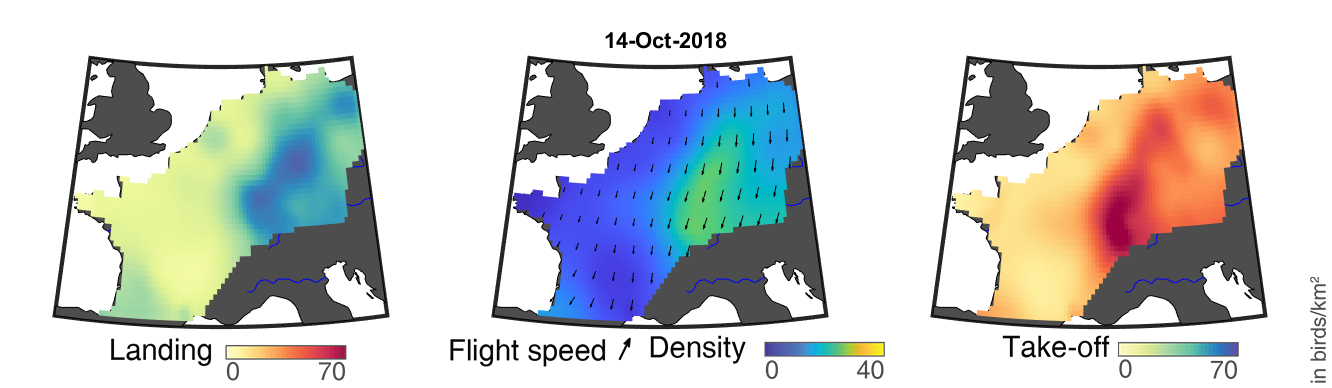

### day_244.png

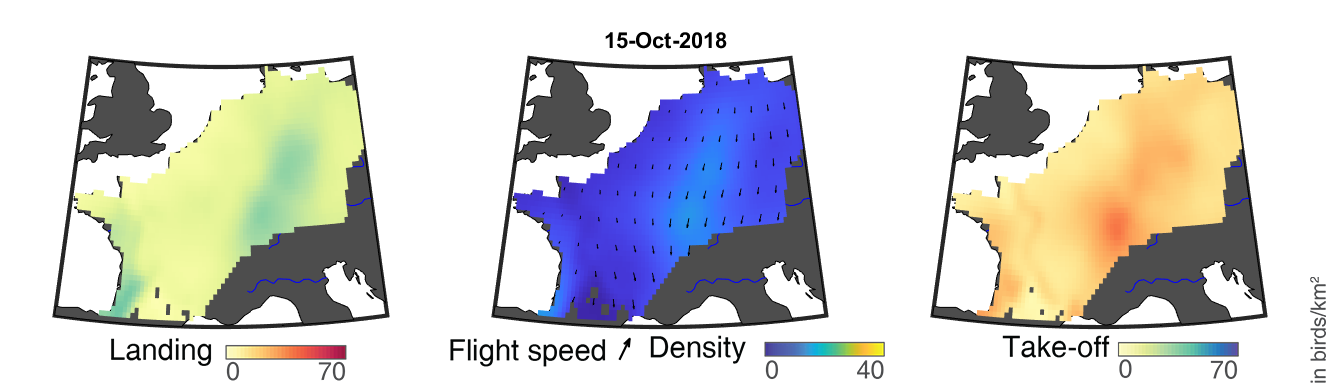

### day_245.png

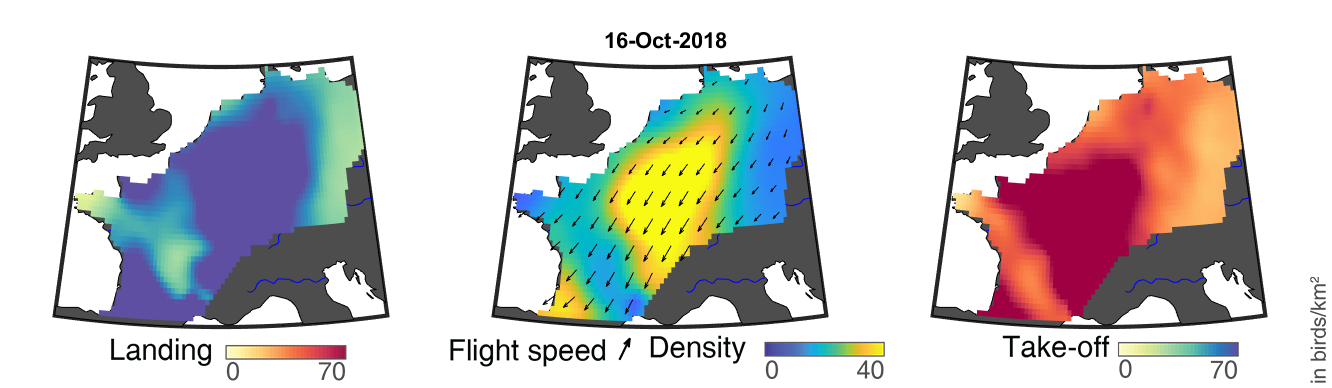

### day_246.png

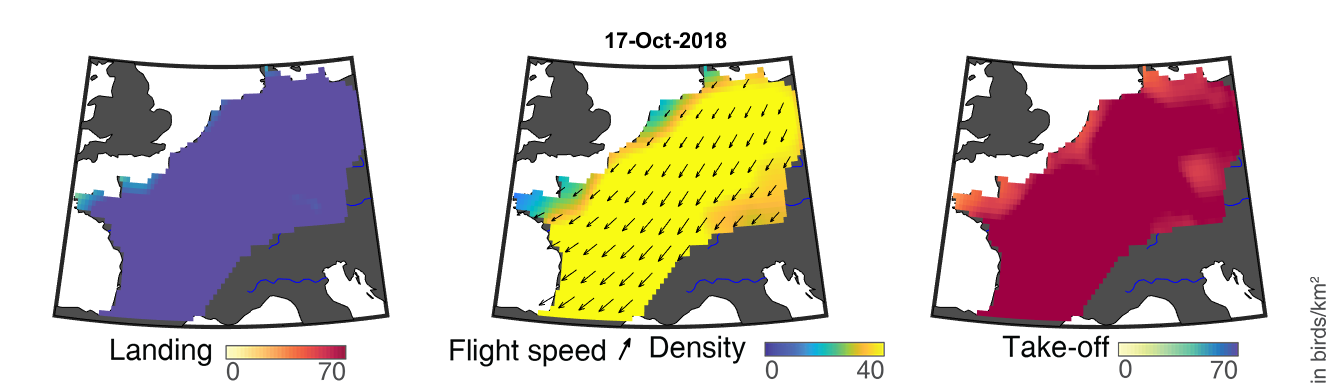

### day_247.png

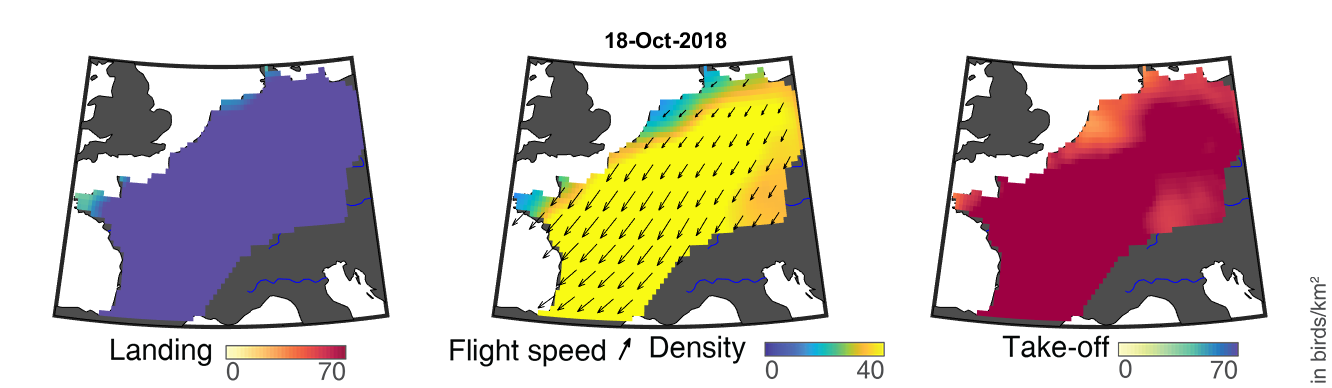

### day_248.png

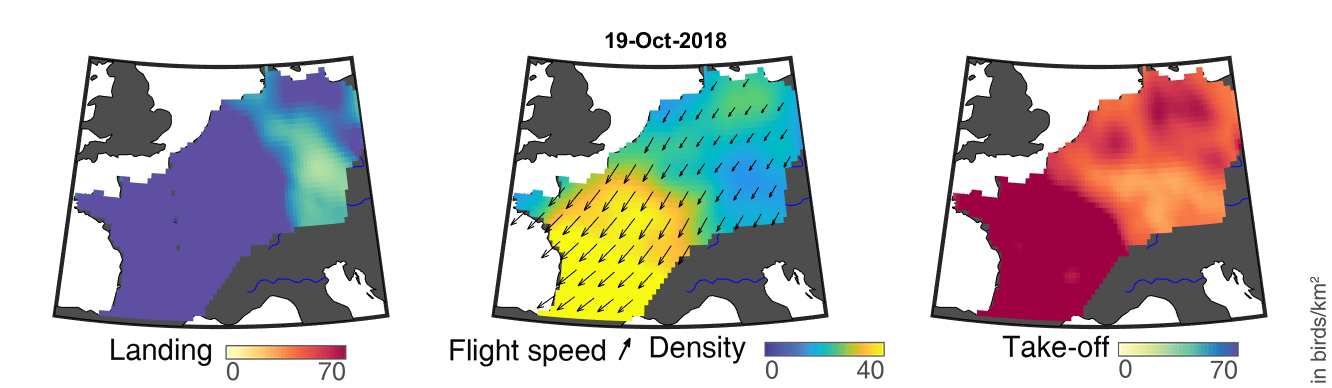

### day_249.png

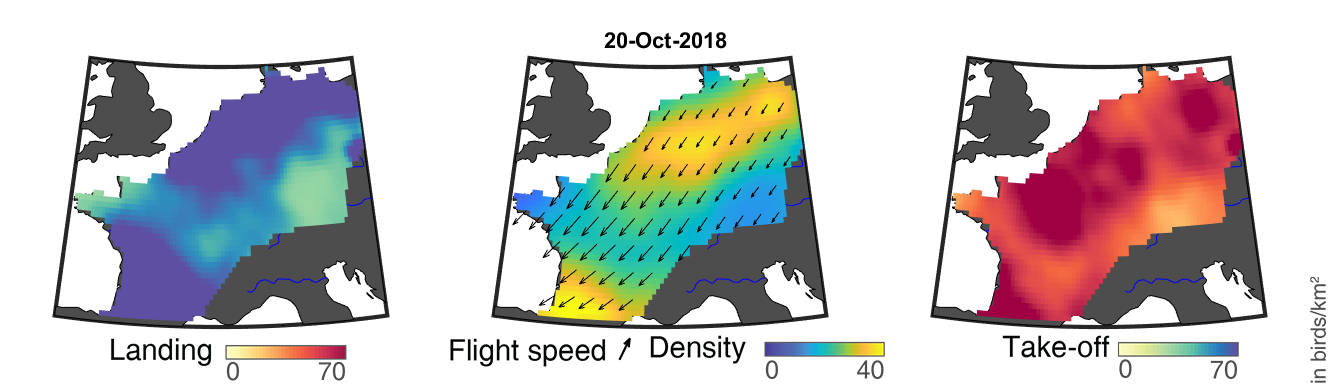

### day_250.png

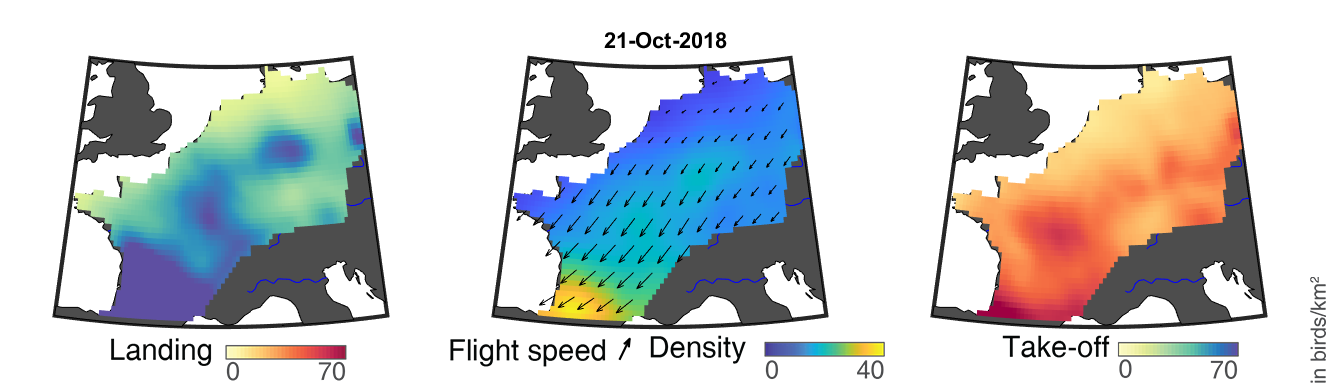

### day_251.png

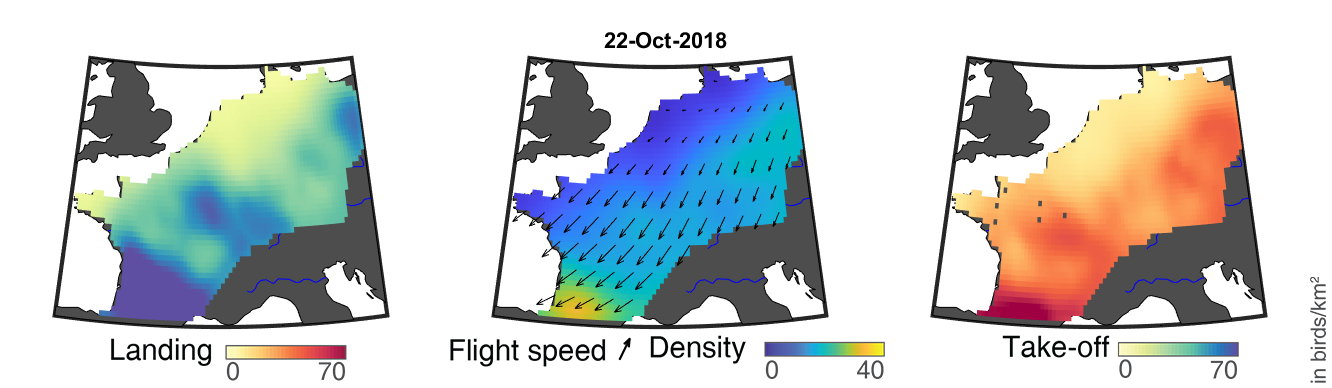

### day_252.png

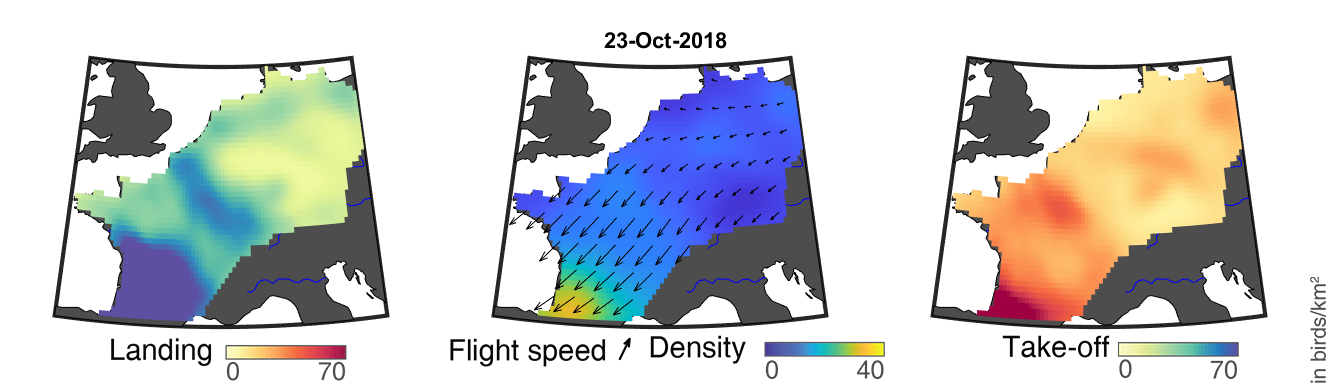

### day_253.png

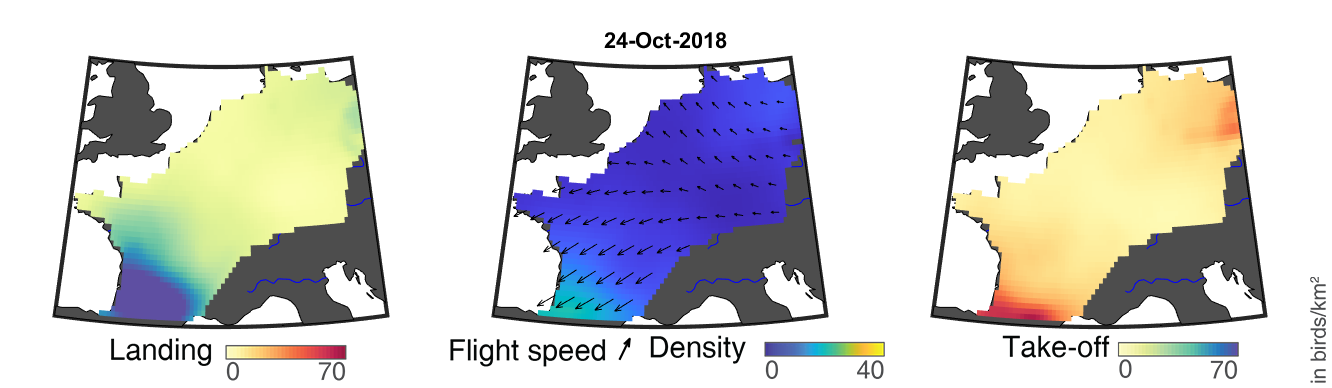

### day_254.png

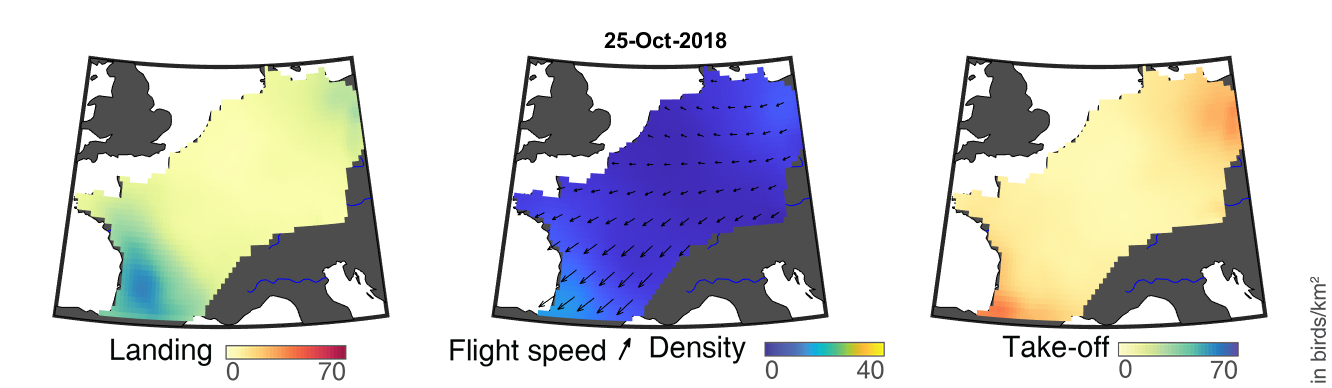

### day_255.png

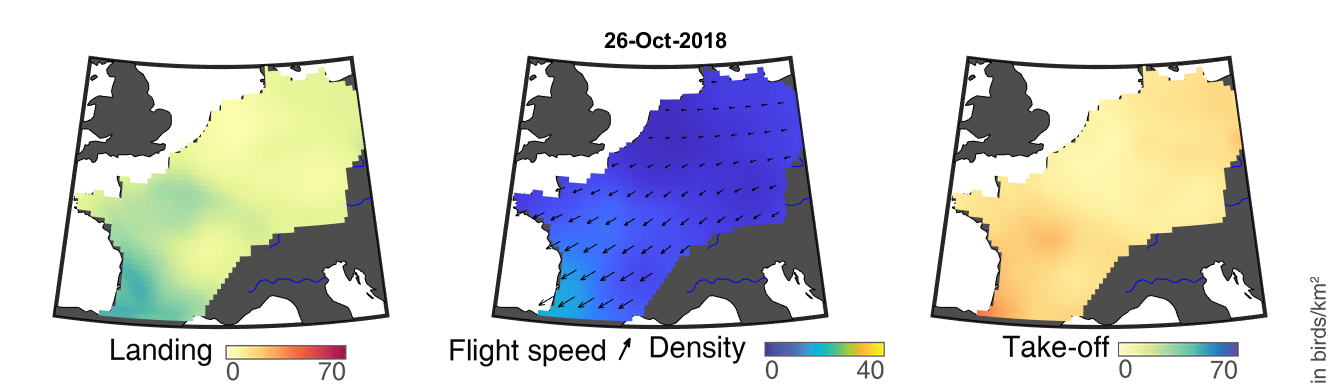

### day_256.png

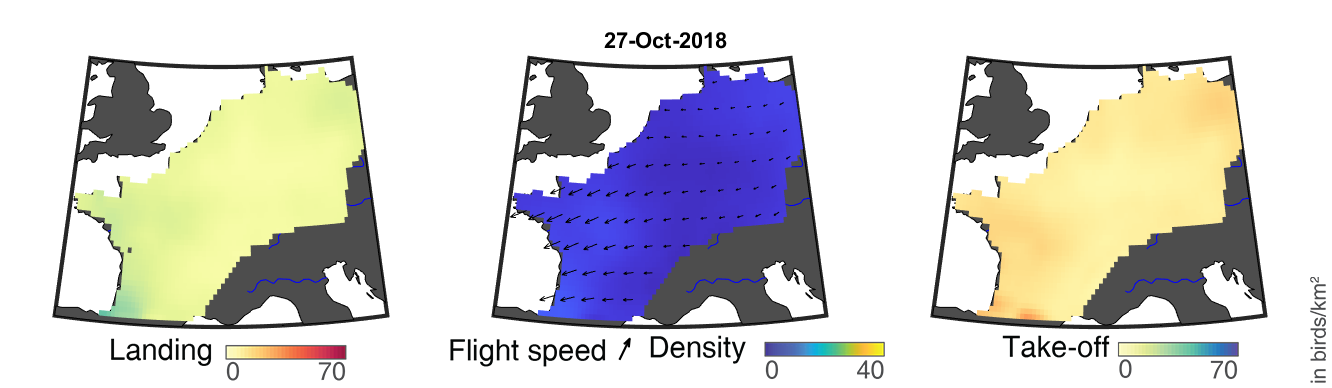

### day_257.png

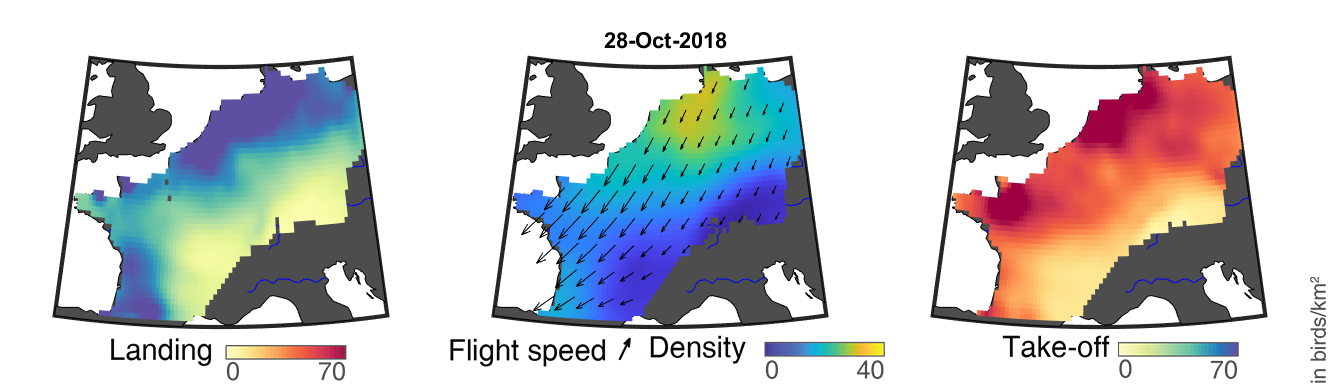

### day_258.png

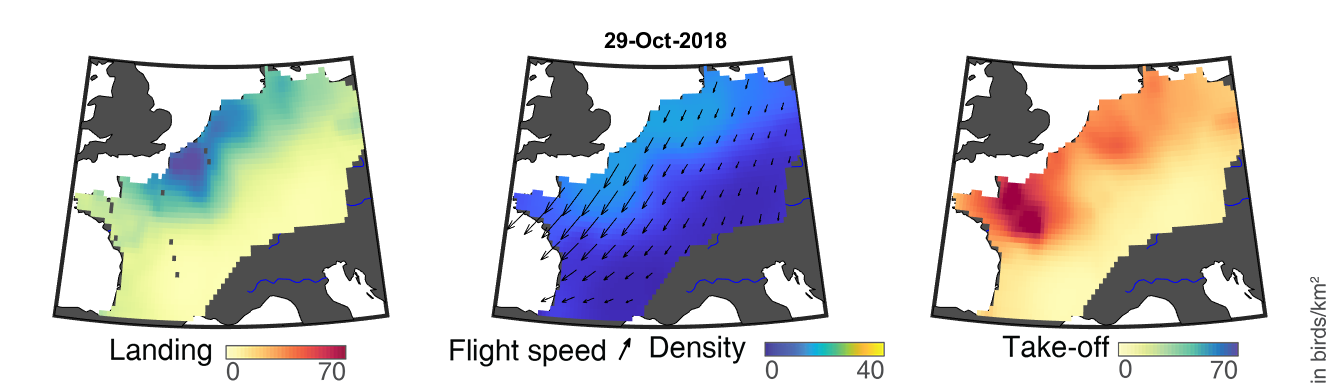

### day_259.png

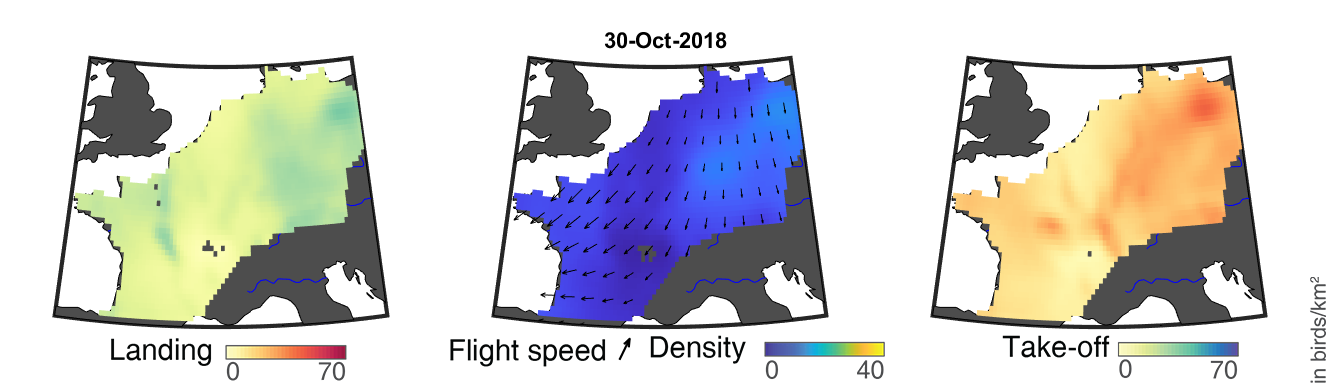

### day_260.png

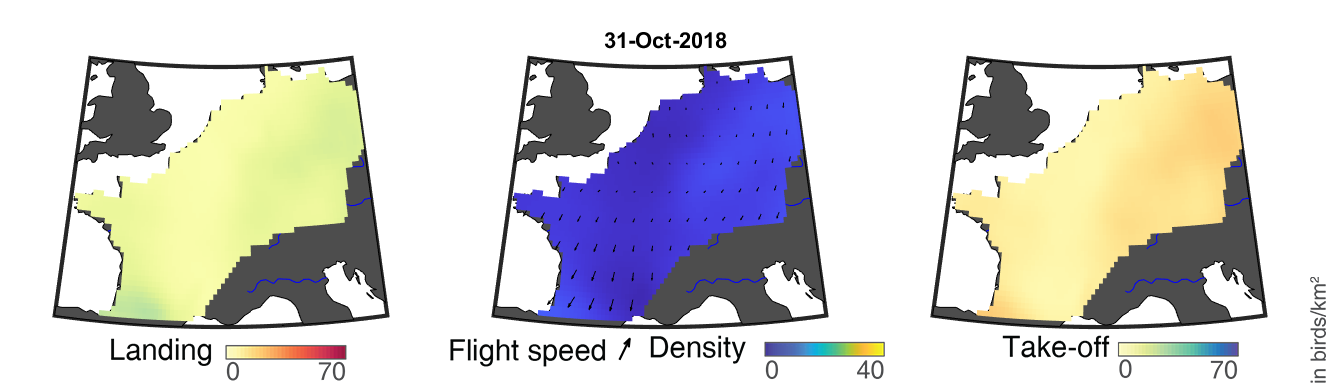

### day_261.png

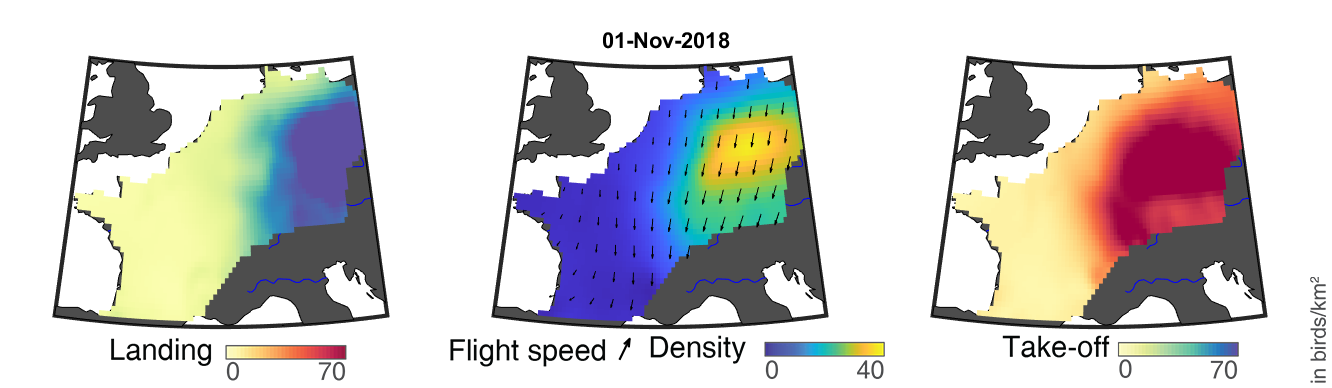

### day_262.png

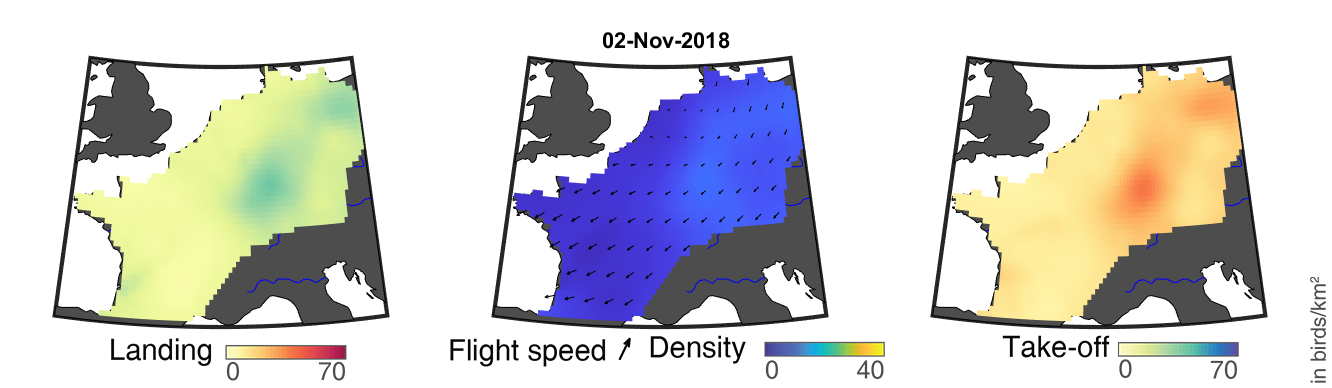

### day_263.png

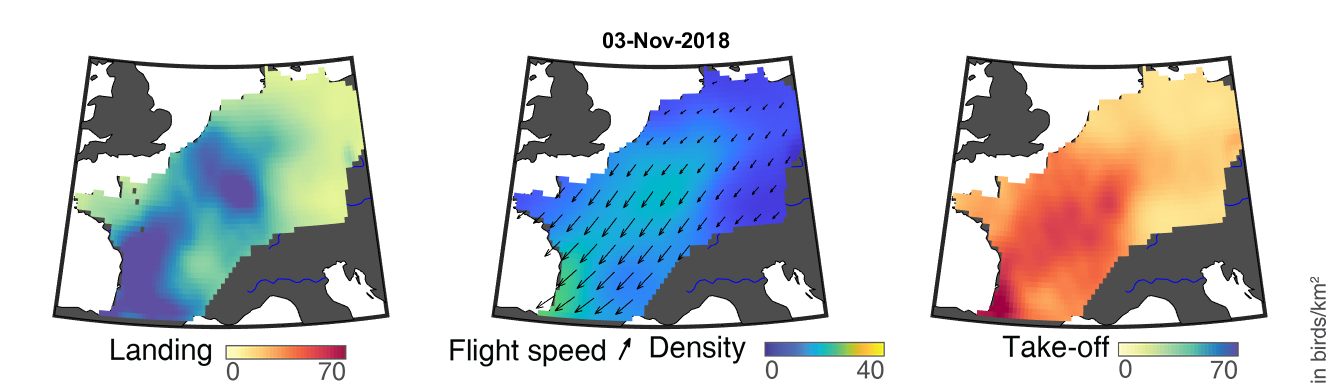

### day_264.png

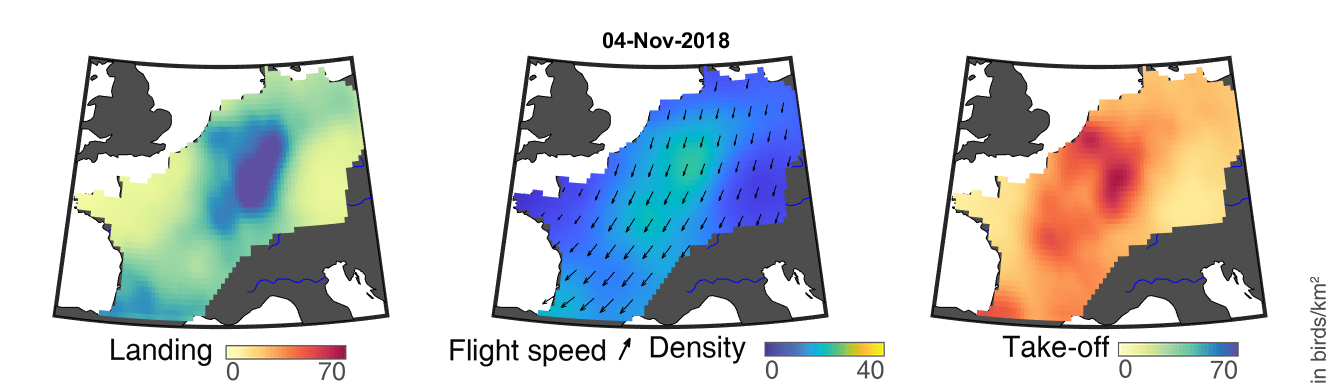

### day_265.png

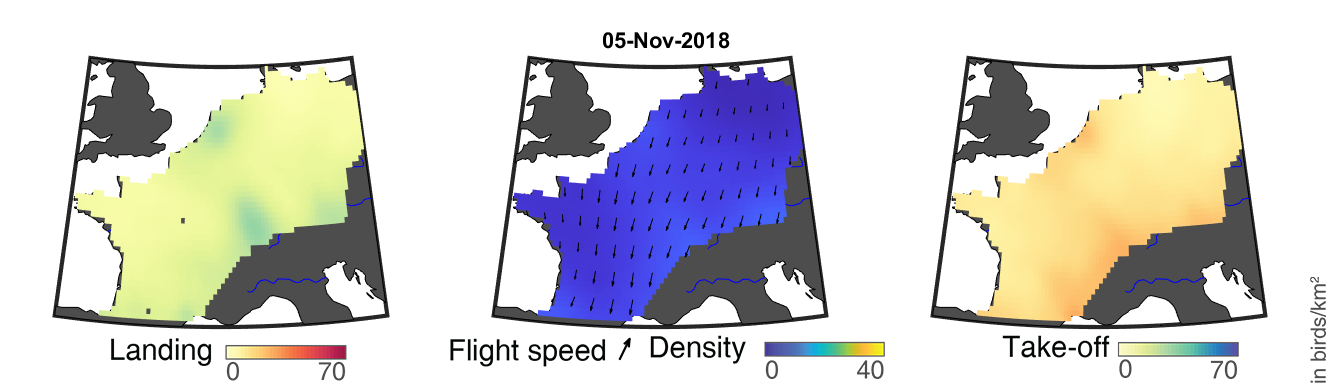

### day_266.png

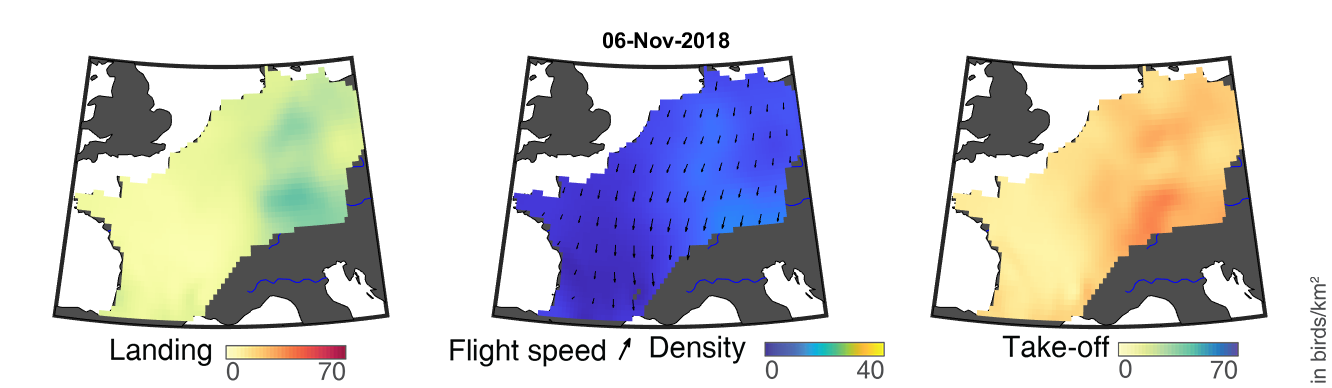

### day_267.png

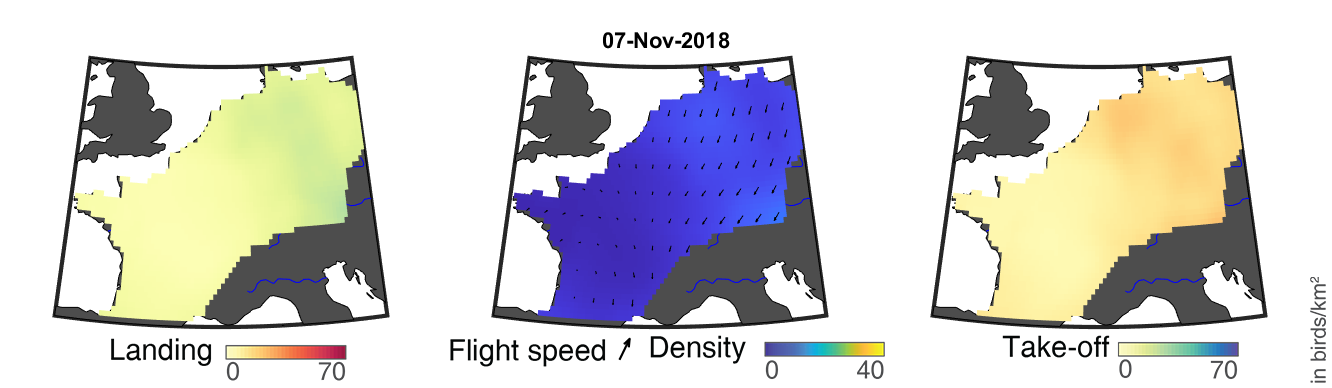

### day_268.png

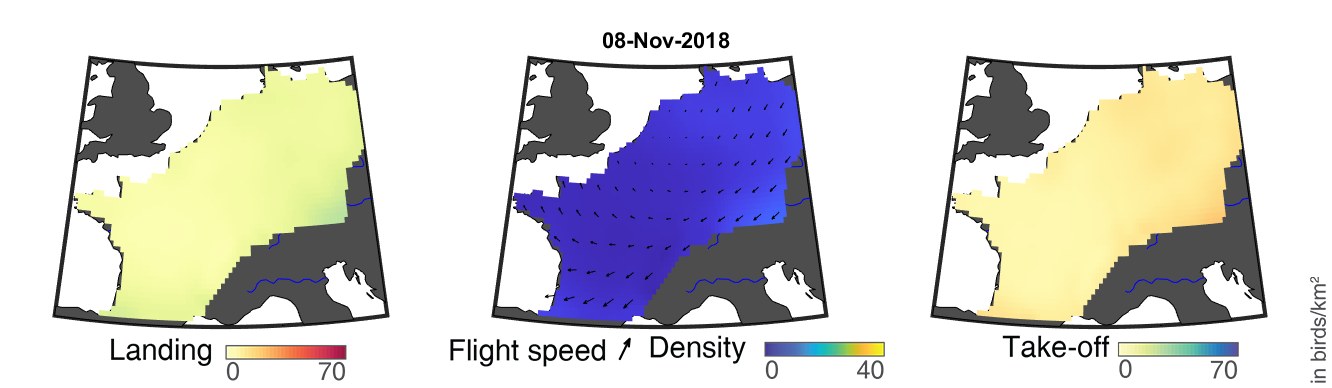

### day_269.png

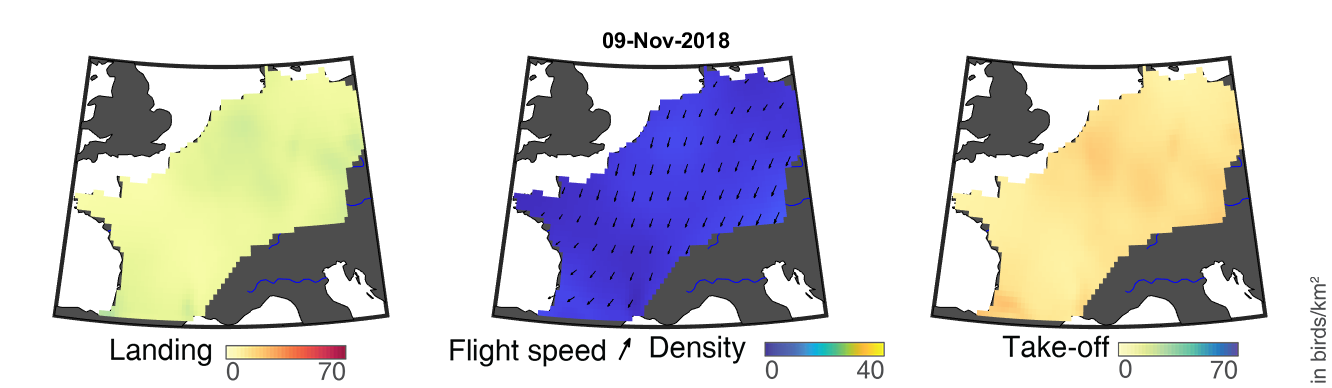

### day_270.png

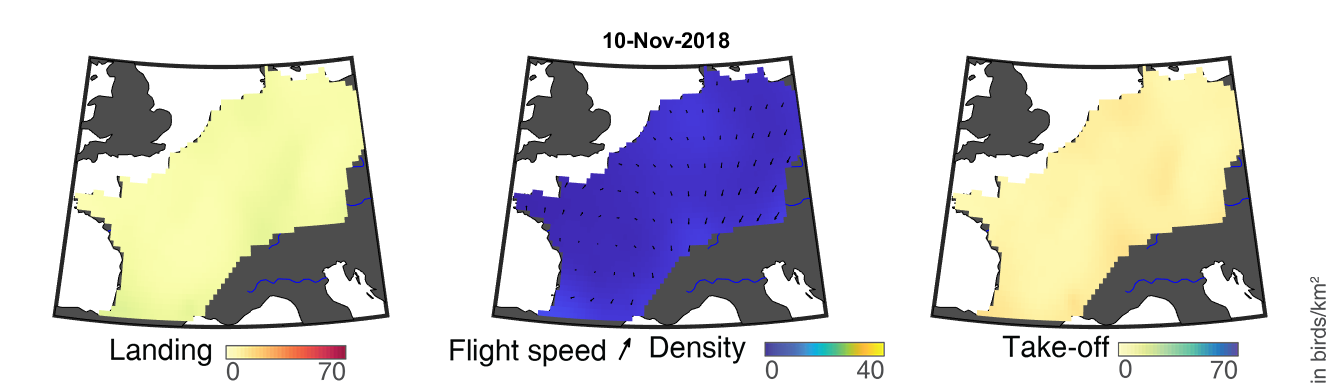

### day_271.png

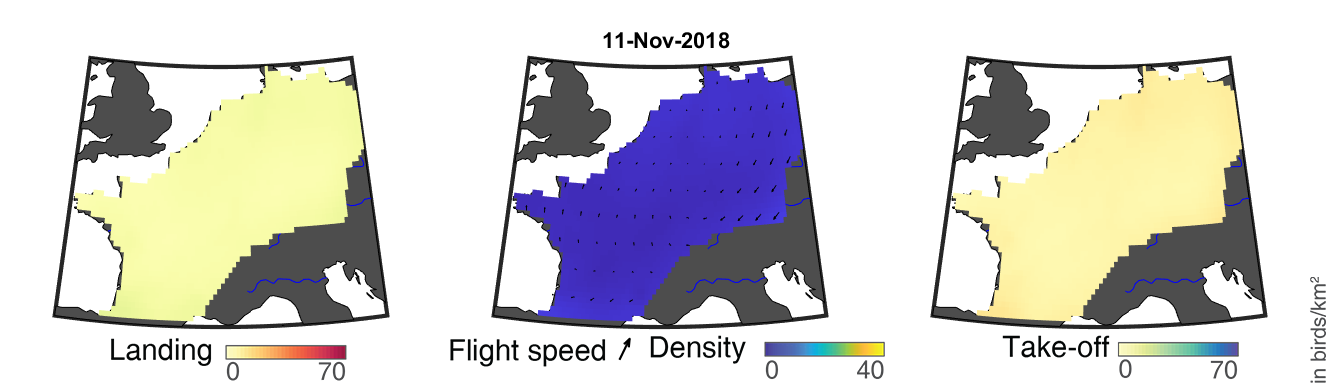

### day_272.png

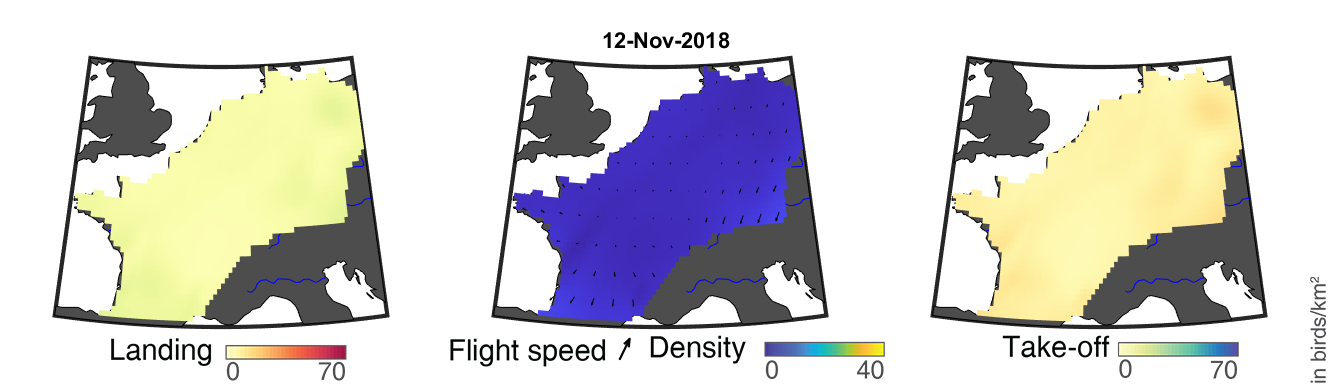

### day_273.png

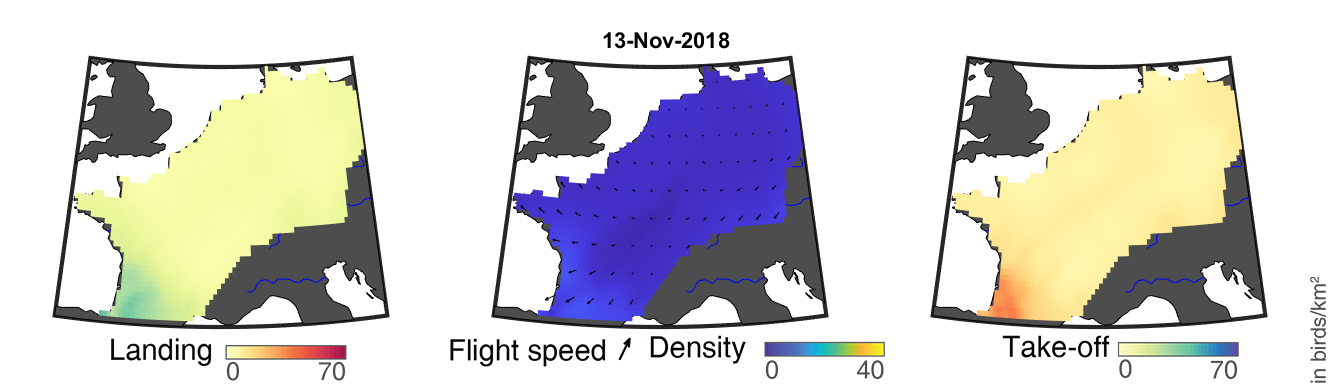

### day_274.png

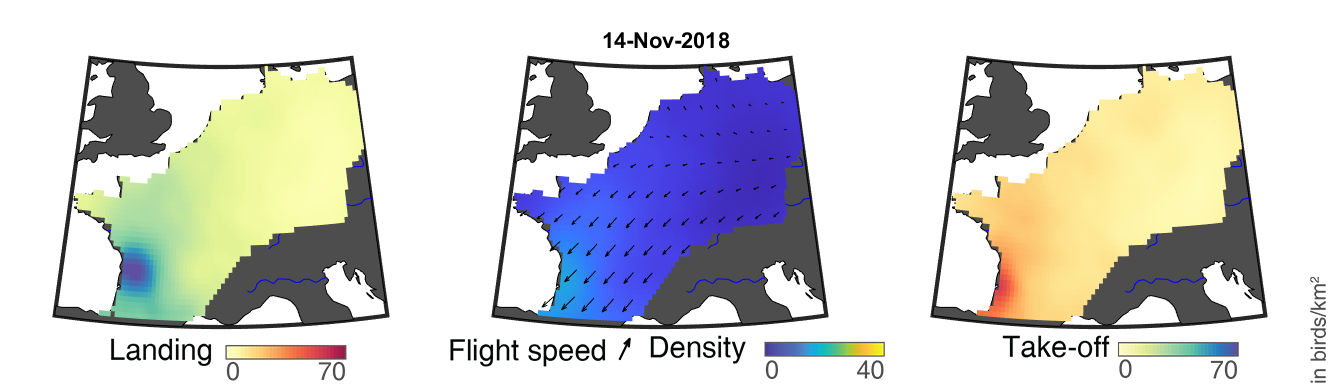

### day_275.png

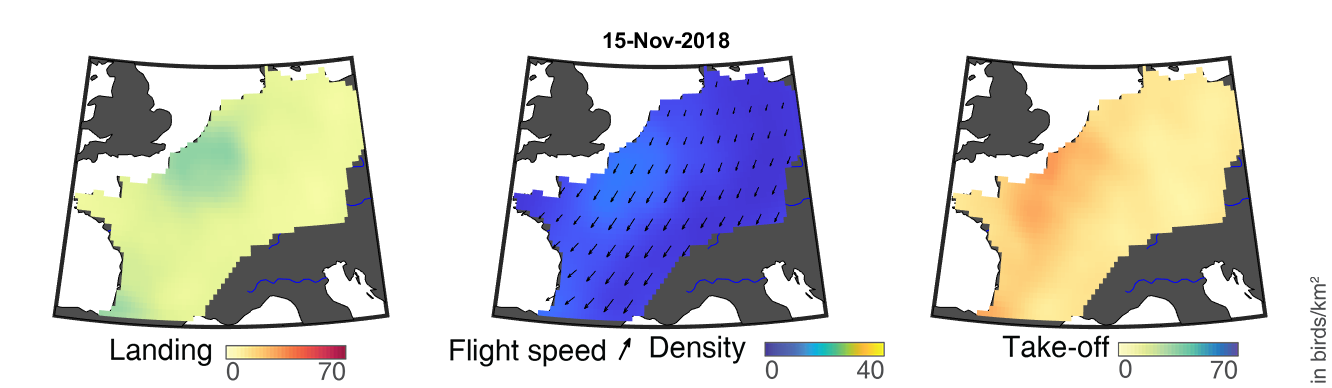

### day_276.png

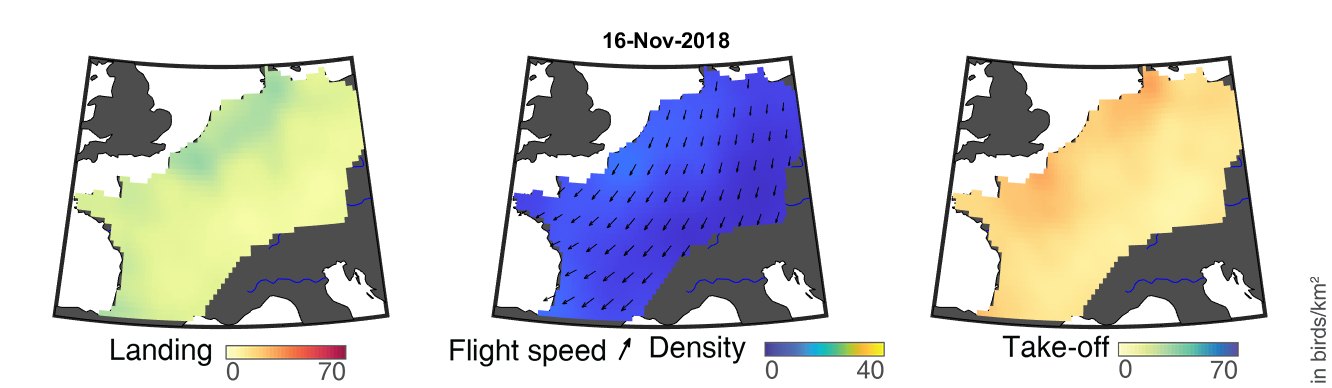

### day_277.png

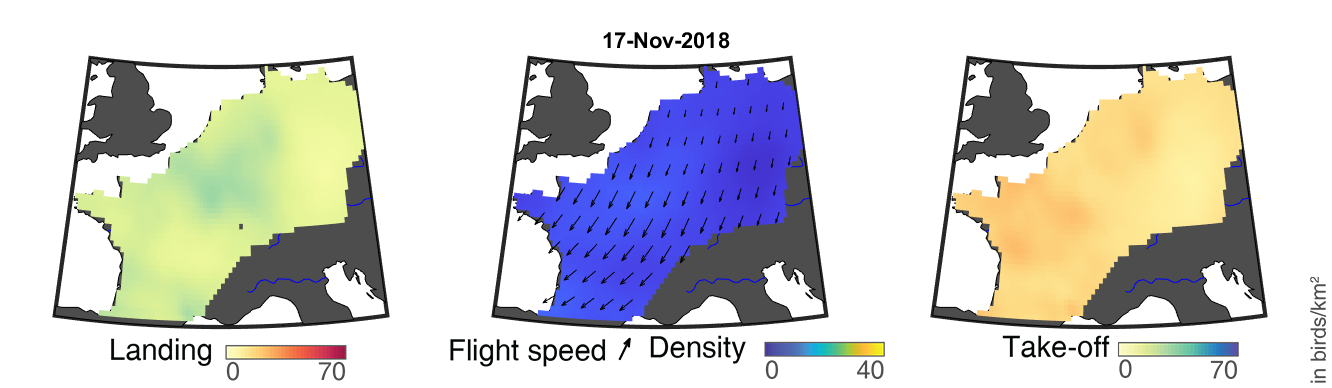

### day_278.png

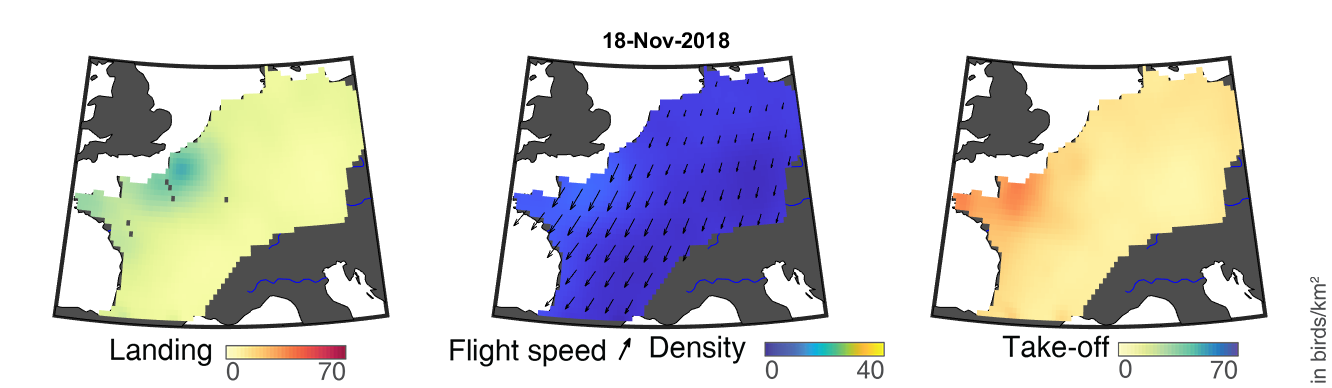

### day_279.png

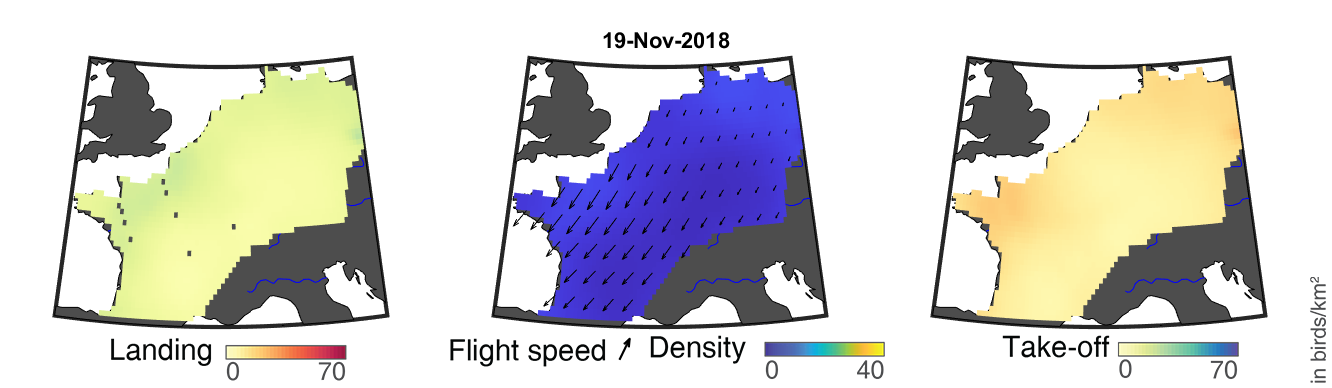
