## Supplementary figures and images for "Quantifying year-round nocturnal bird migration with a fluid dynamics model"

### day_1.png

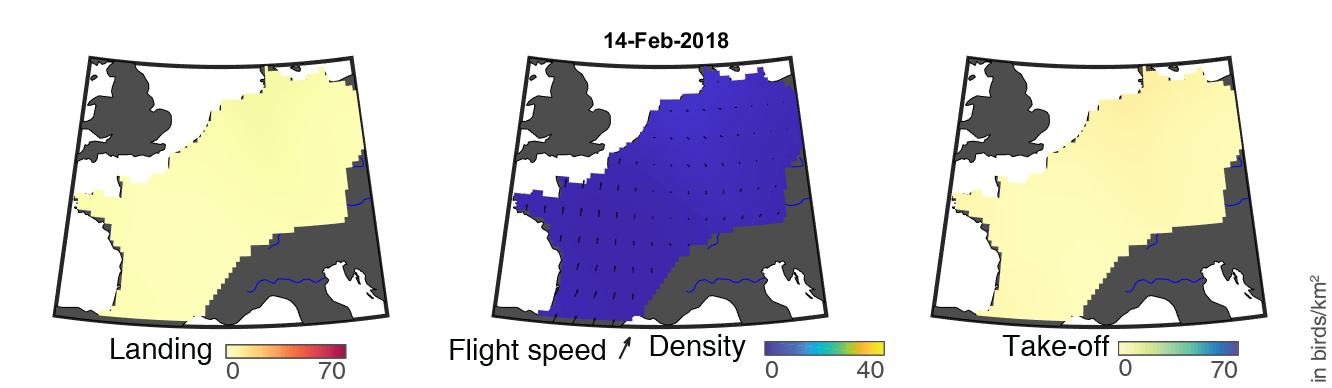

### day_10.png

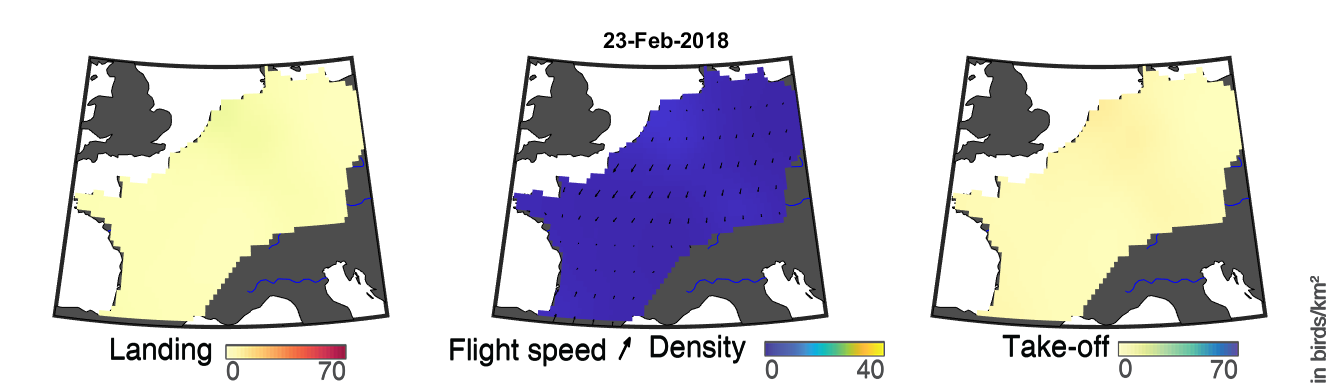

### day_11.png

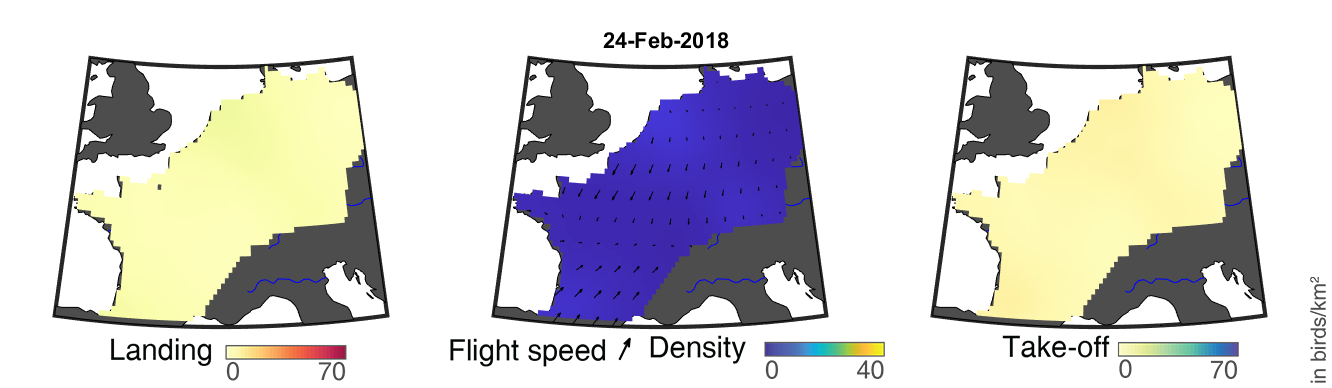

### day_12.png

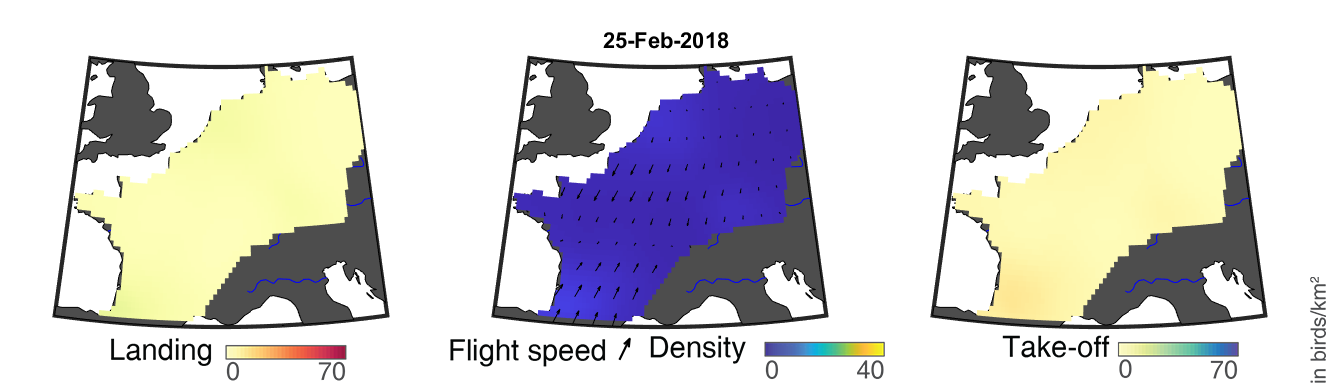

### day_13.png

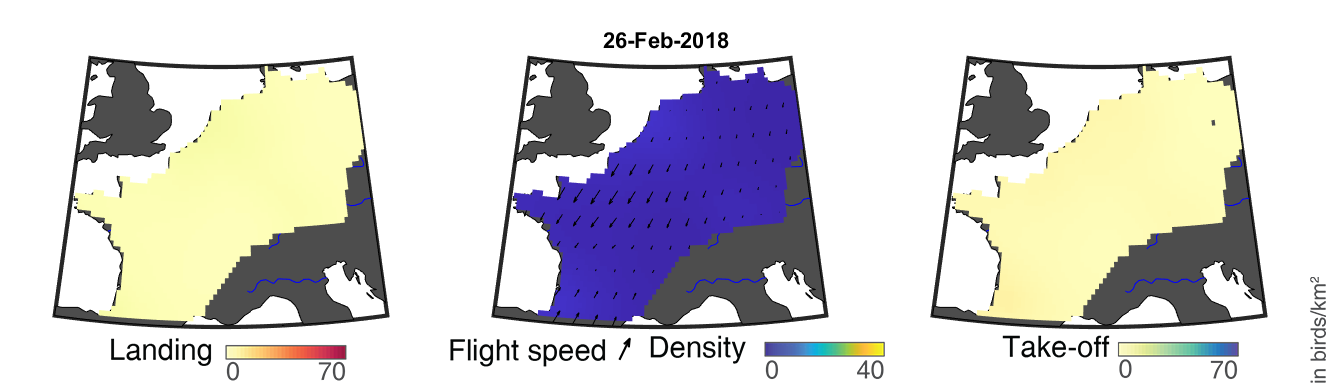

### day_14.png

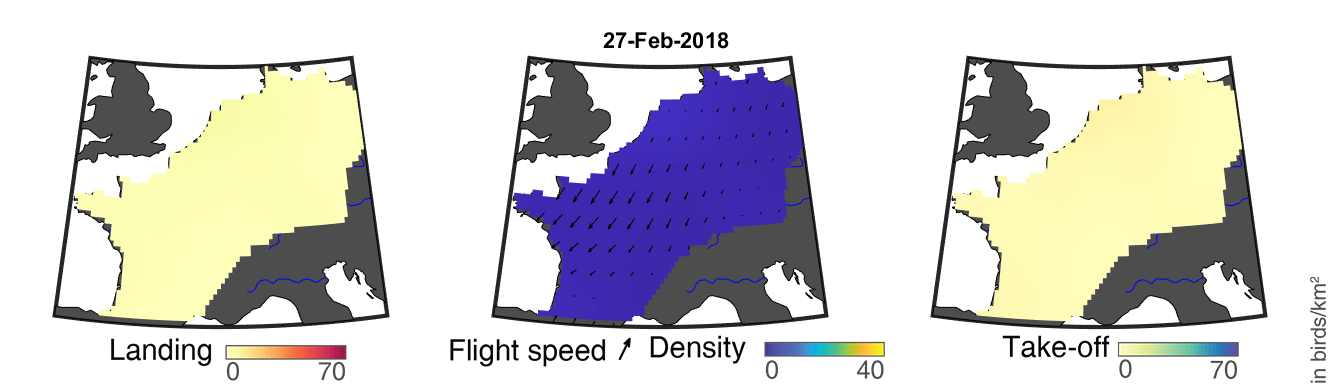

### day_15.png

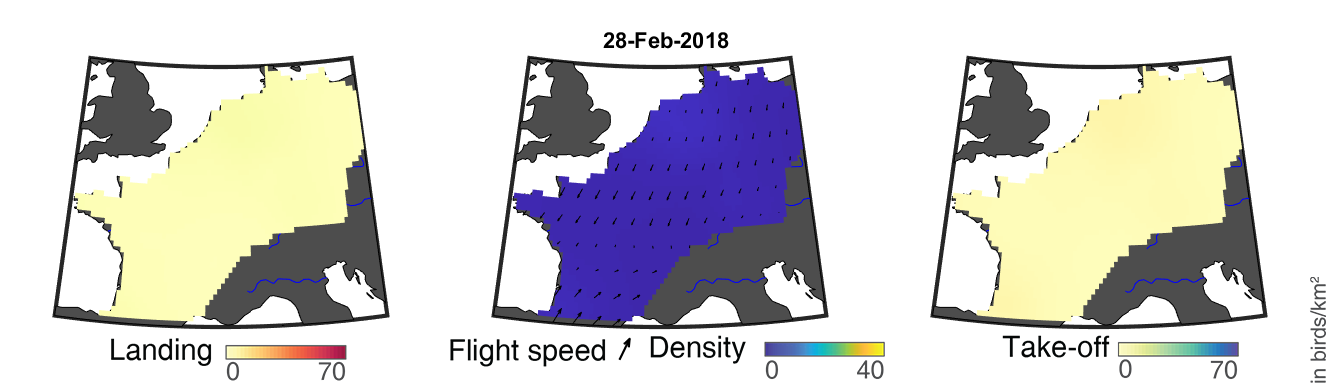

### day_16.png

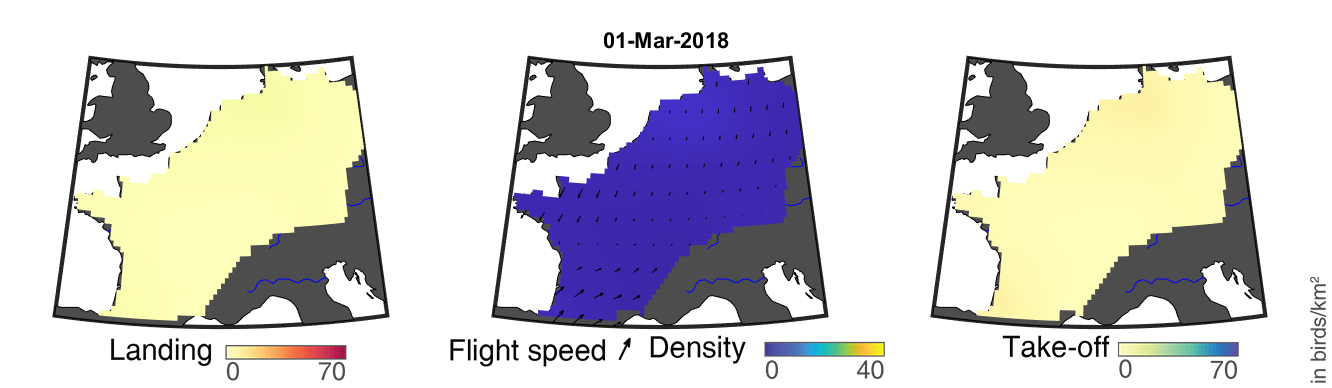

### day_17.png

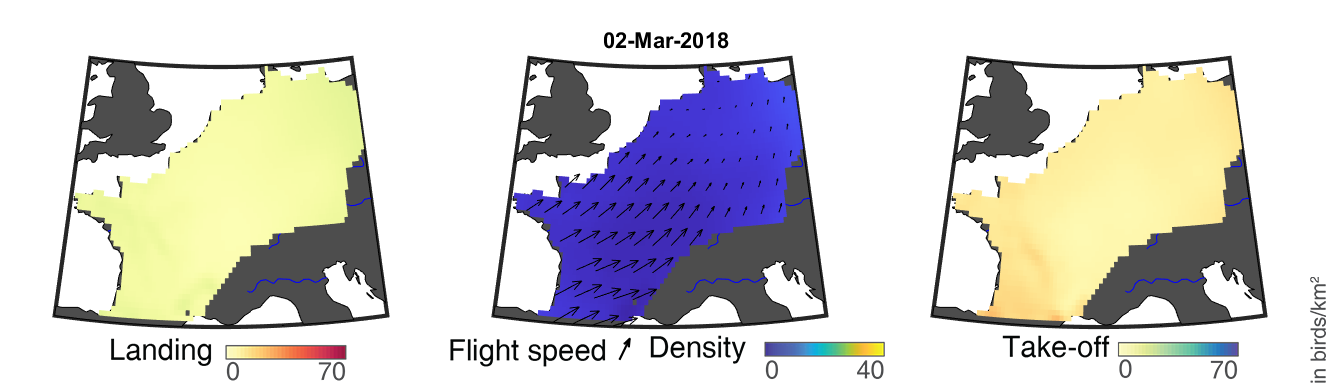

### day_18.png

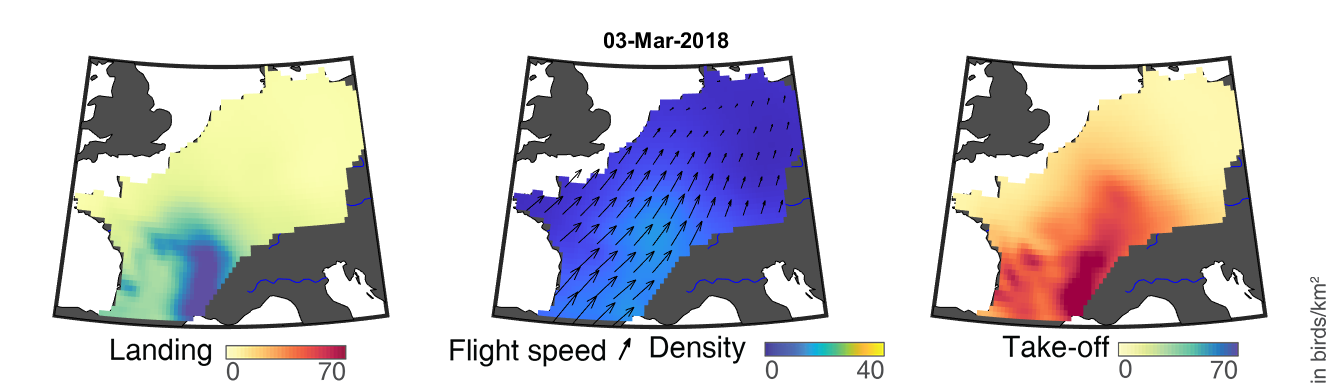

### day_100.png

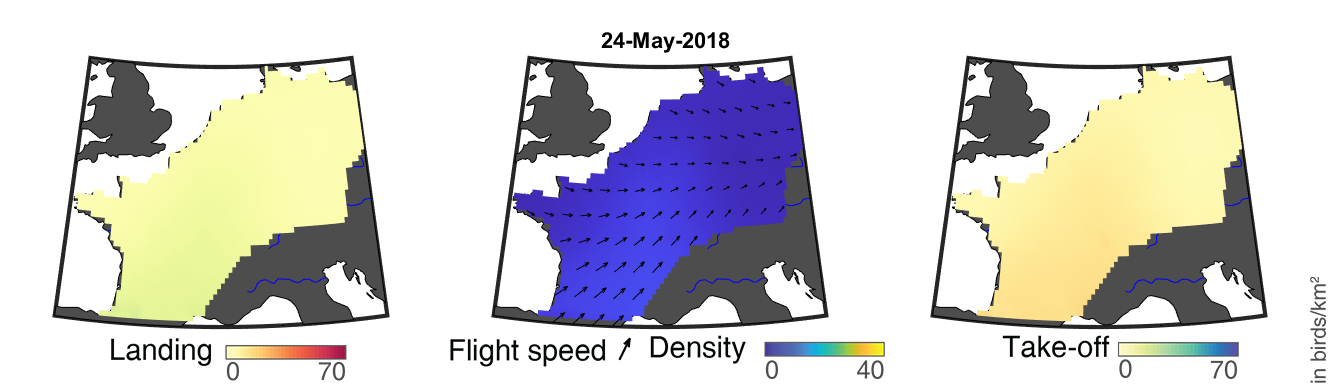

### day_101.png

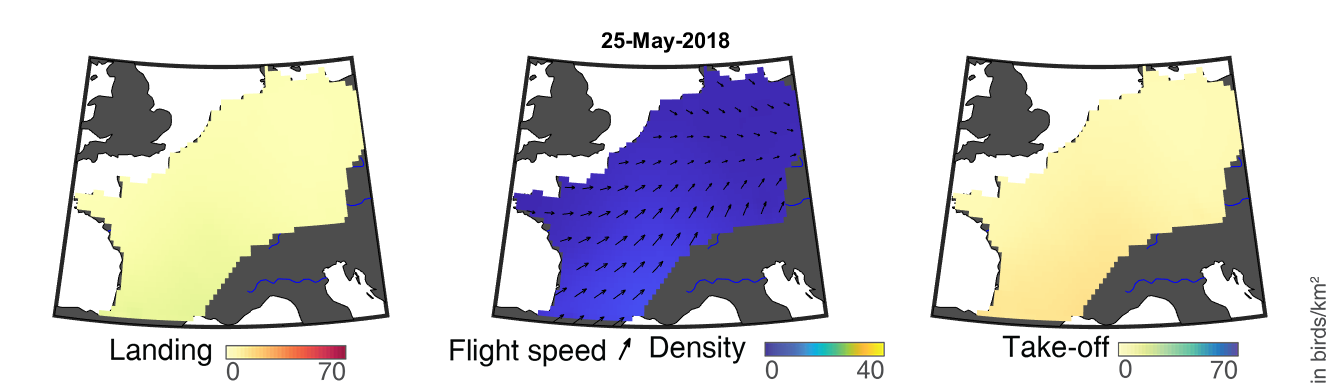

### day_102.png

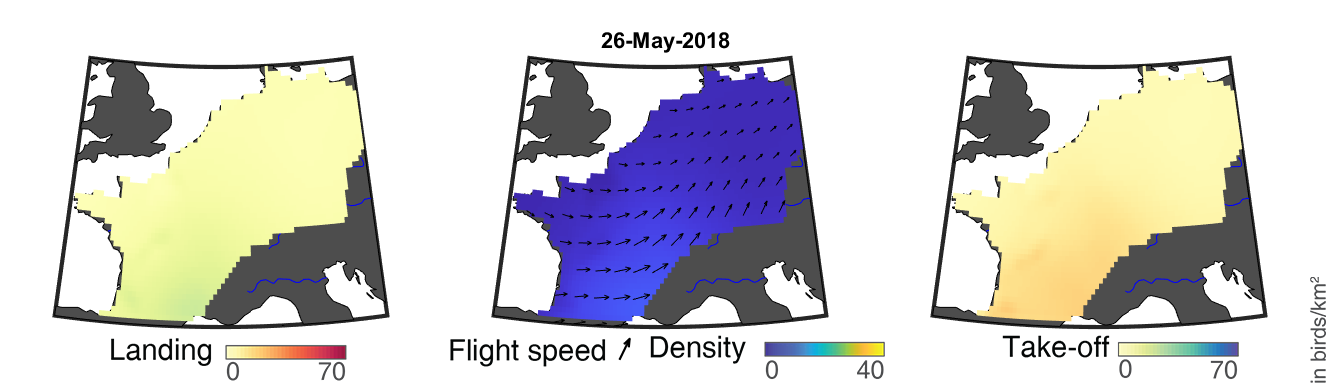

### day_103.png

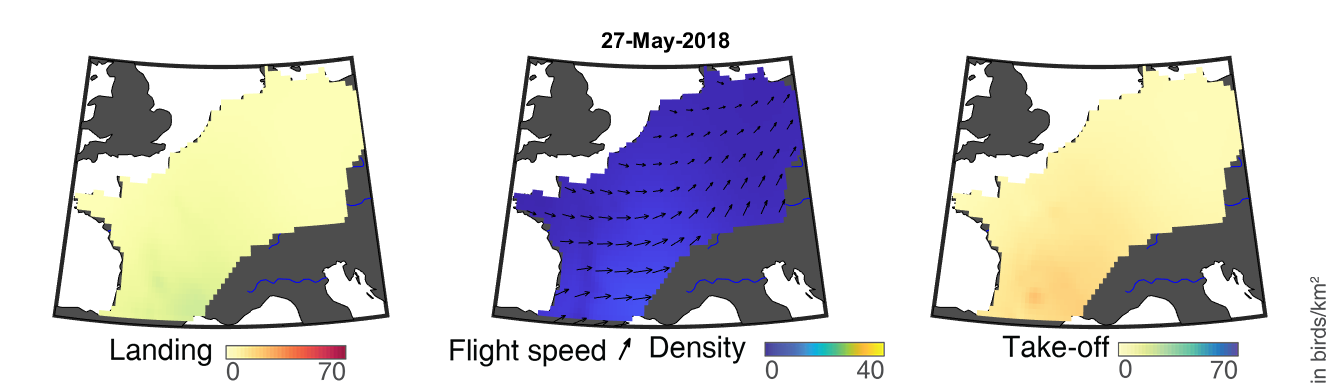

### day_104.png

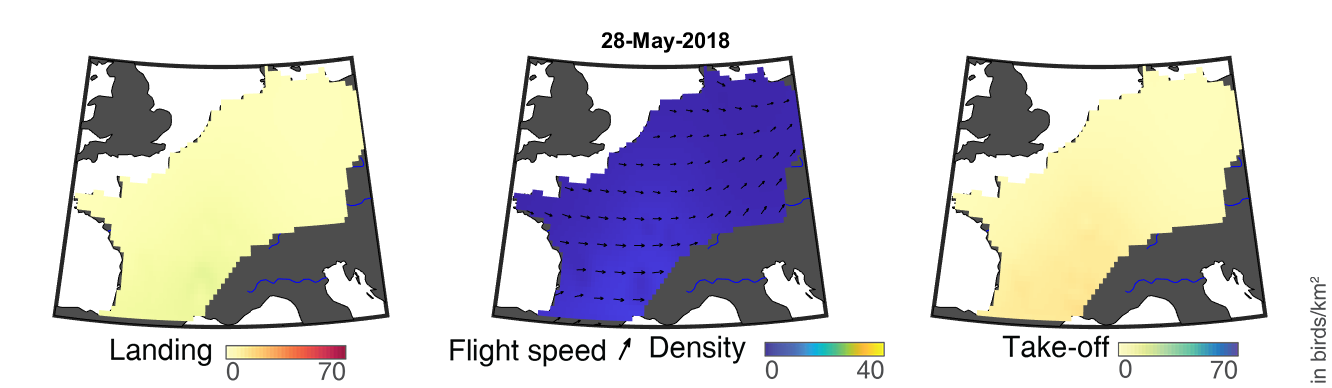

### day_105.png

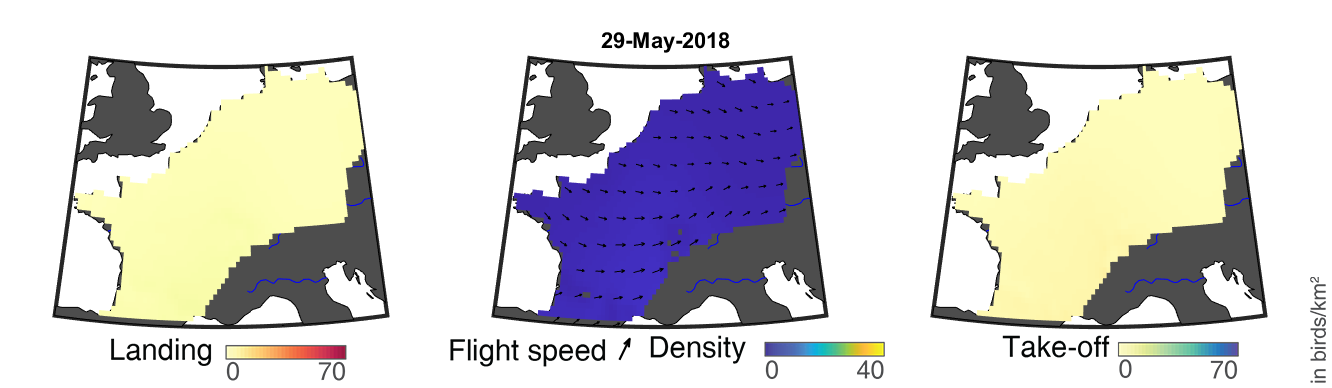

### day_106.png

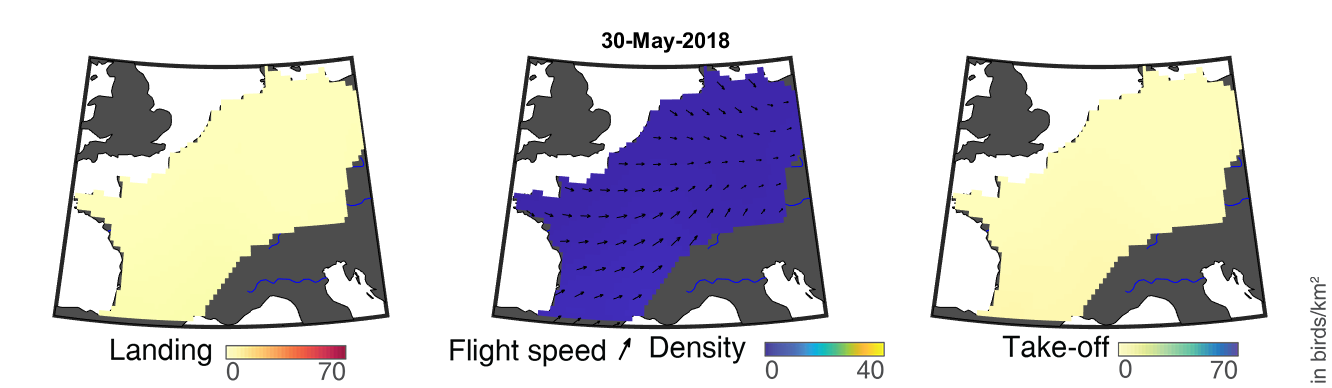

### day_107.png

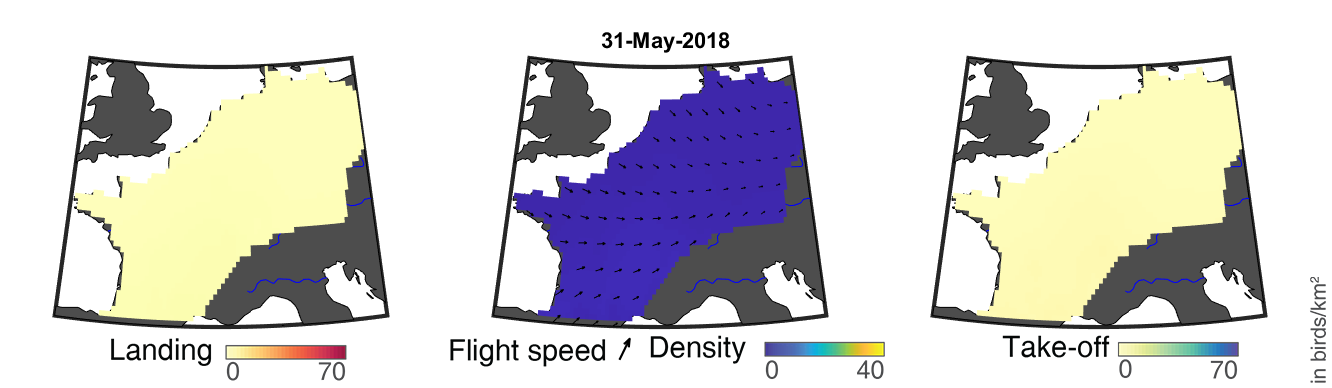

### day_108.png

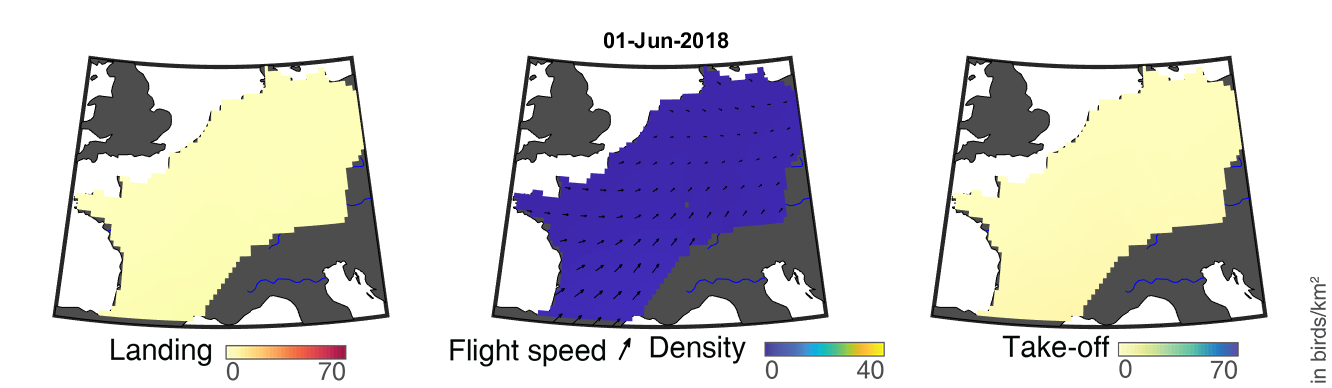

### day_109.png

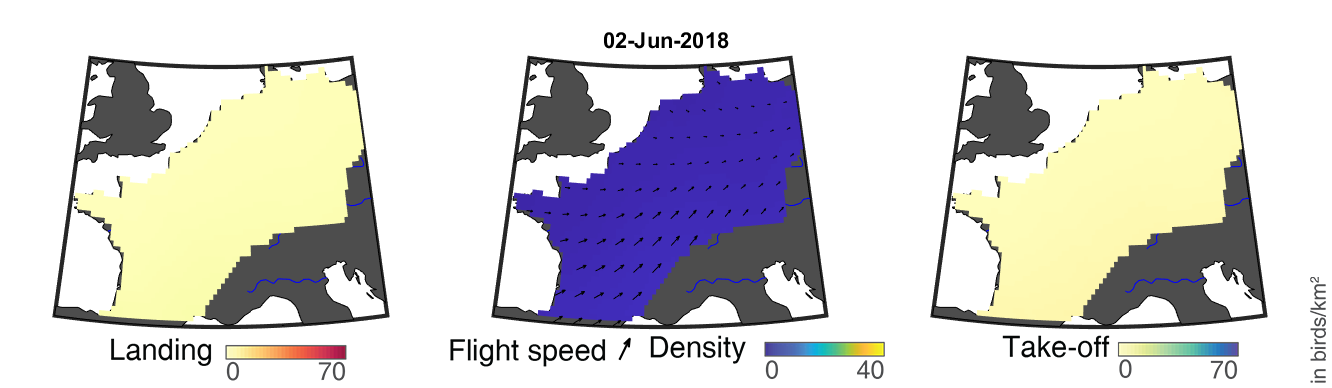

### day_110.png

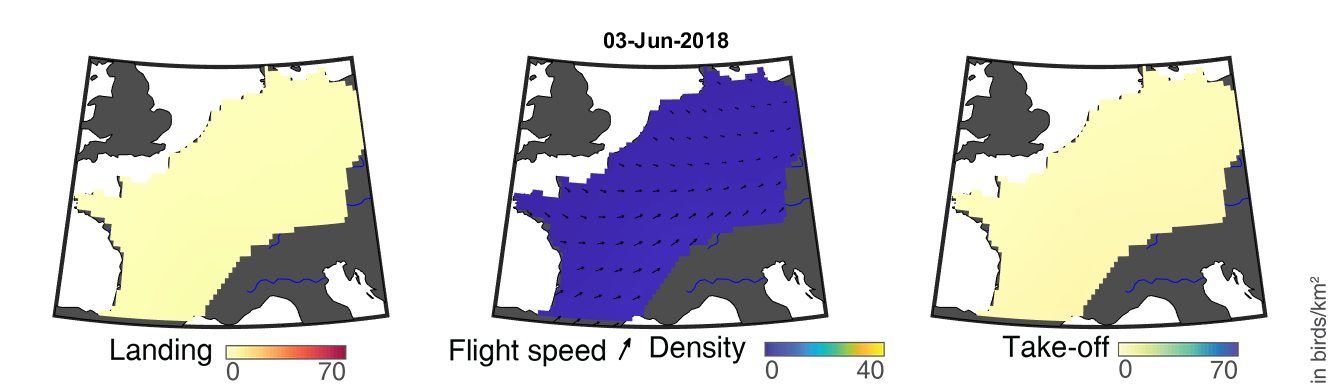

### day_111.png

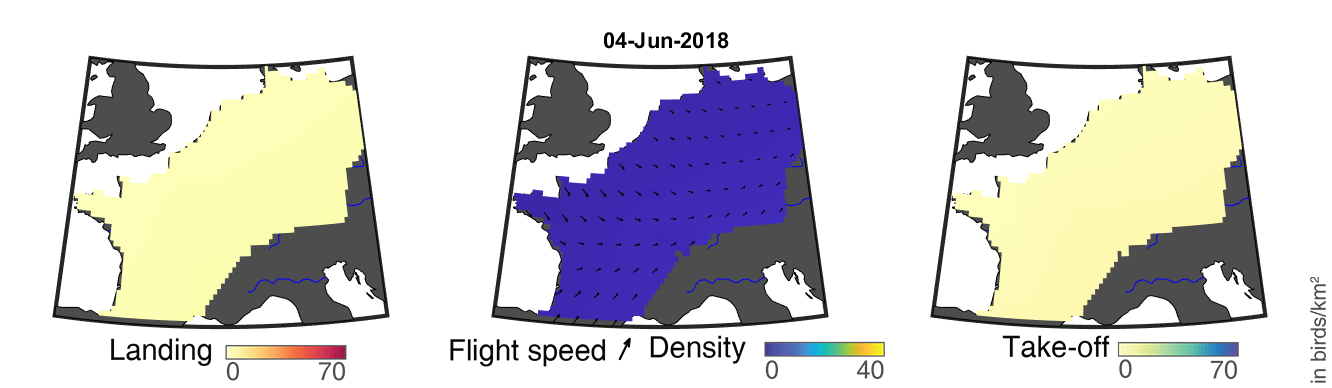

### day_112.png

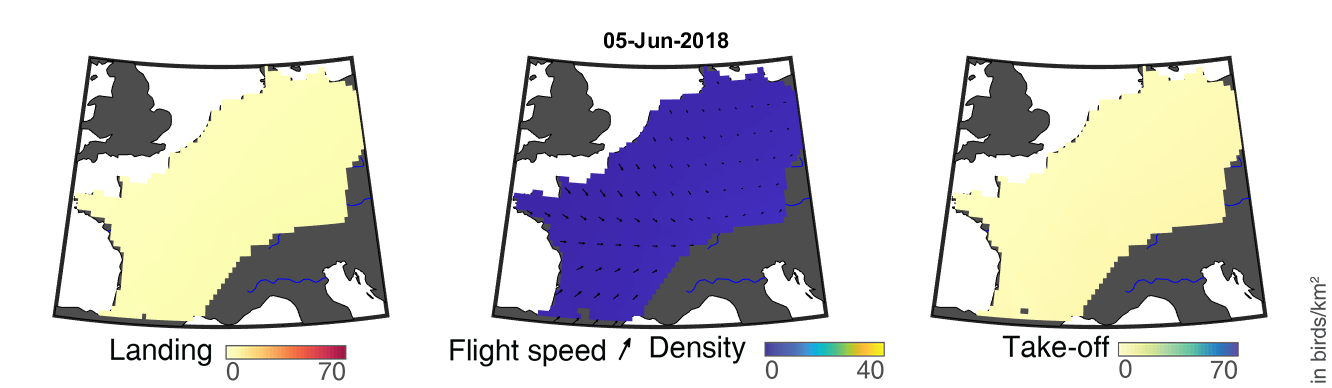

### day_113.png

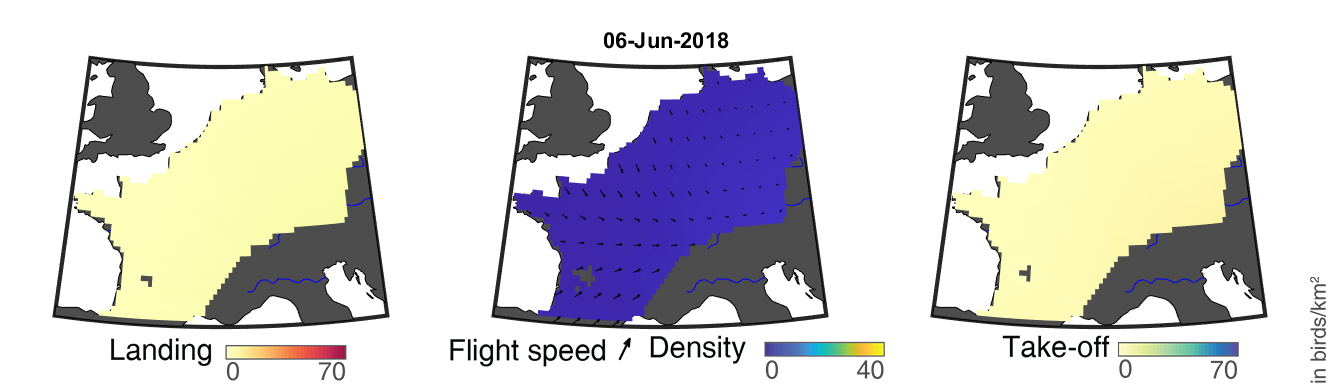

### day_114.png

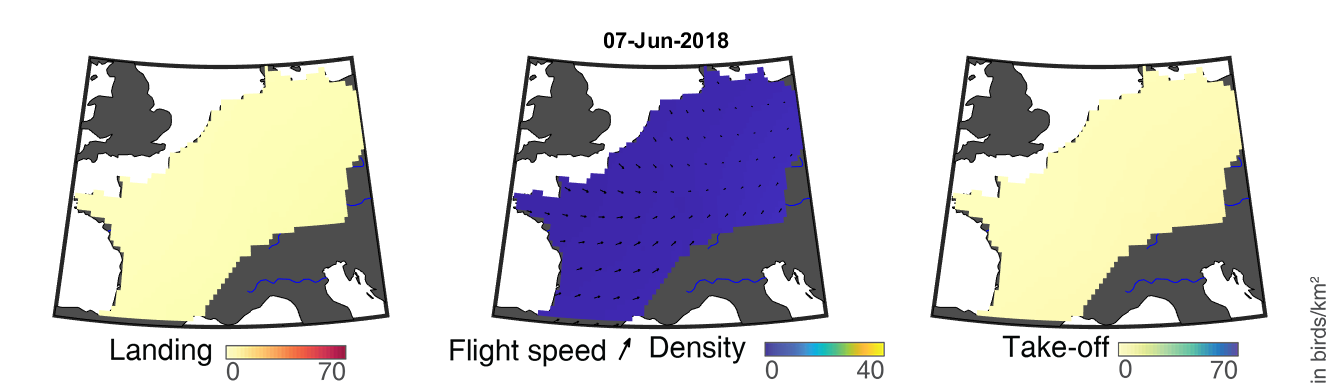

### day_115.png

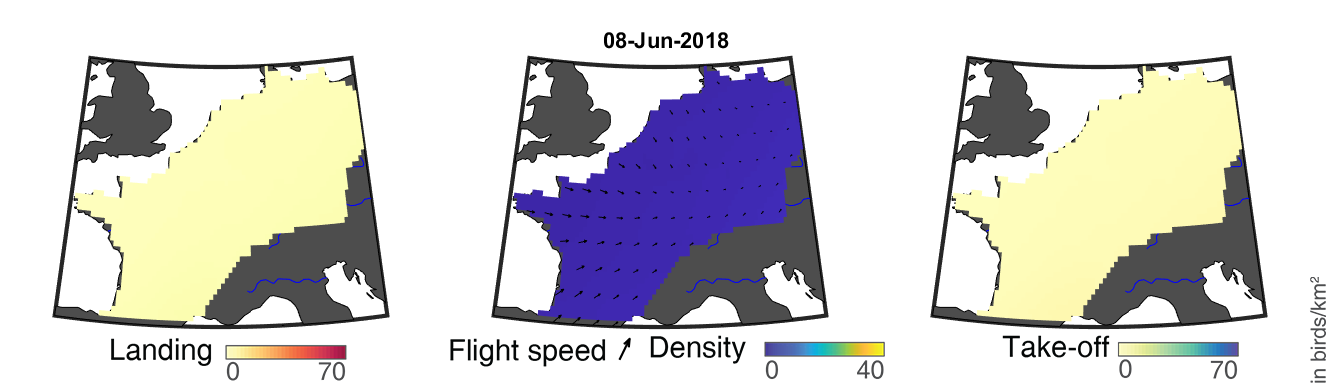

### day_116.png

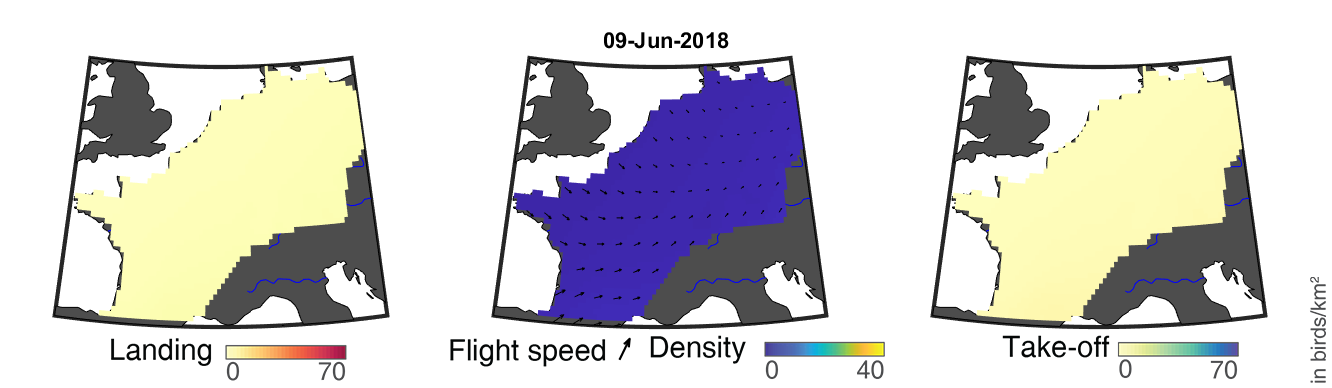

### day_117.png

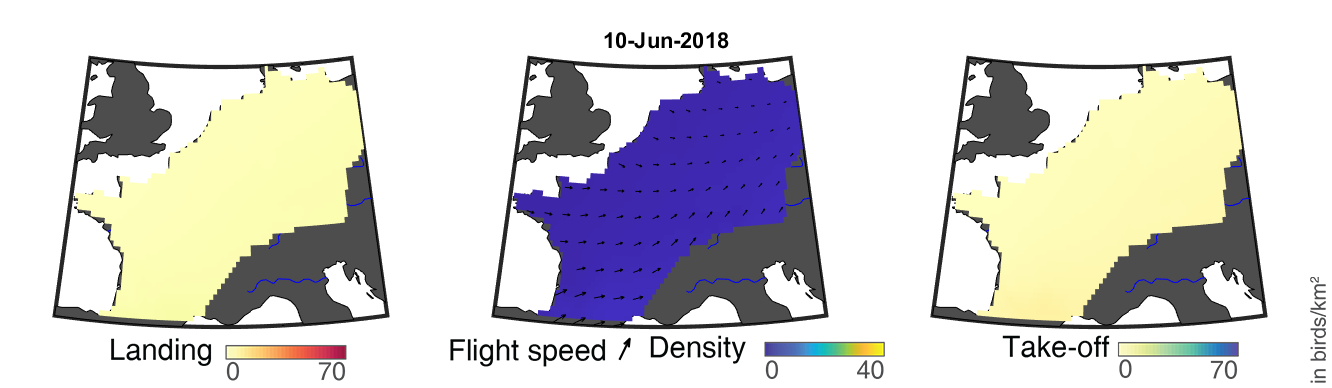

### day_118.png

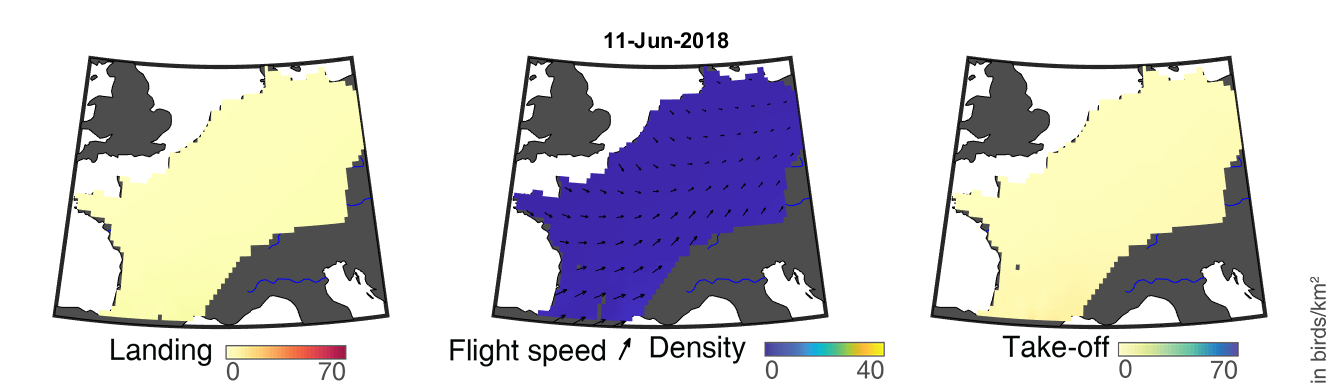

### day_119.png

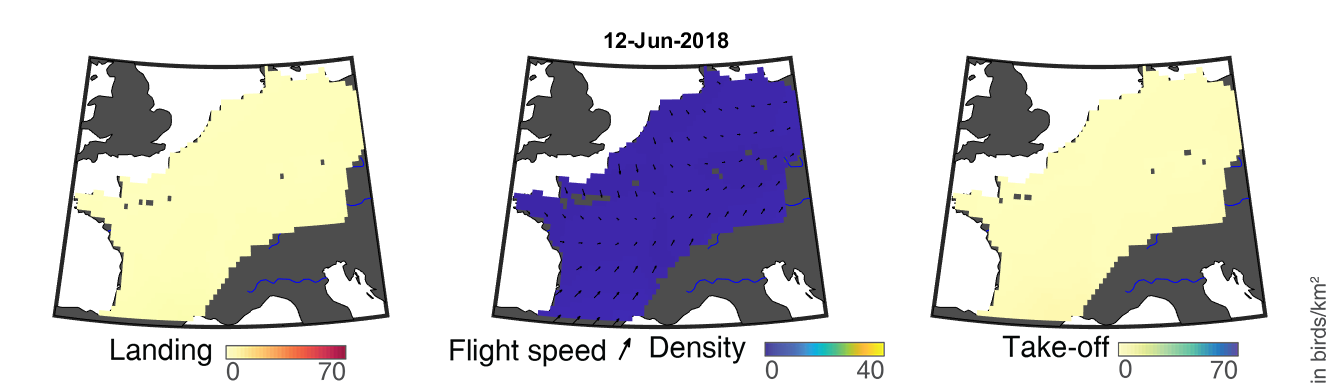

### day_120.png

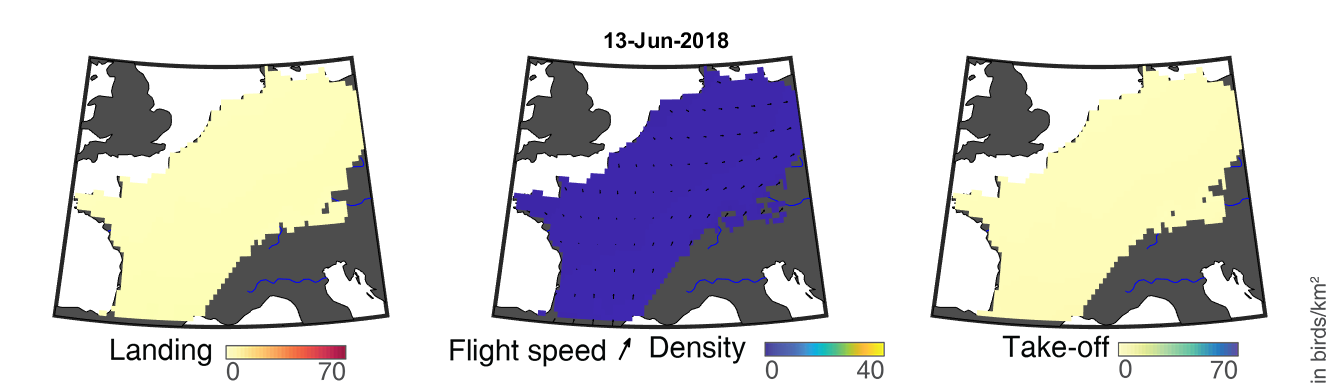

### day_121.png

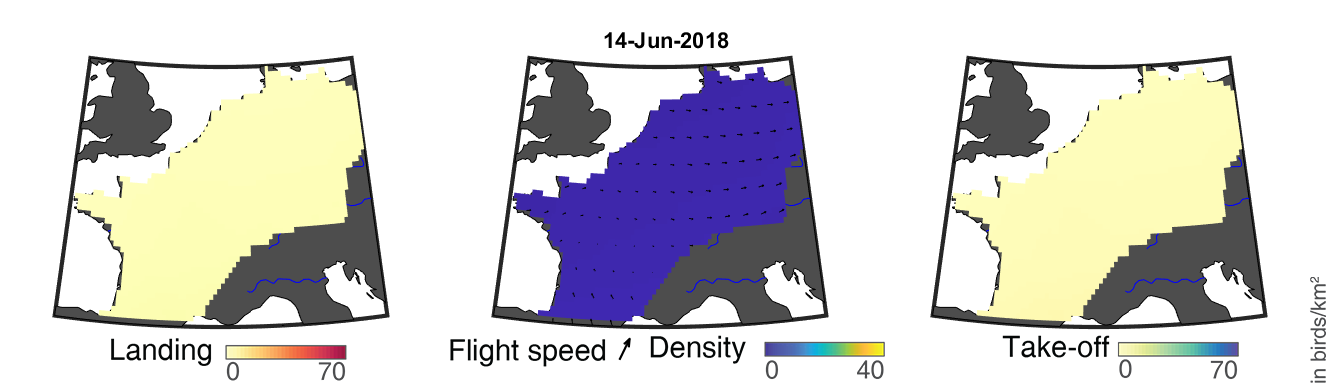

### day_122.png

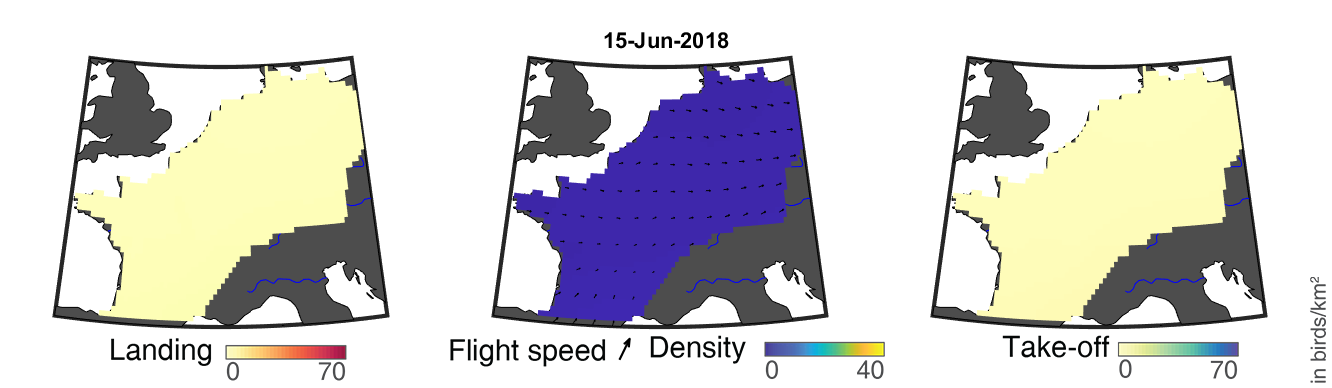

### day_123.png

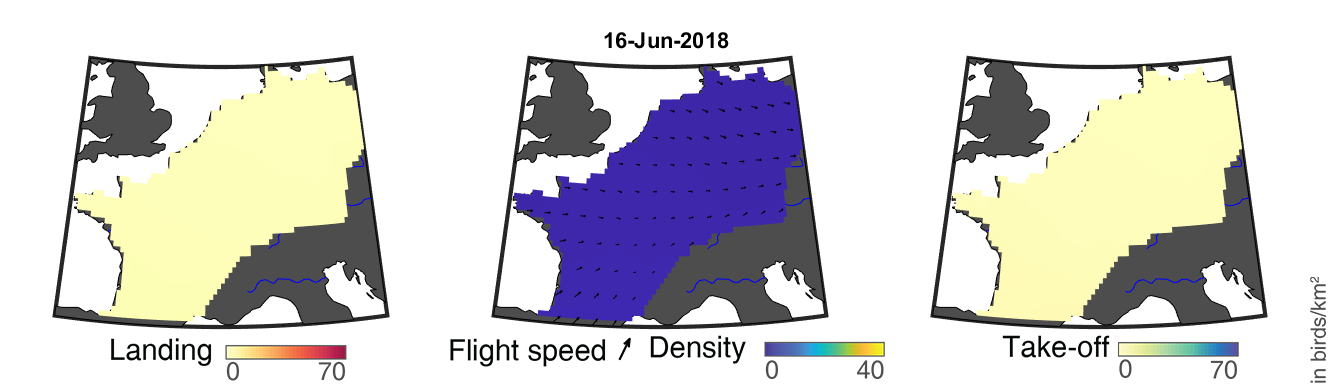

### day_124.png

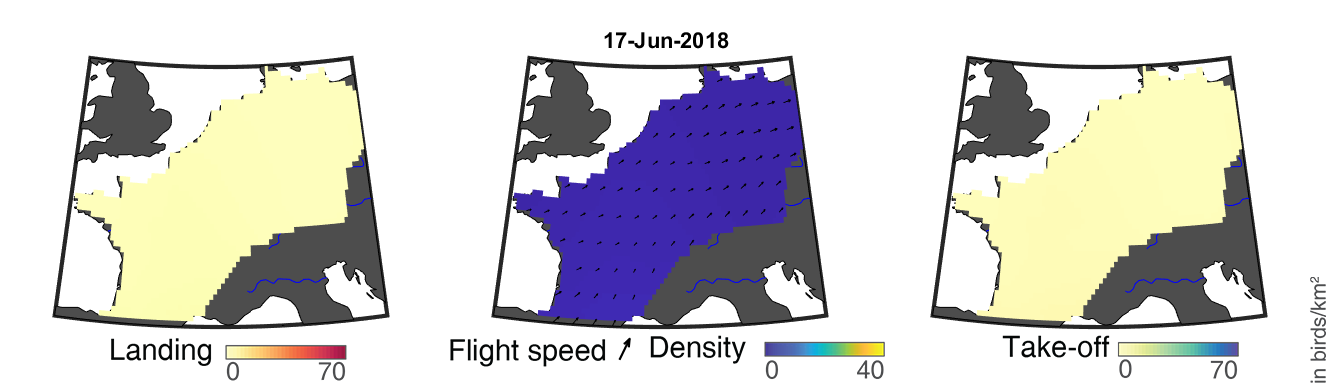

### day_125.png

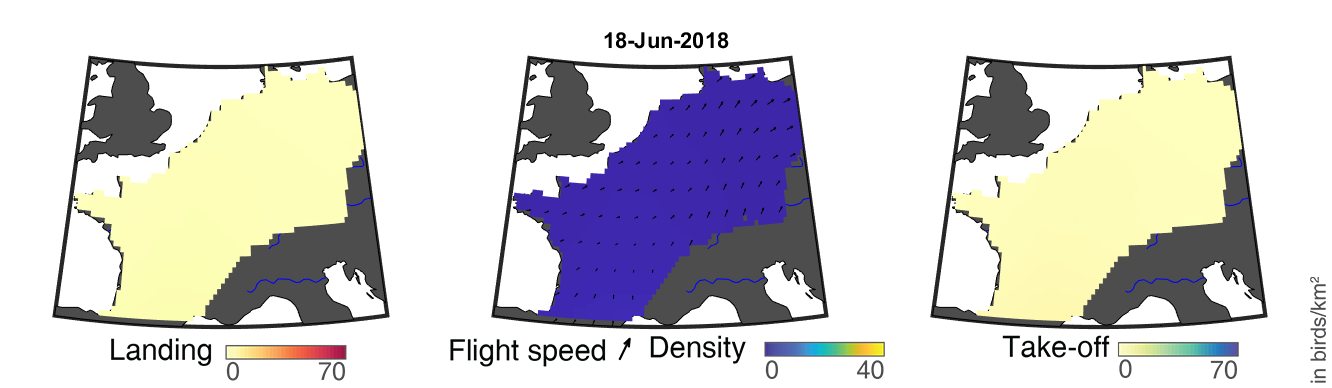

### day_126.png

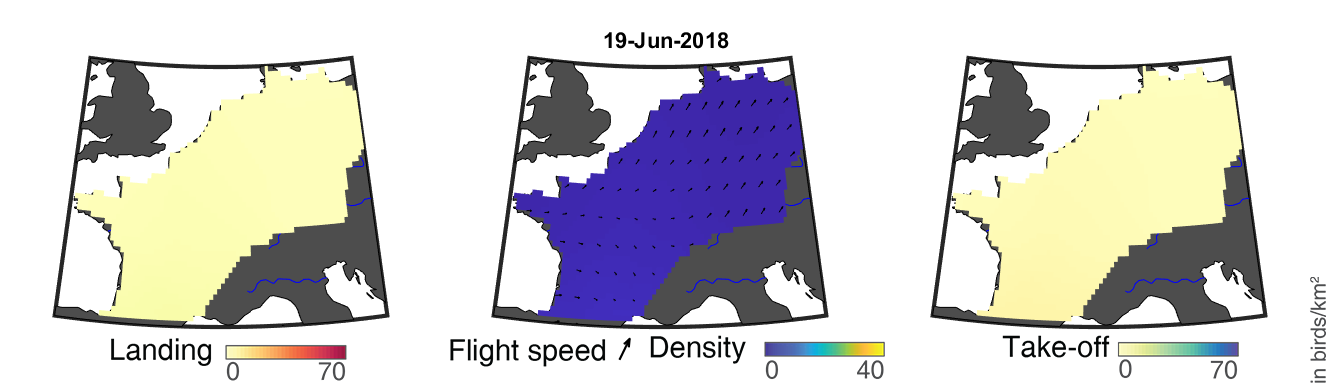

### day_127.png

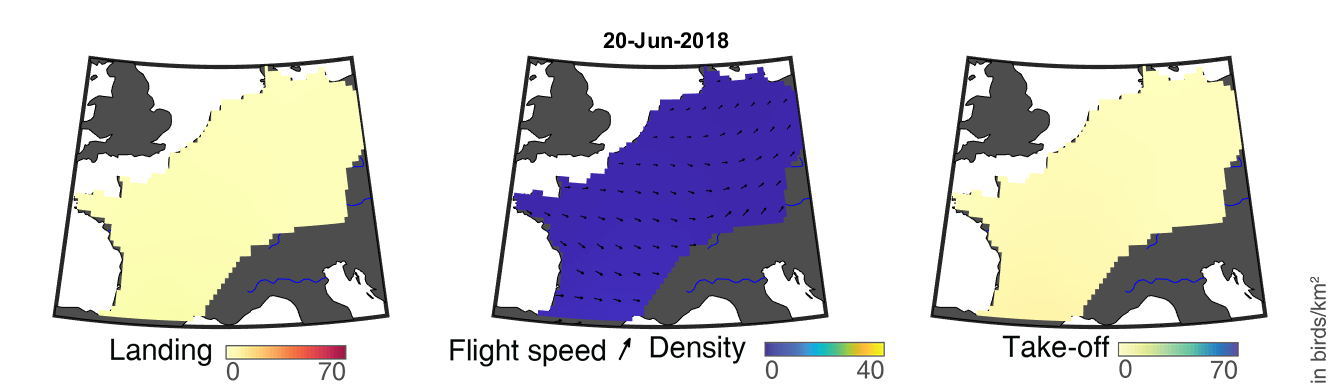

### day_128.png

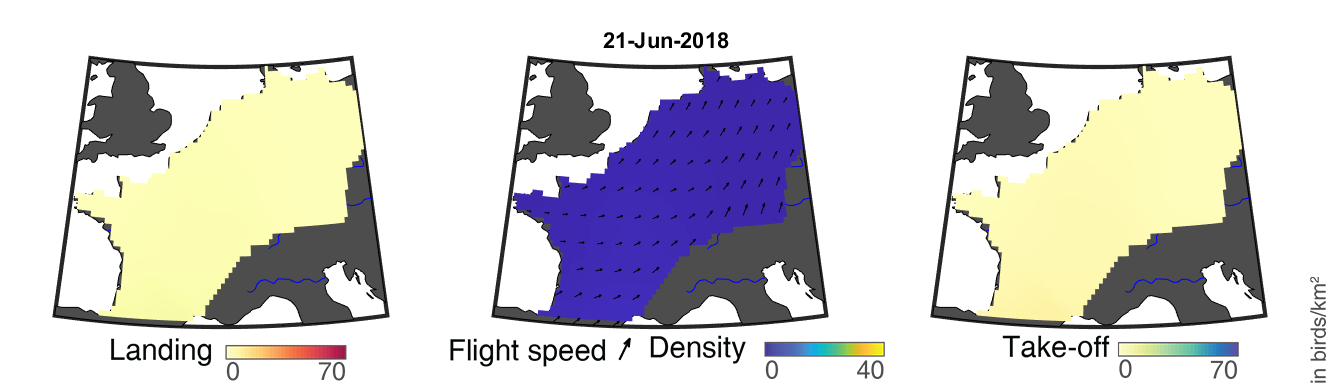

### day_129.png

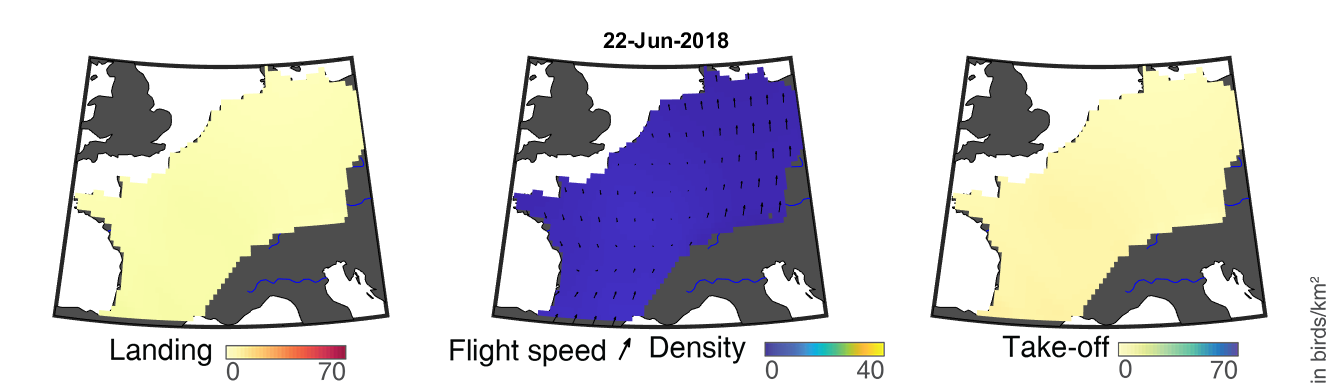

### day_130.png

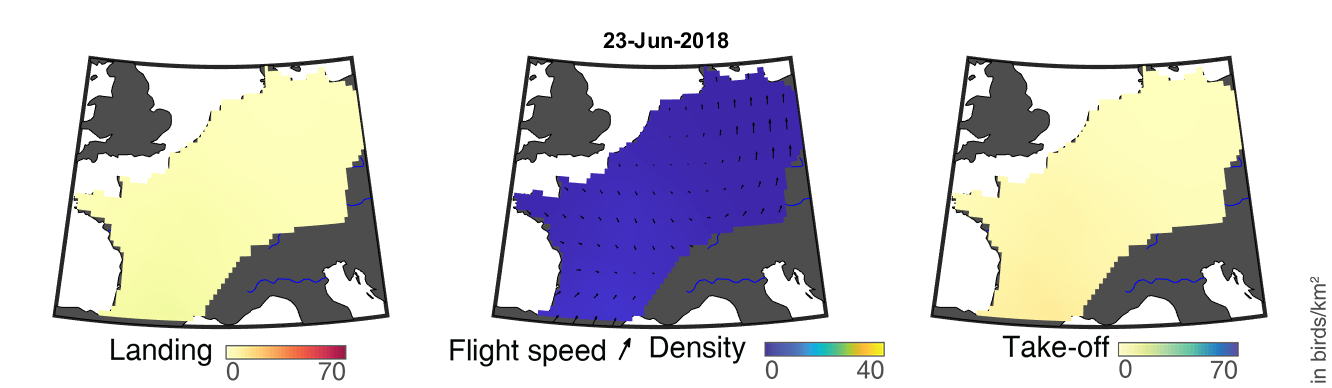

### day_131.png

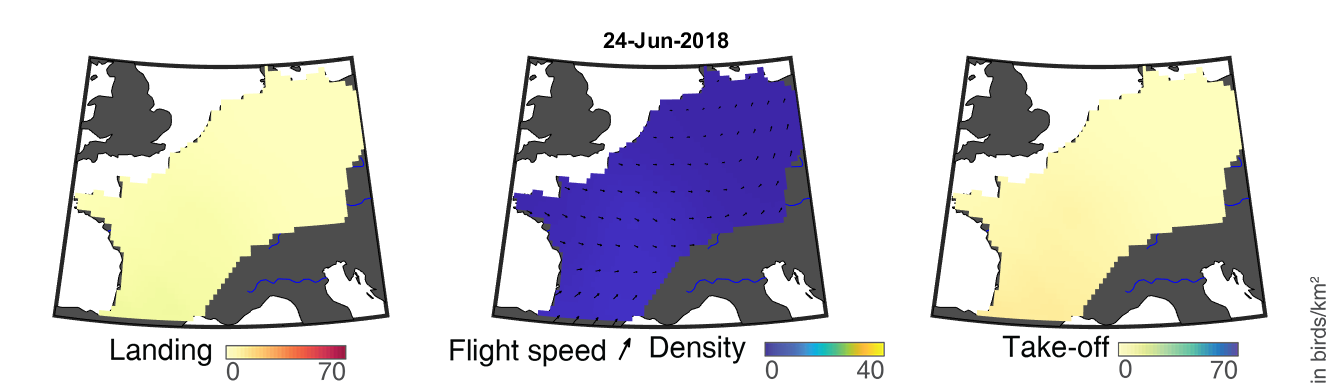

### day_132.png

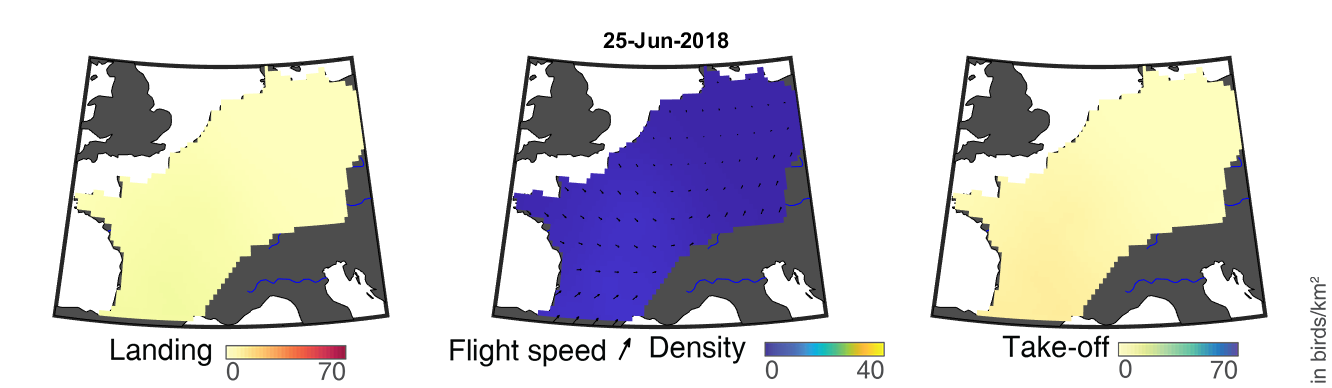

### day_133.png

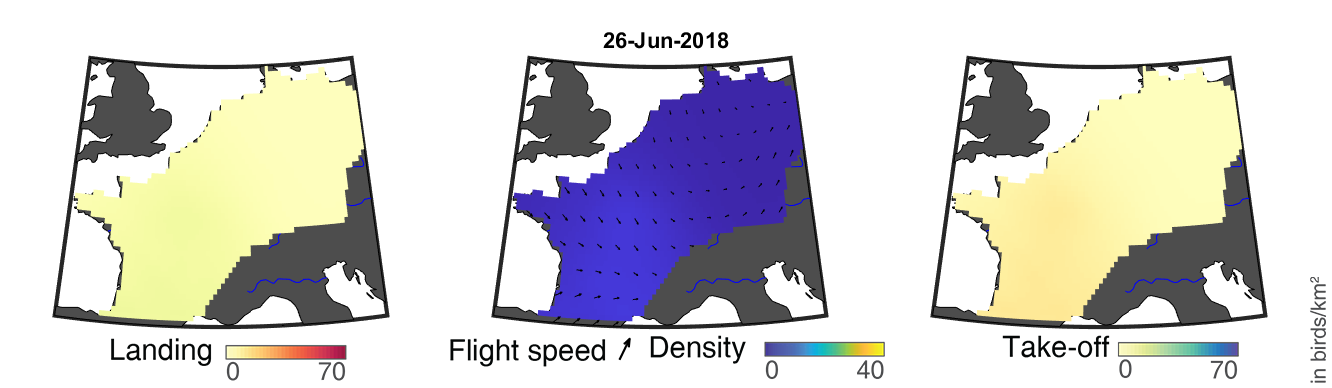

### day_134.png

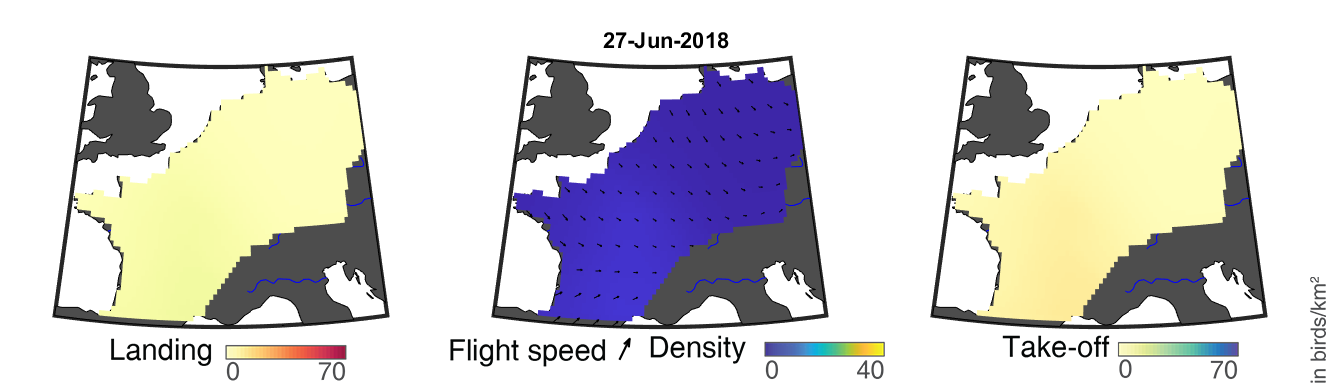

### day_135.png

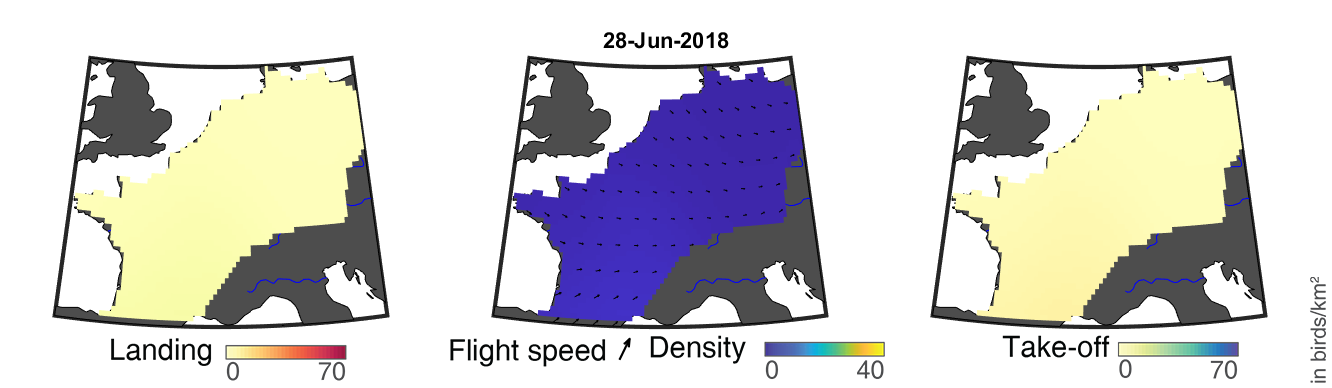

### day_136.png

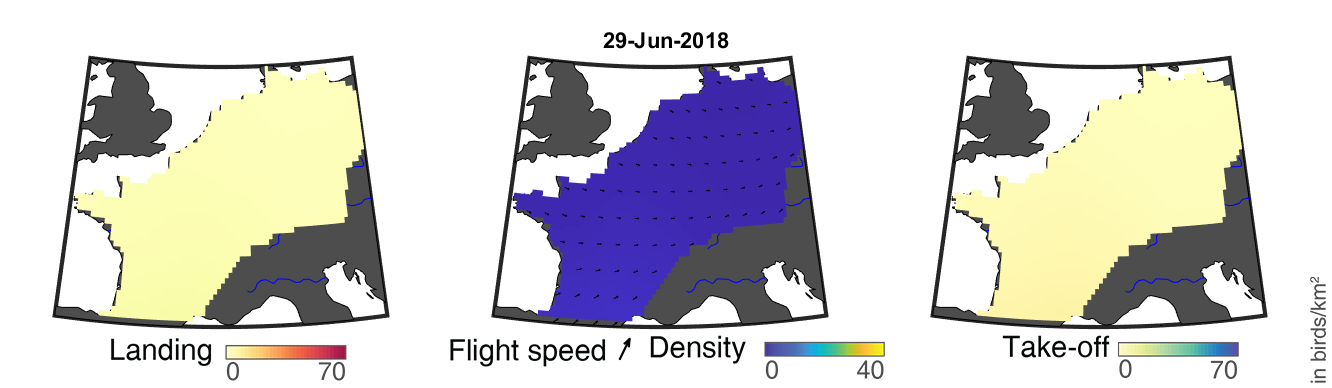

### day_137.png

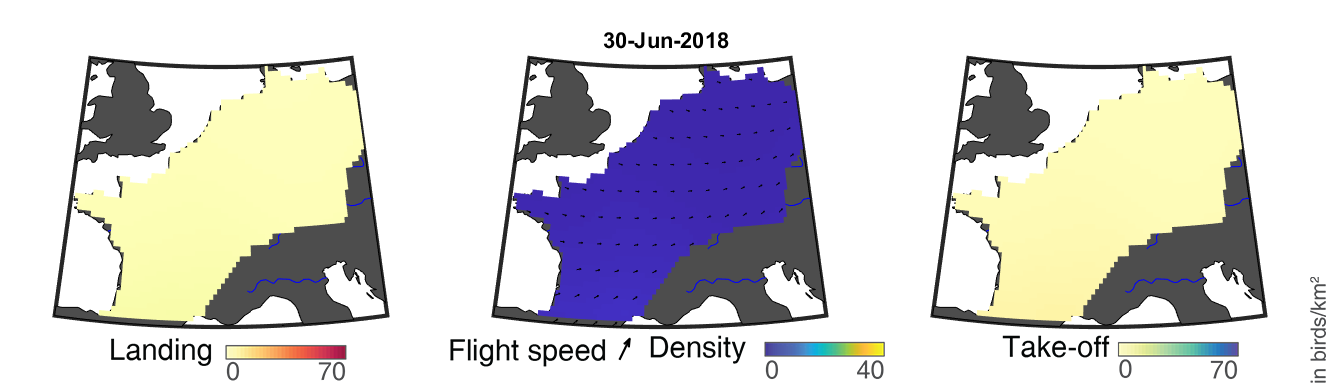

### day_138.png

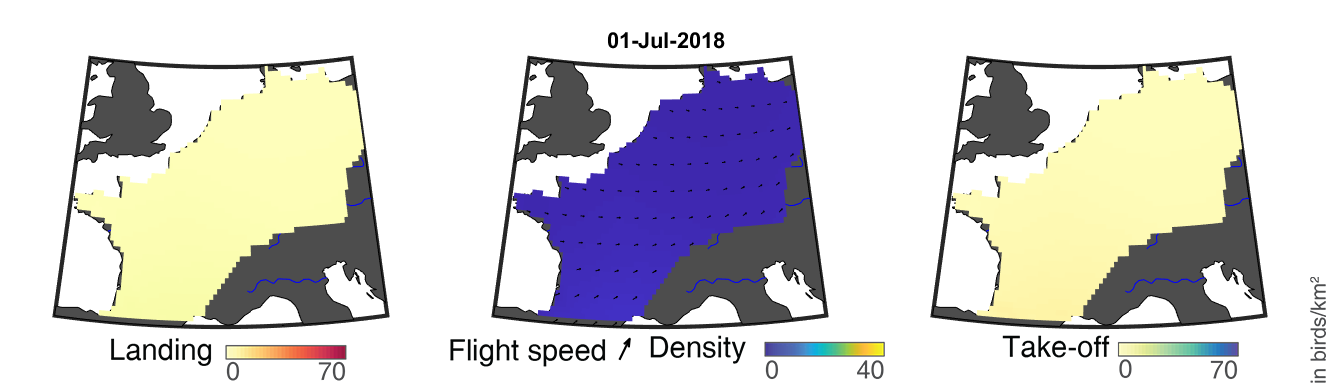

### day_139.png

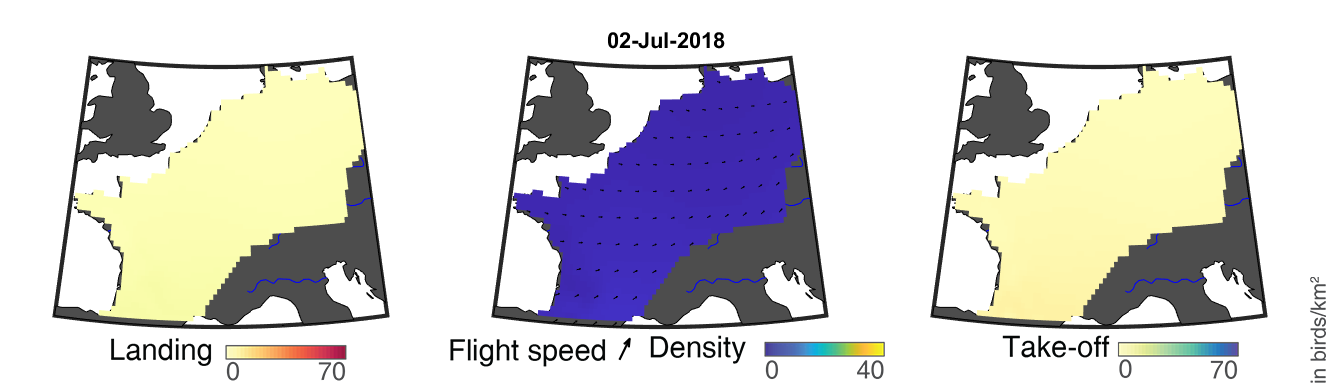

### day_140.png

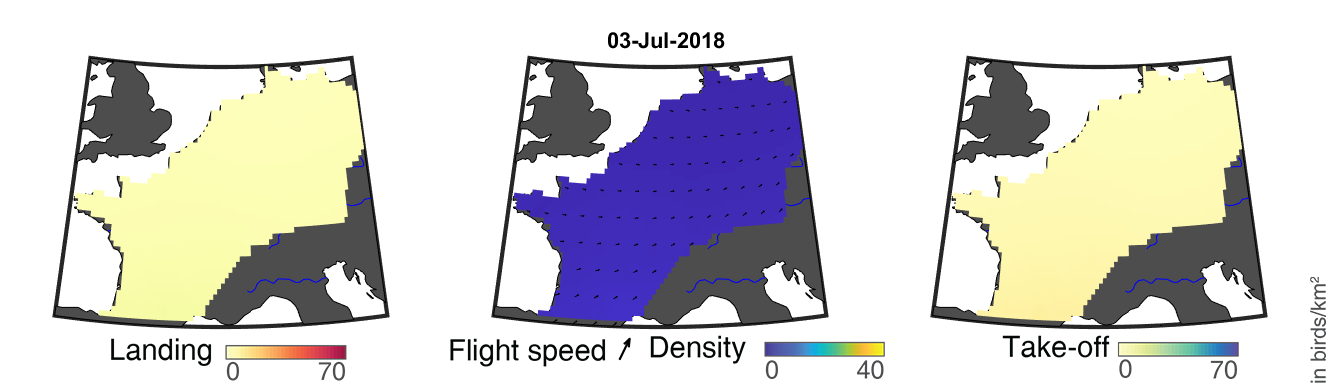

### day_141.png

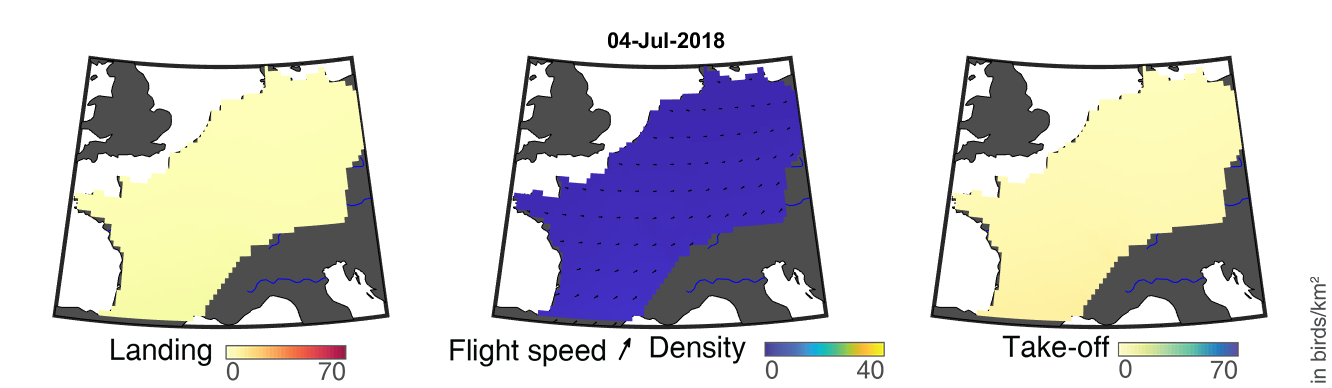

### day_142.png

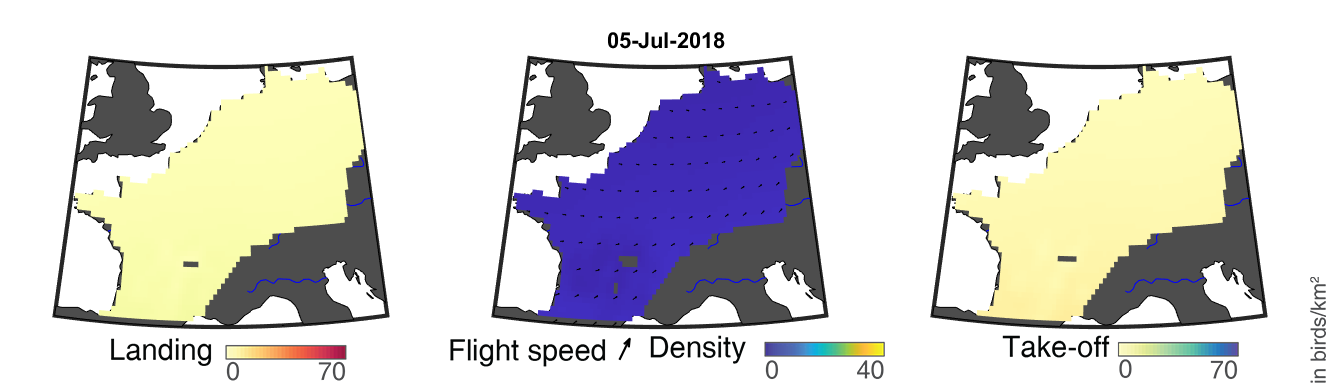

### day_143.png

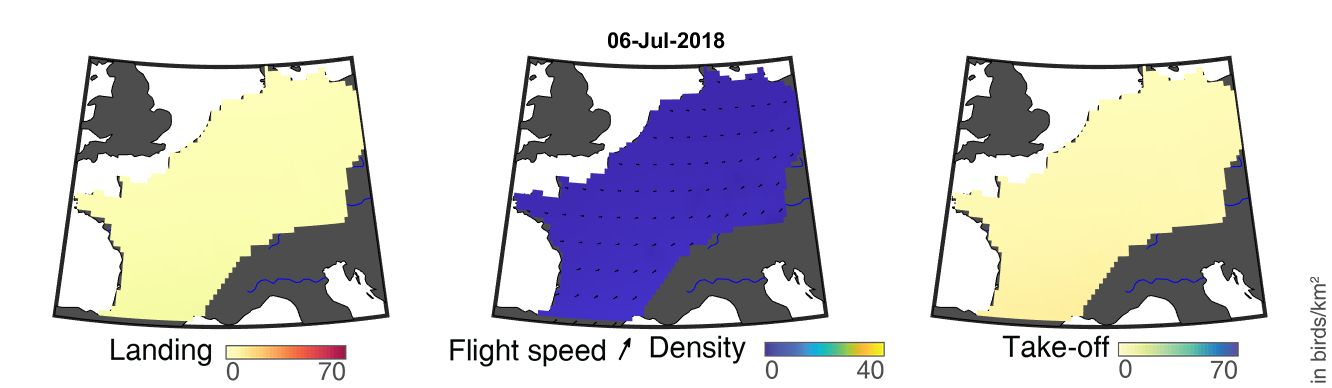

### day_144.png

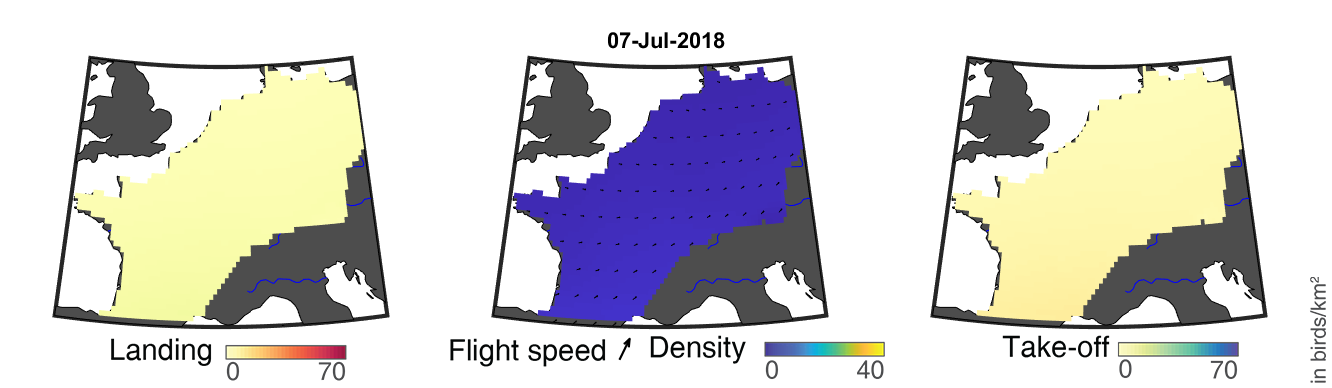

### day_145.png

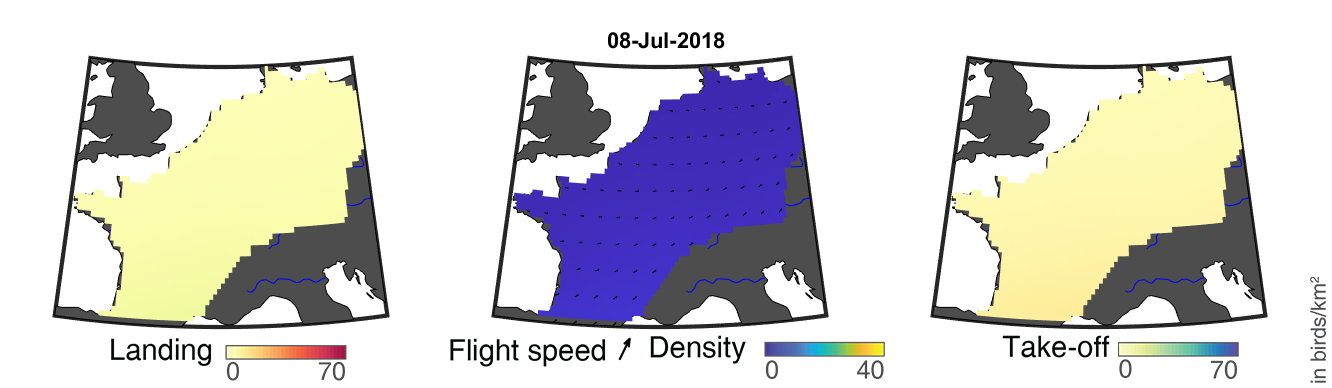

### day_146.png

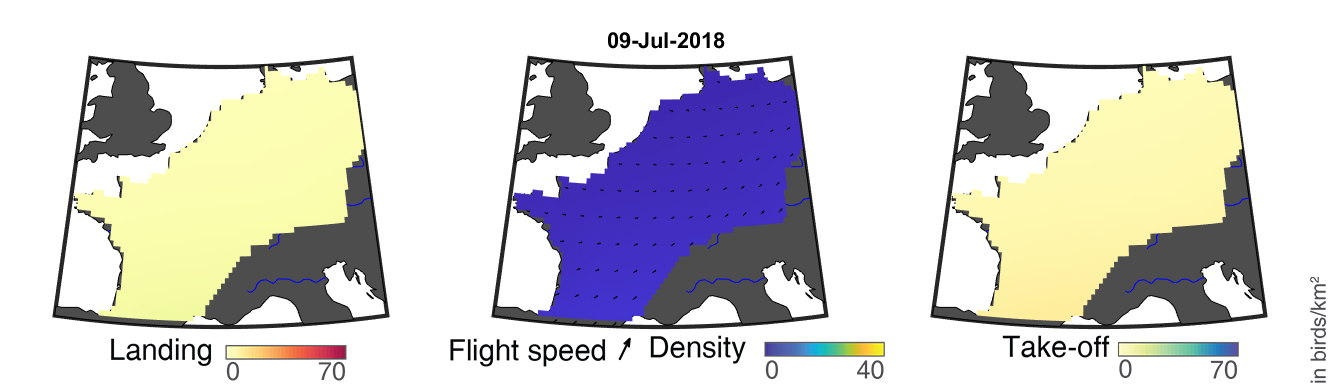

### day_147.png

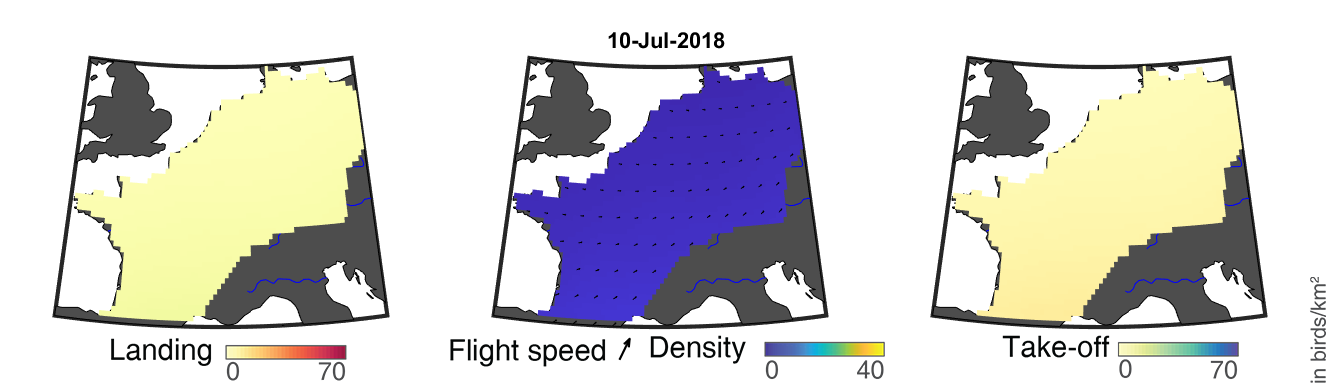

### day_148.png

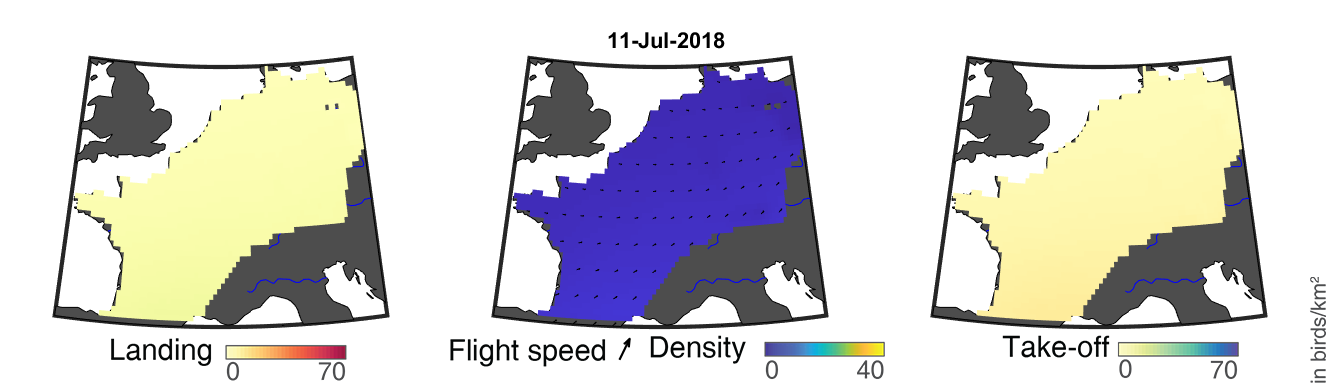

### day_149.png

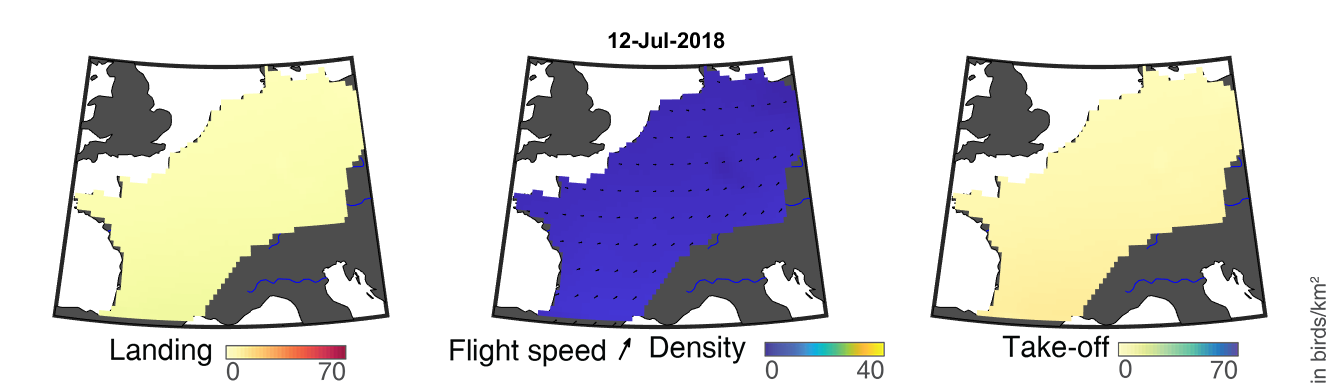

### day_150.png

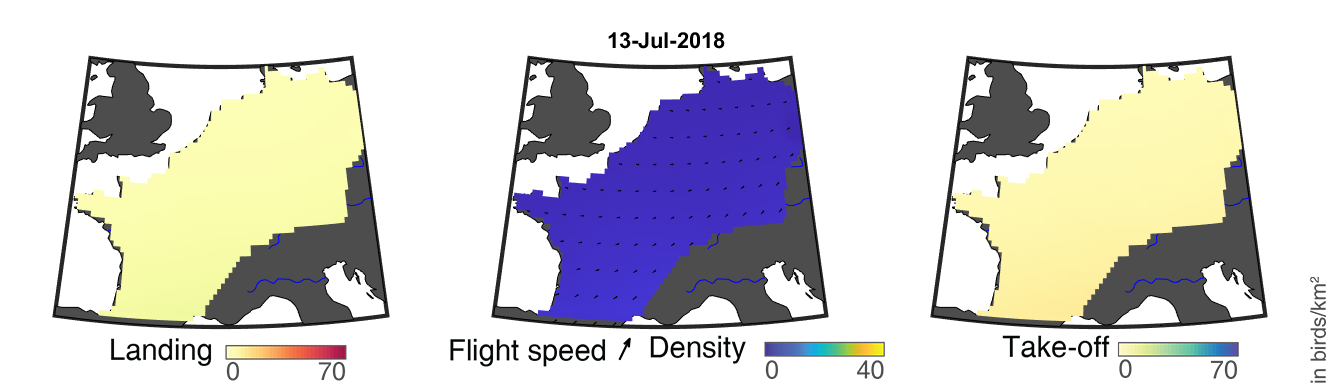

### day_151.png

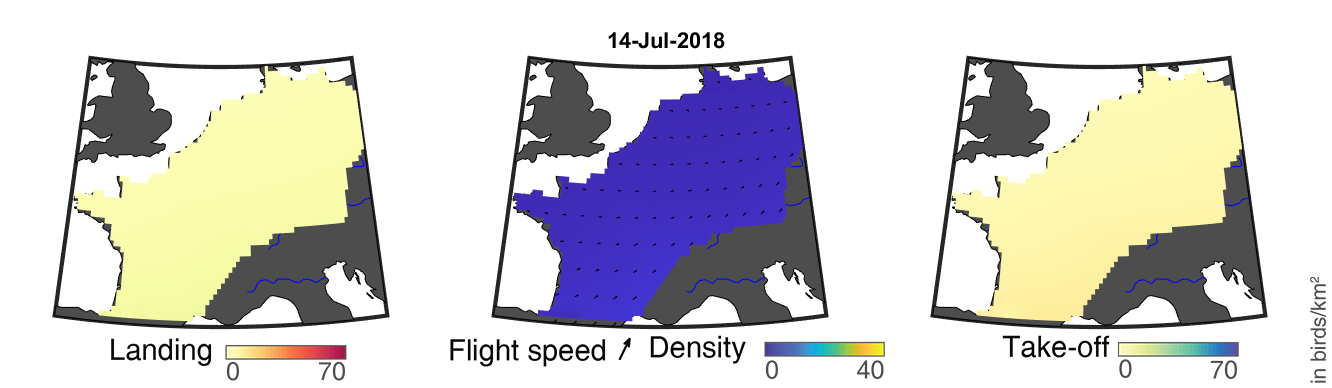

### day_152.png

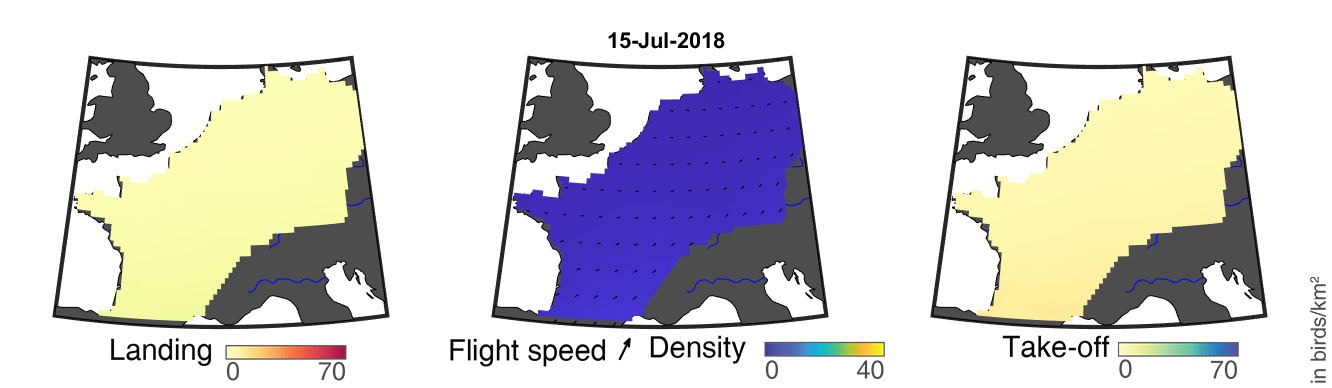

### day_153.png

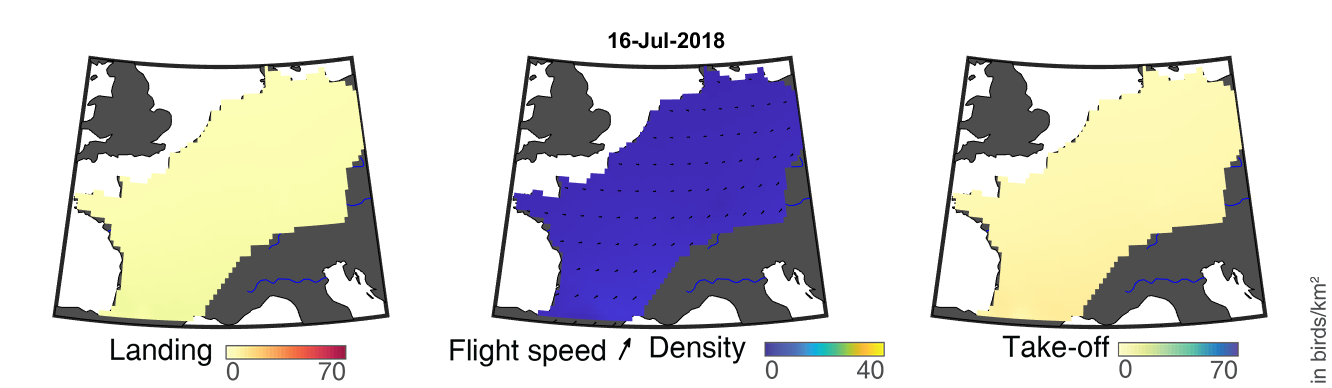

### day_154.png

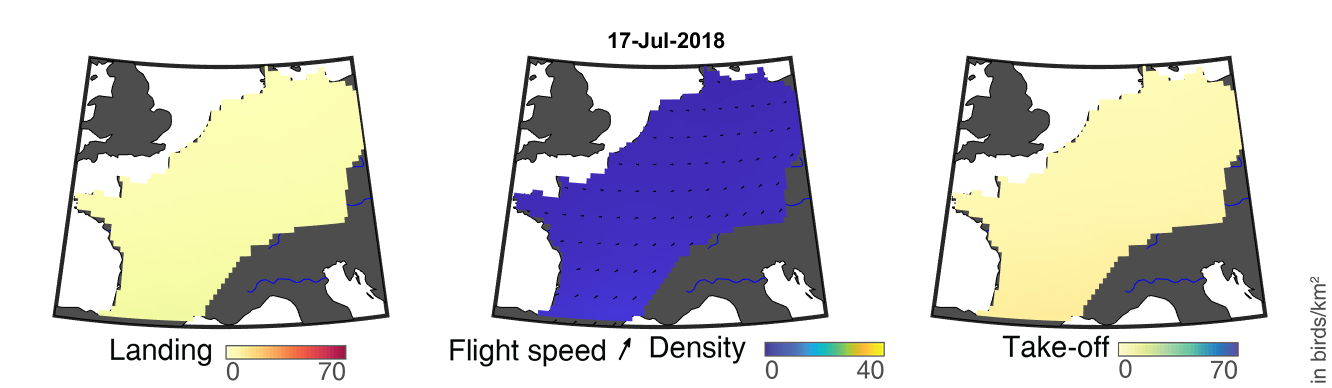

### day_155.png

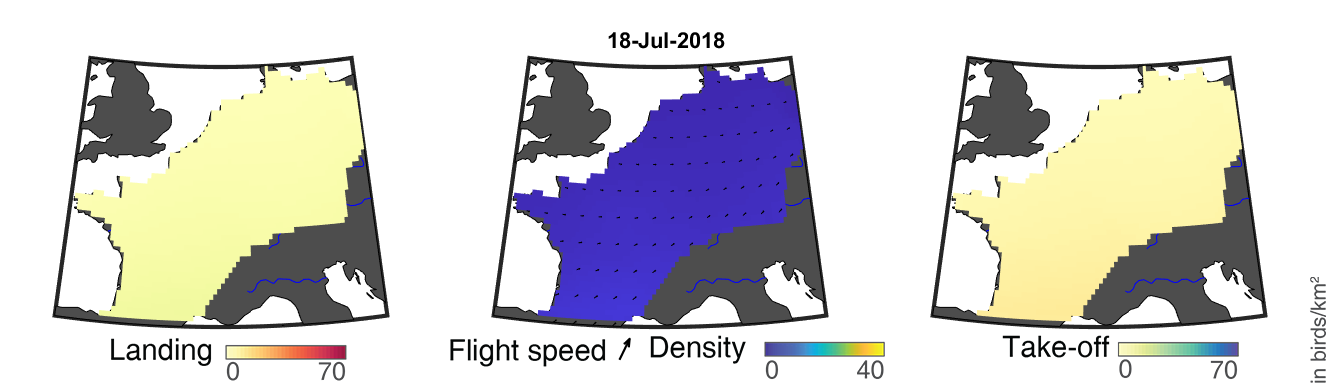

### day_156.png

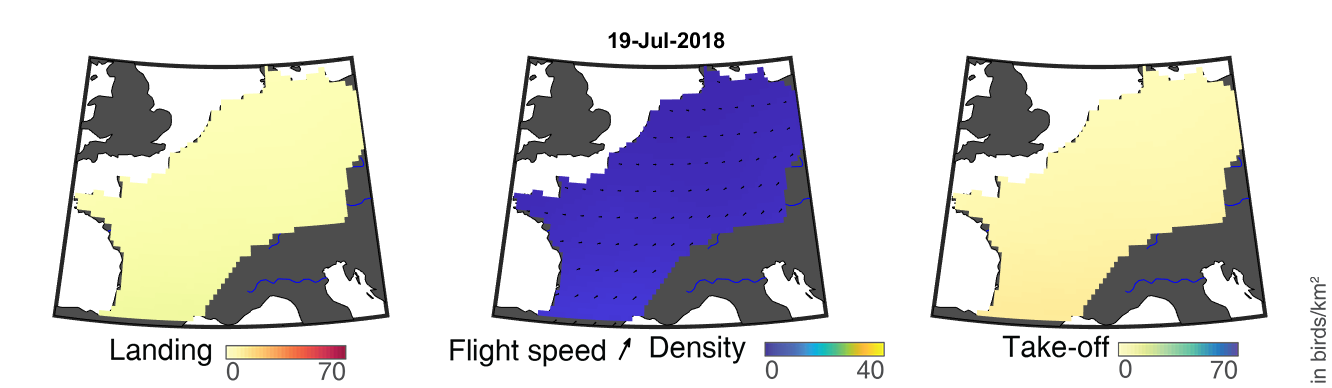

### day_157.png

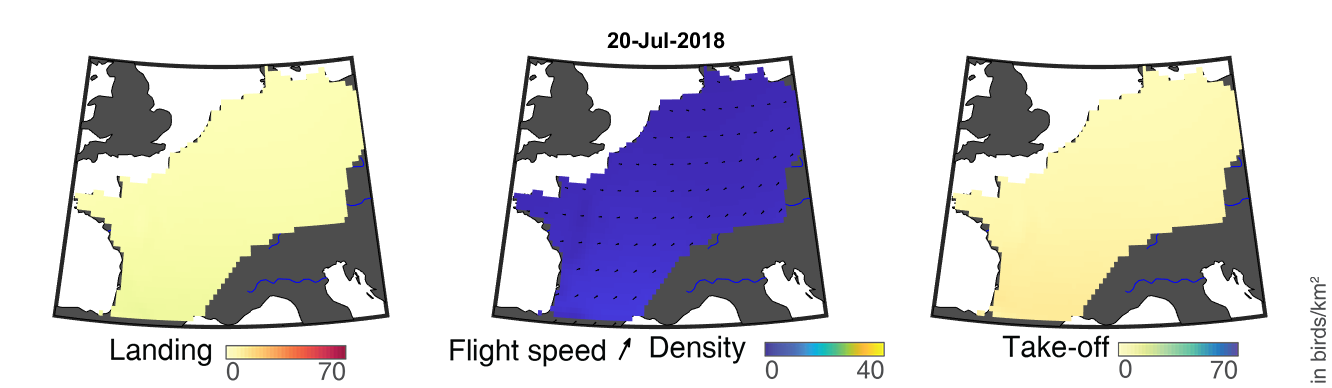

### day_158.png

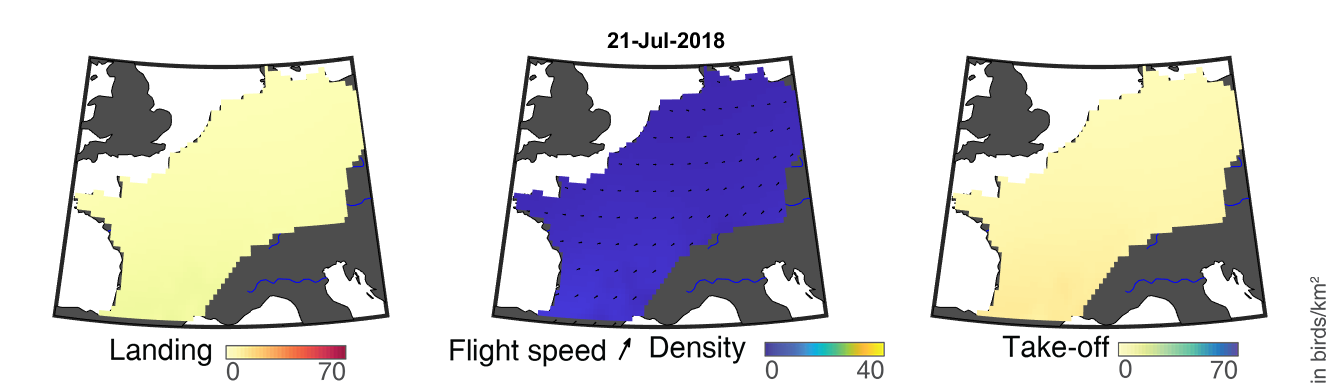

### day_159.png

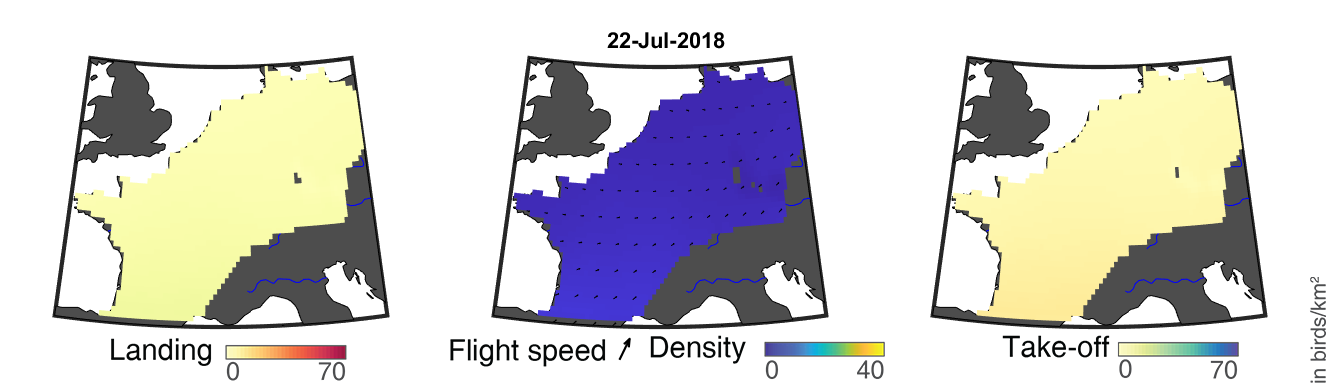

### day_160.png

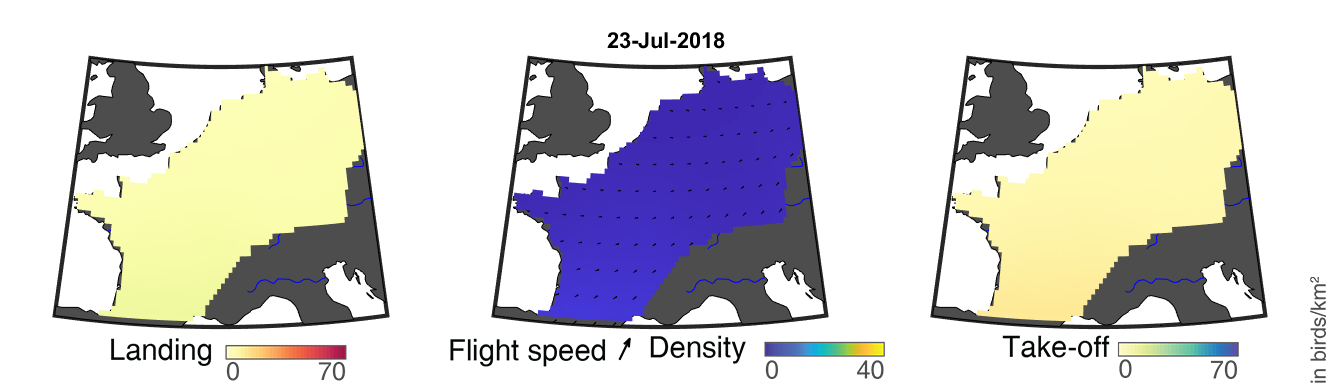

### day_161.png

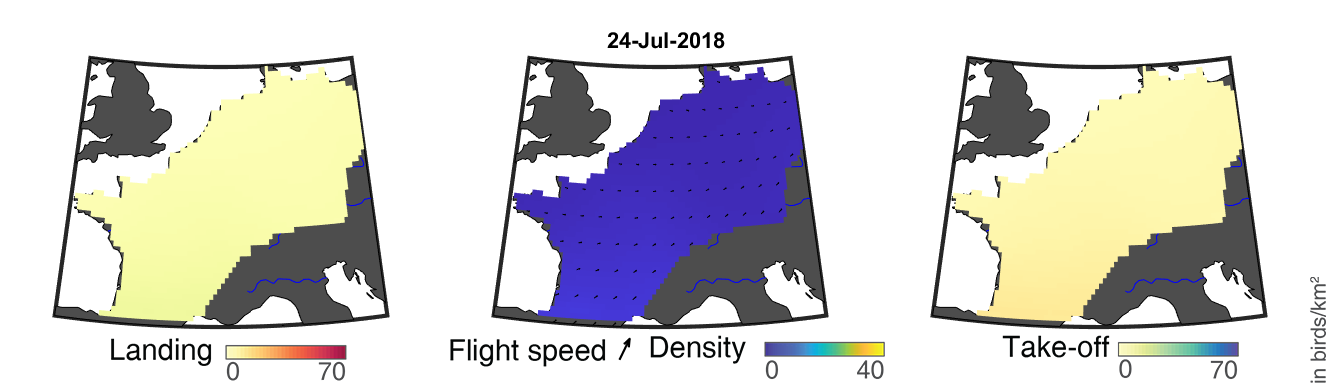

### day_162.png

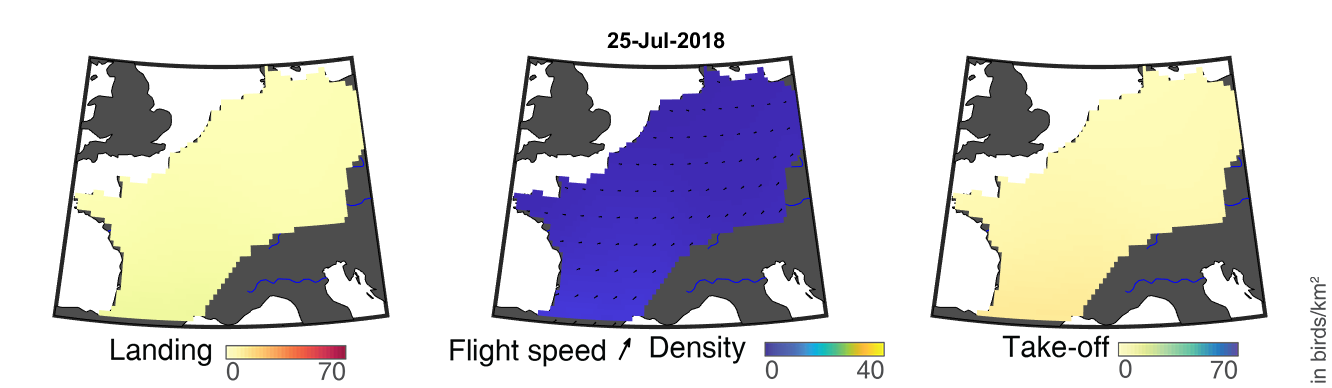

### day_163.png

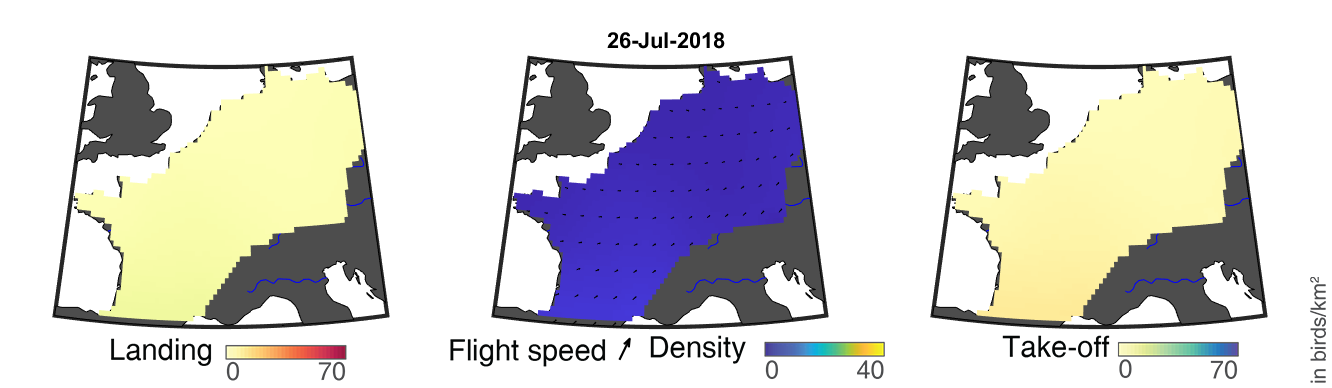

### day_164.png

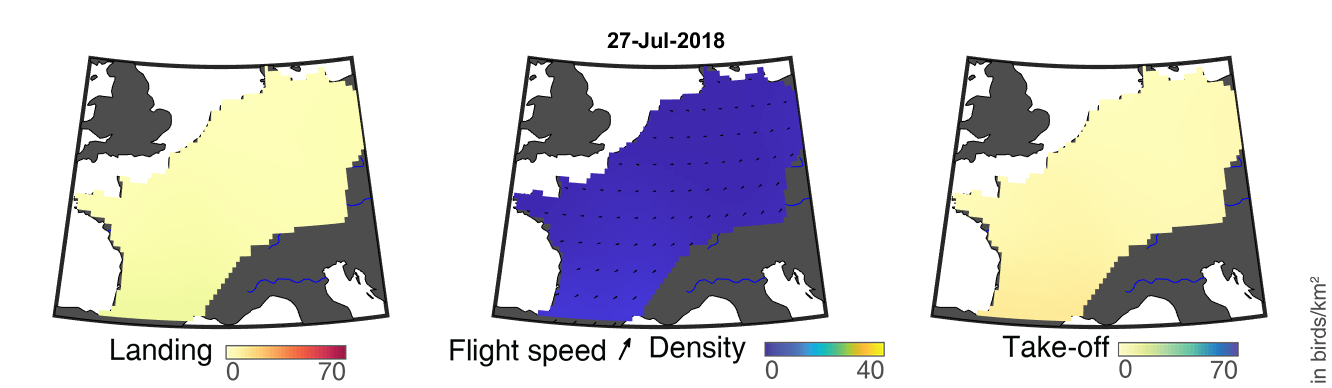

### day_165.png

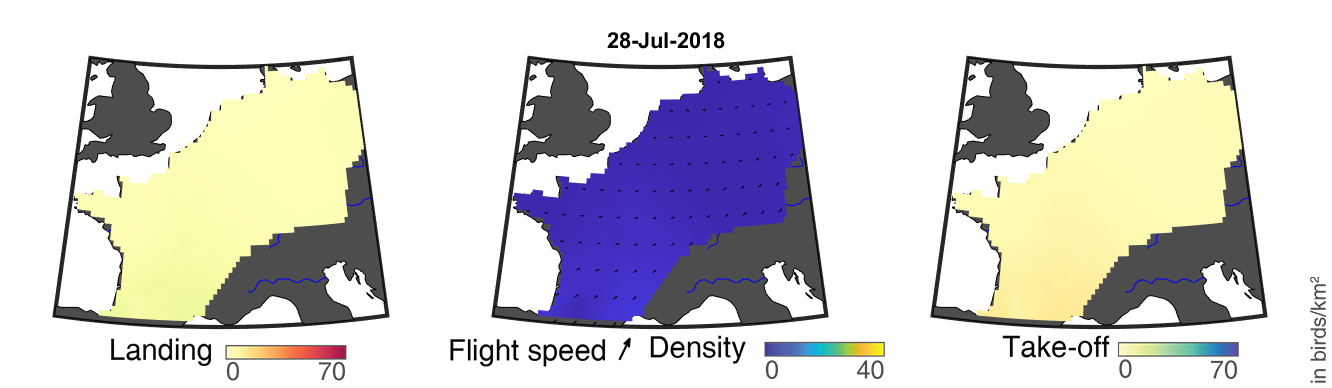

### day_166.png

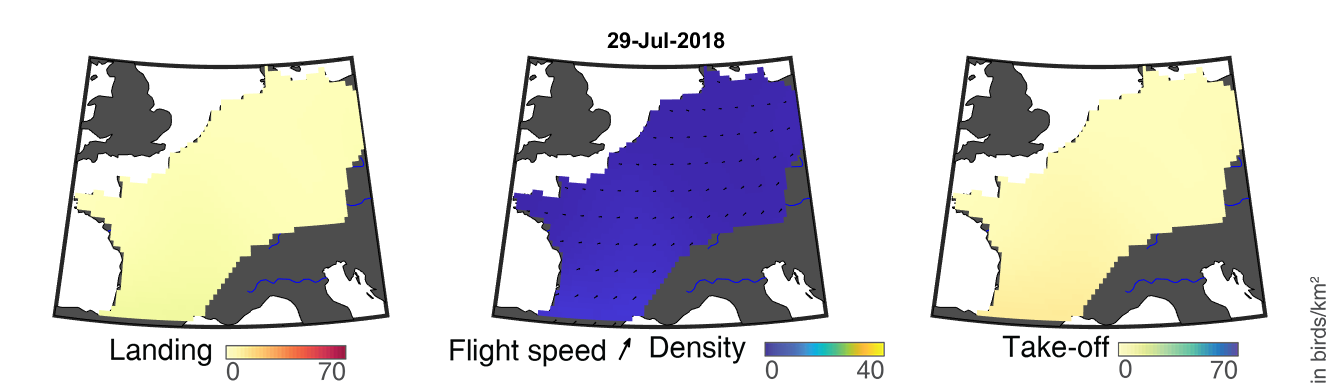

### day_167.png

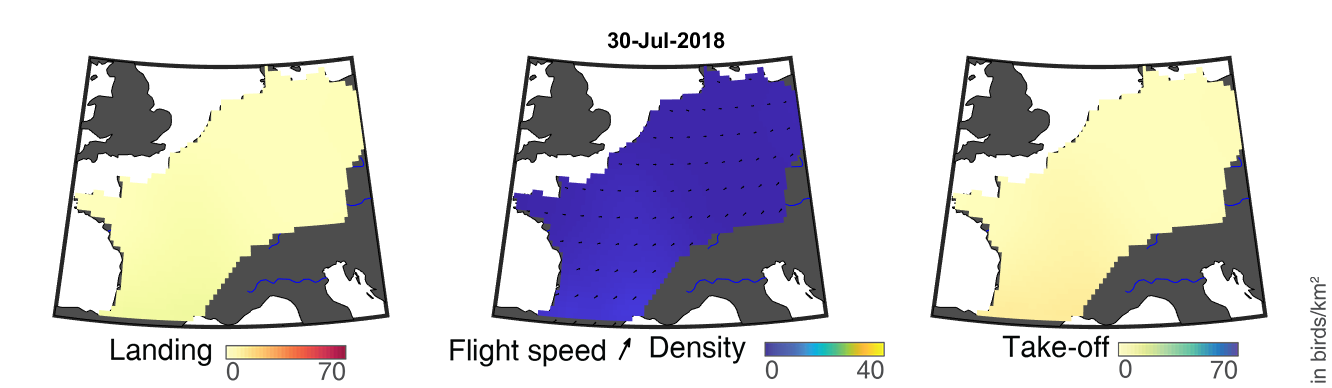

### day_168.png

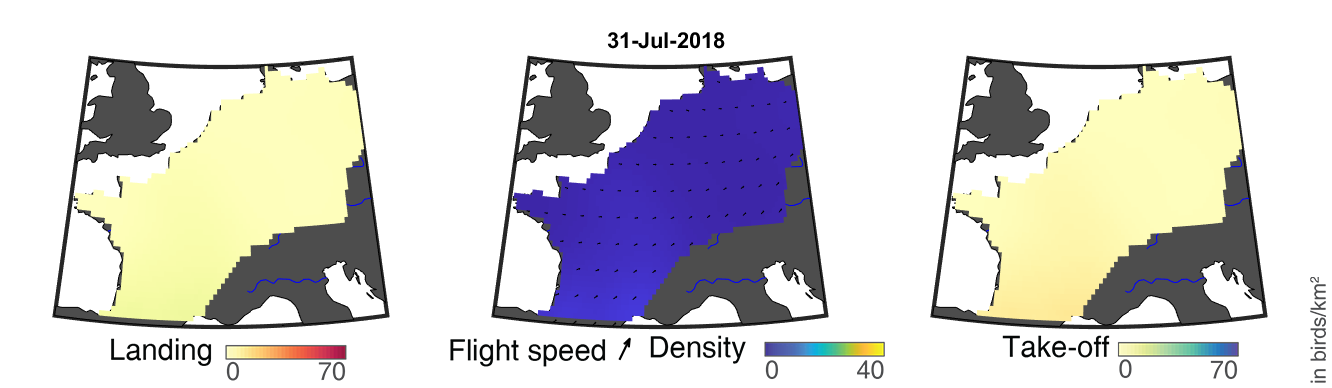

### day_169.png

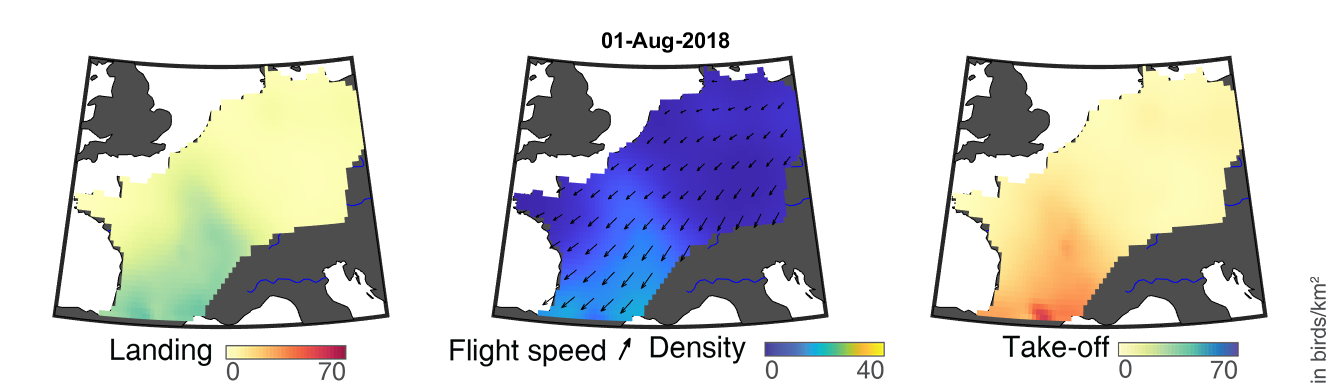

### day_170.png

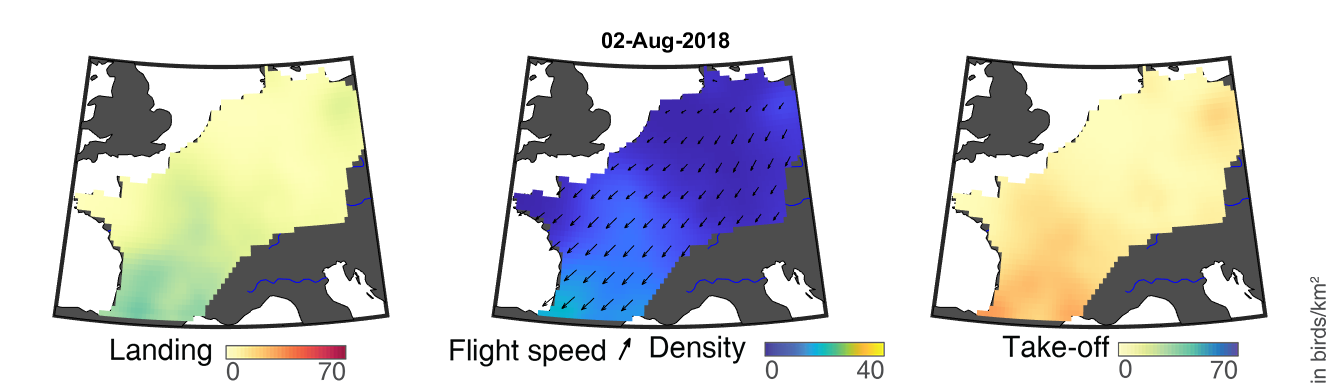

### day_171.png

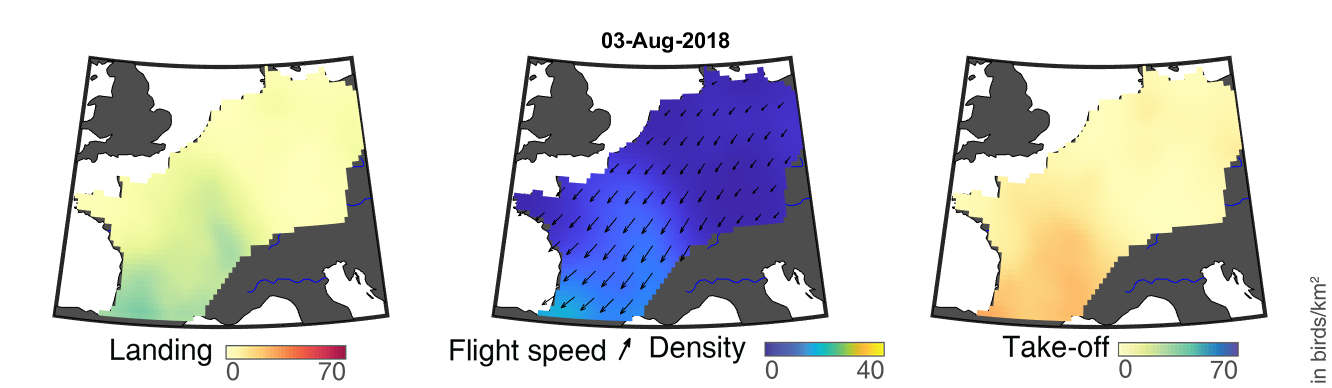

### day_172.png

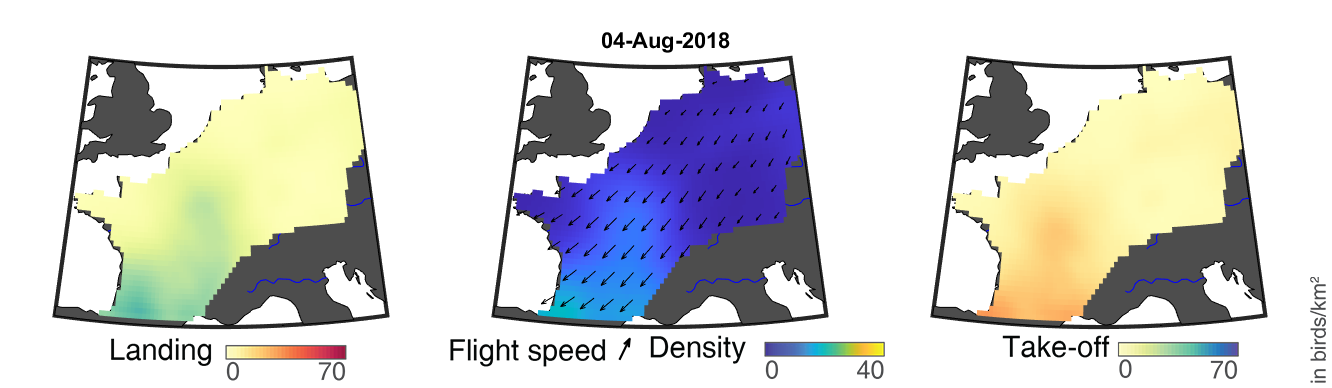

### day_173.png

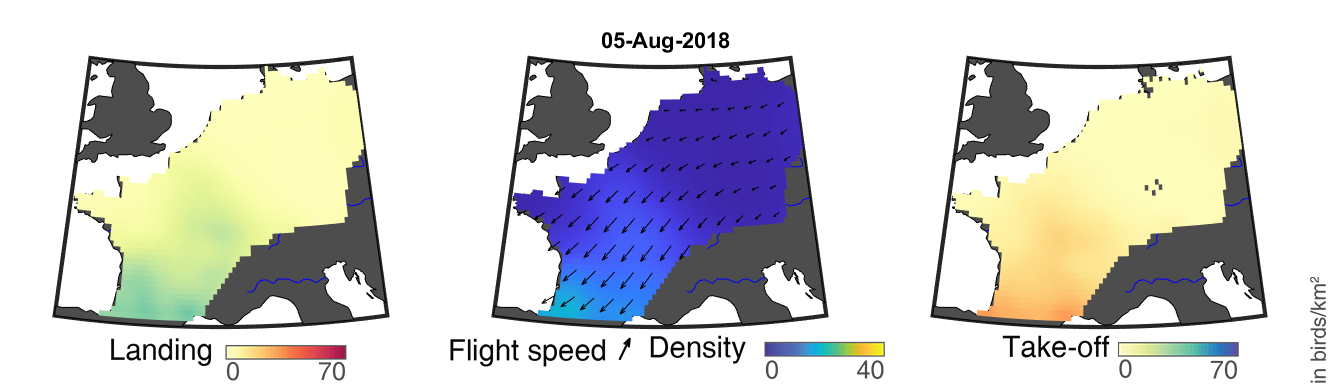

### day_174.png

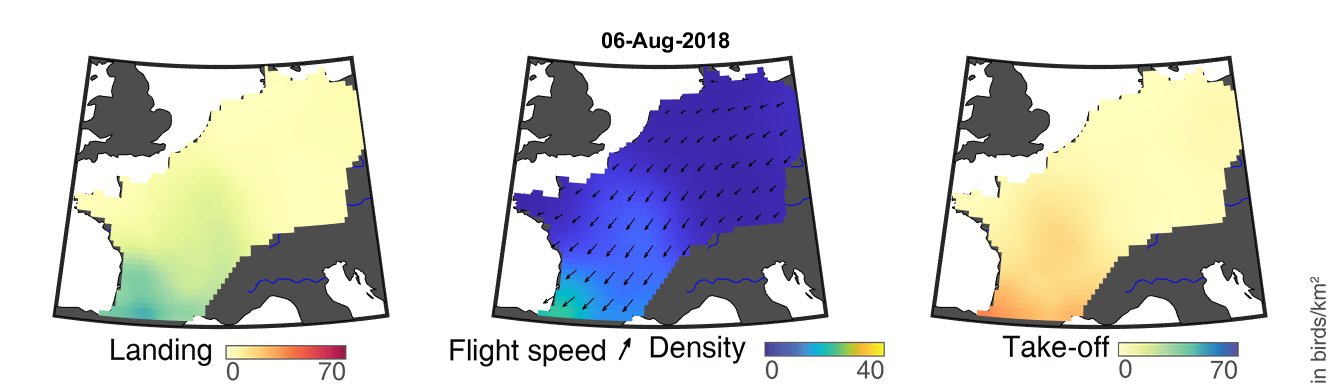

### day_175.png

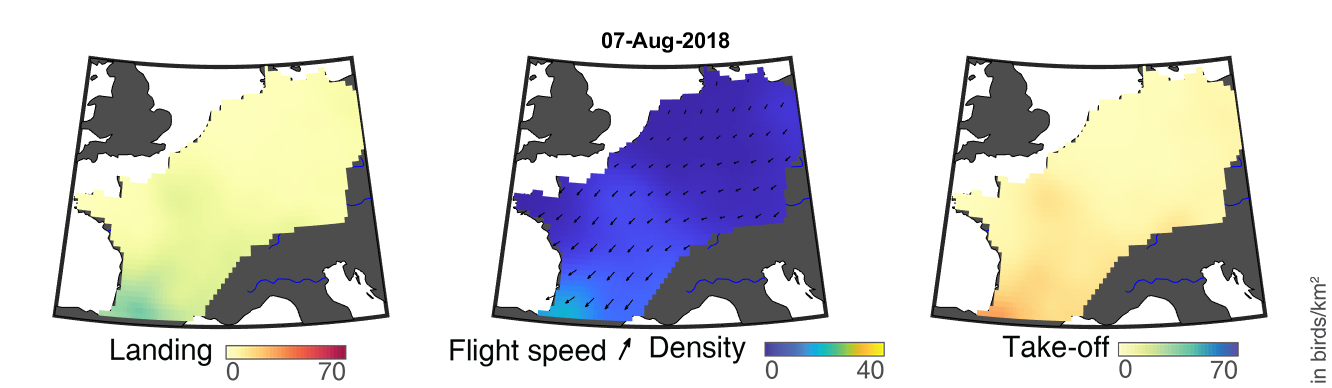

### day_176.png

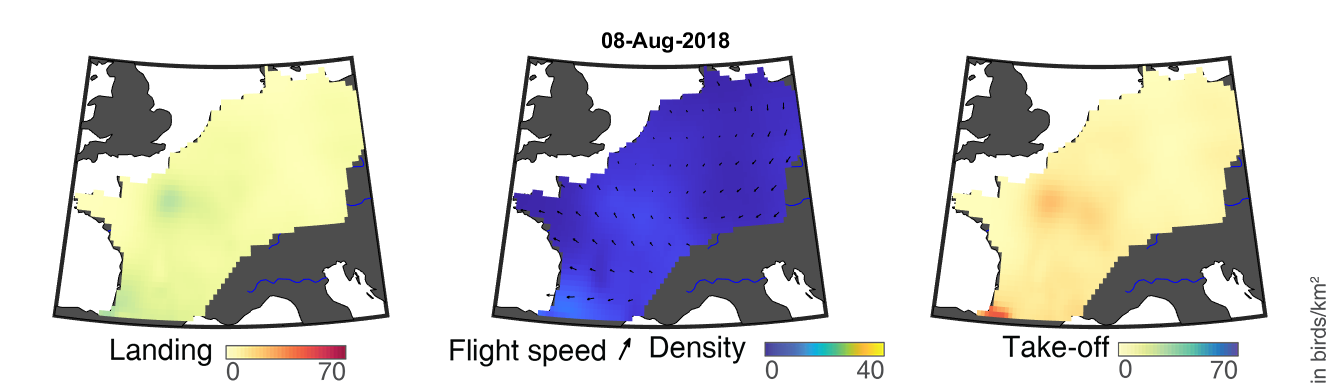

### day_177.png

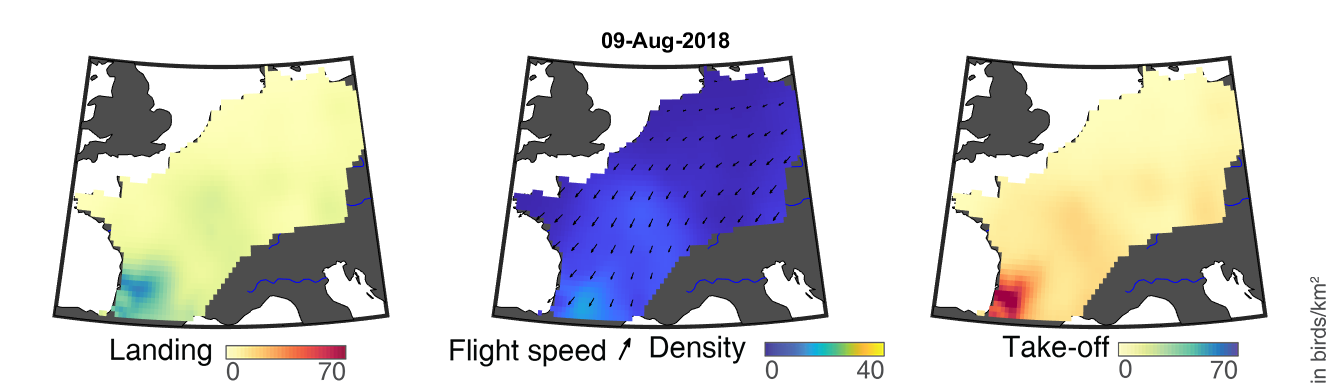

### day_178.png

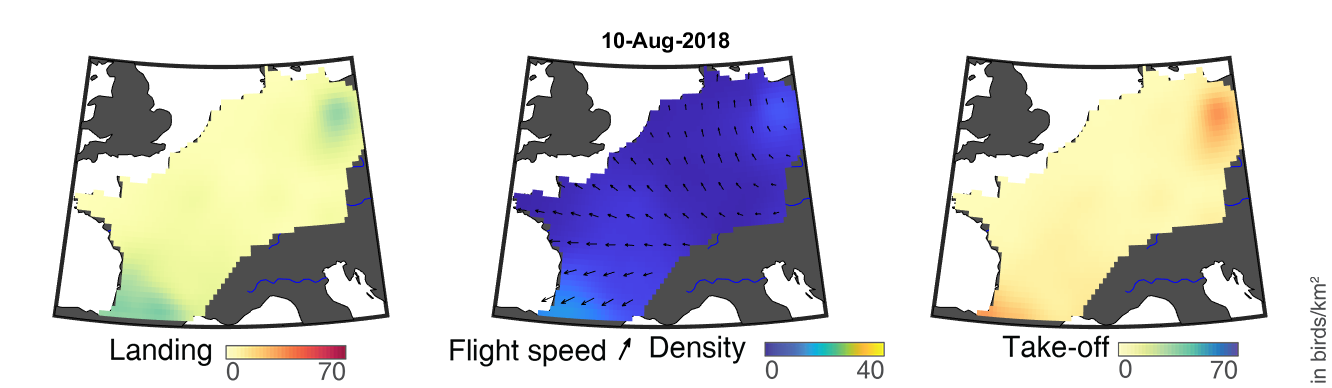

### day_179.png

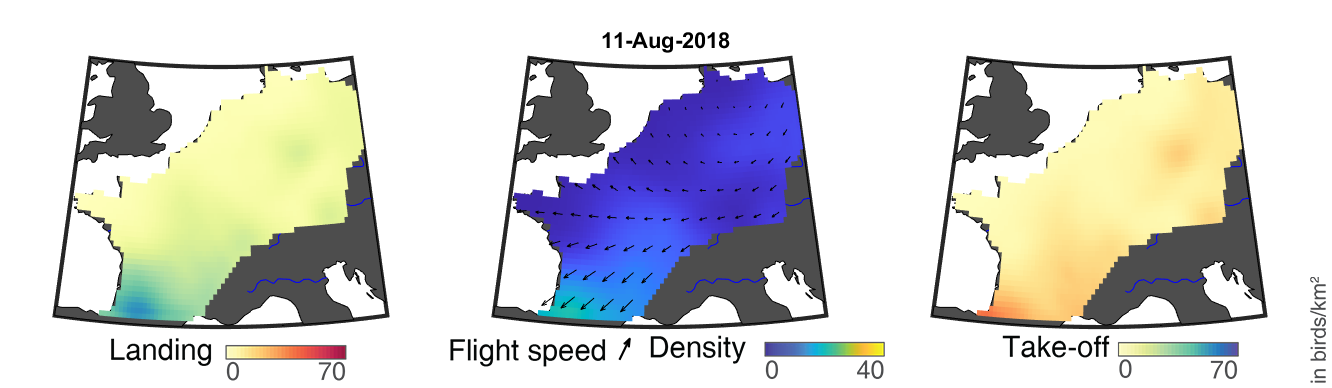

### day_180.png

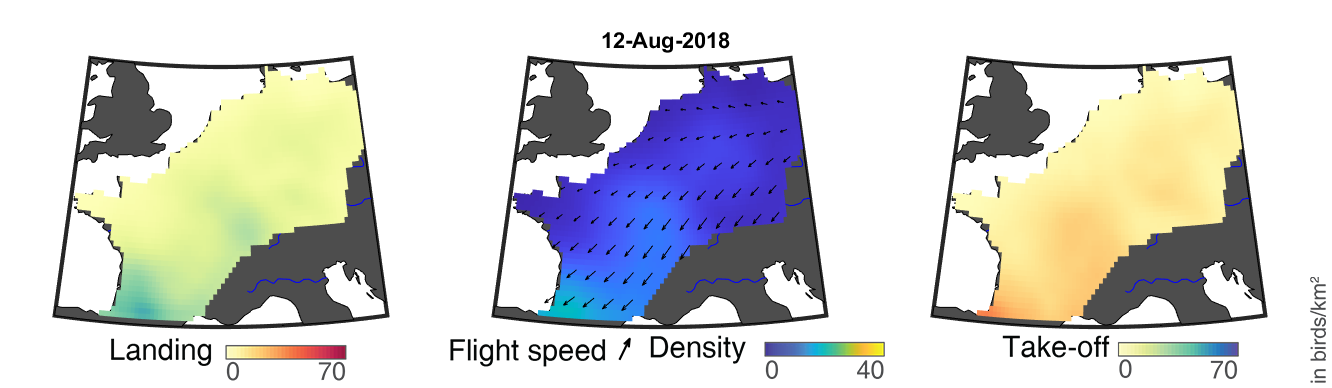

### day_181.png

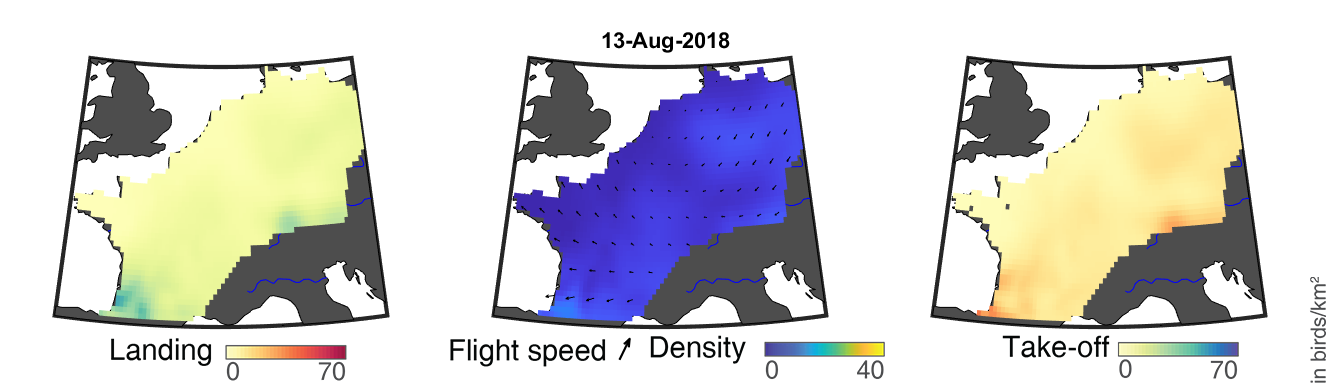

### day_182.png

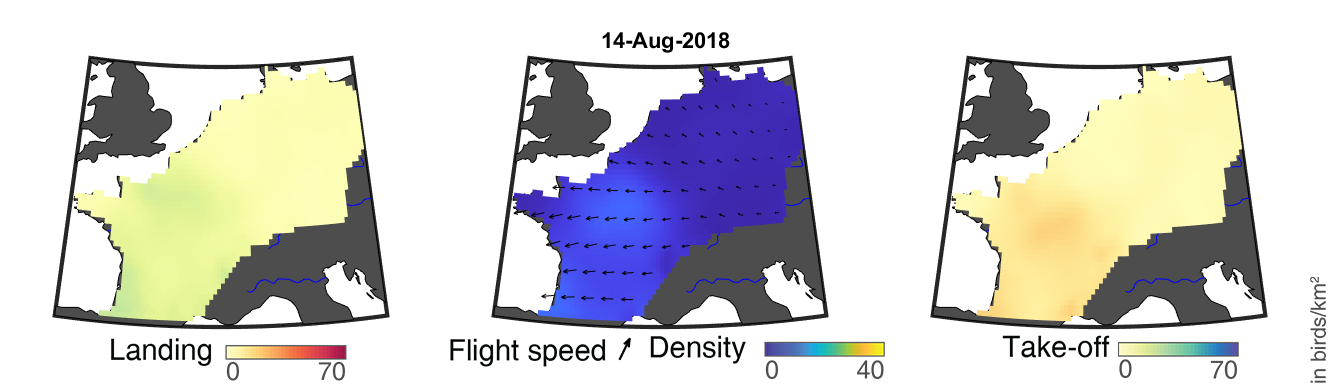

### day_183.png

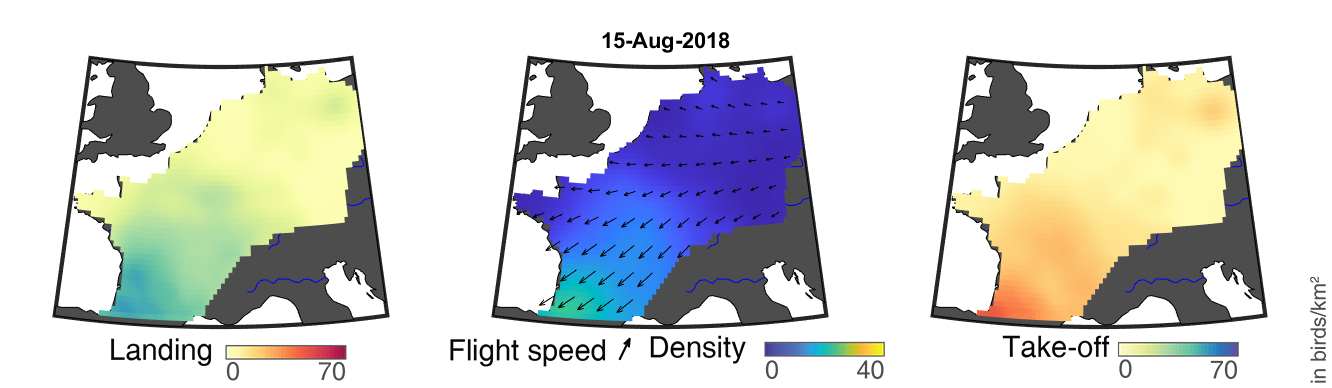

### day_184.png

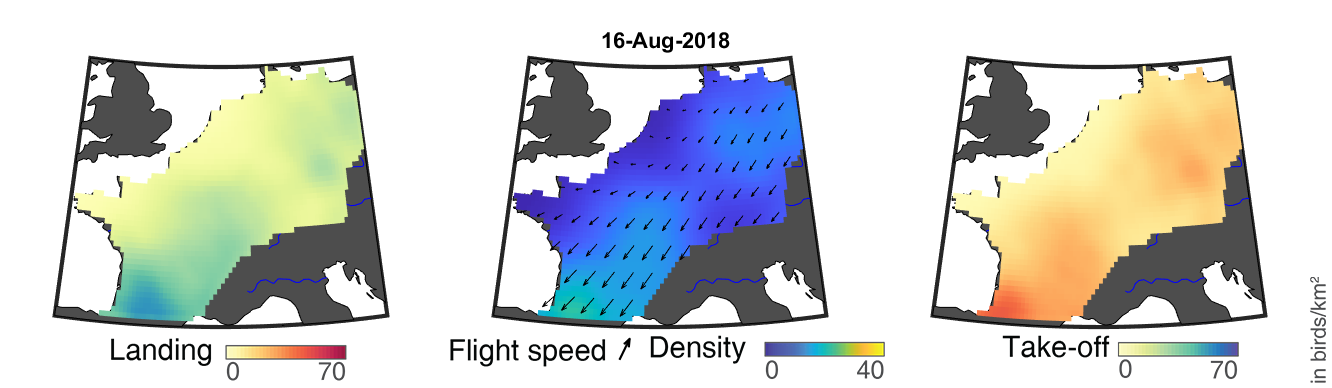

### day_185.png

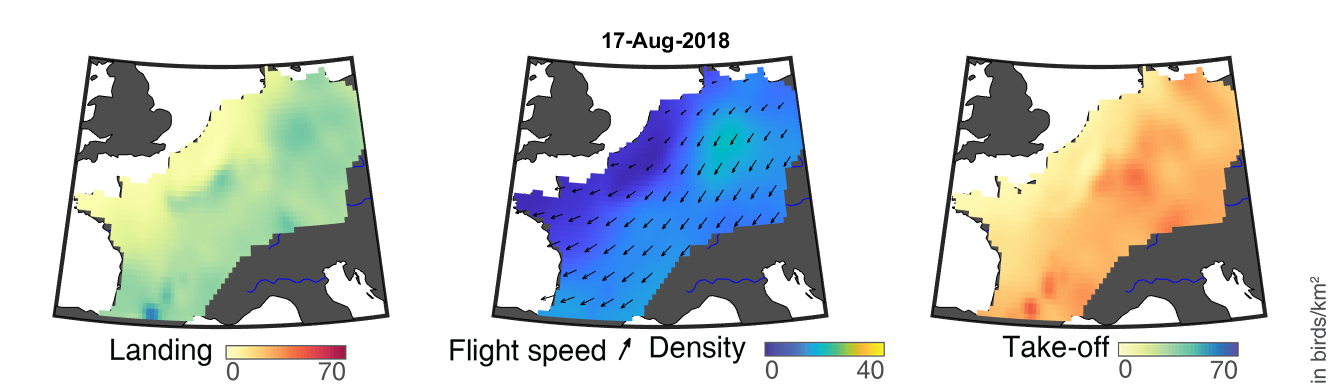

### day_186.png

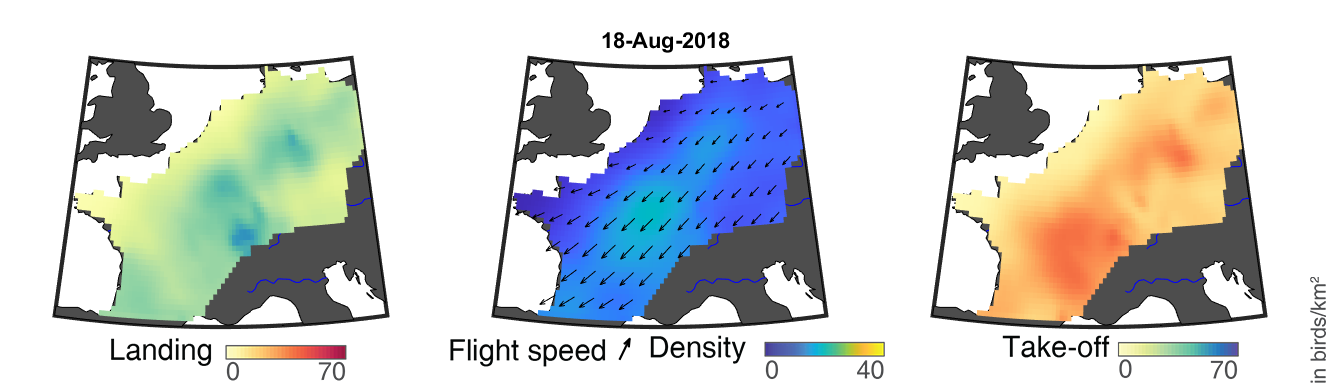

### day_187.png

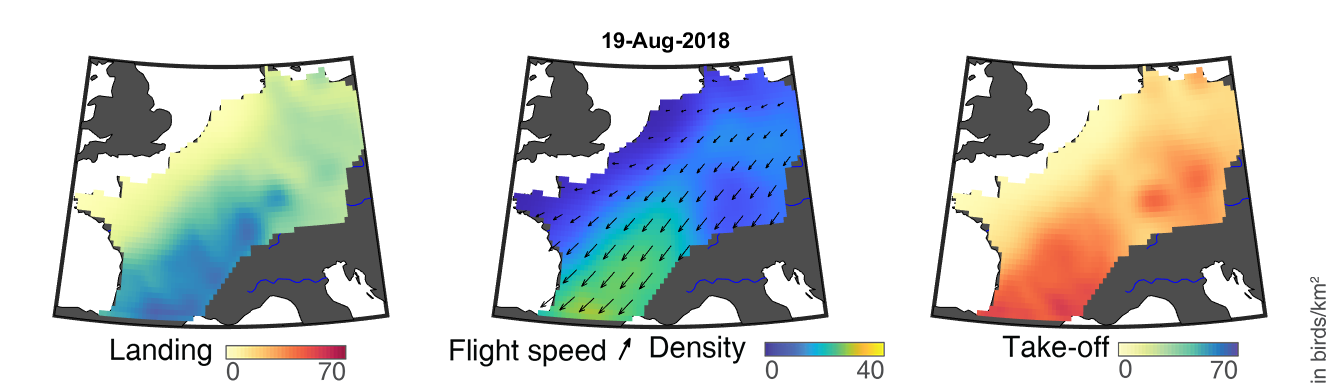

### day_188.png

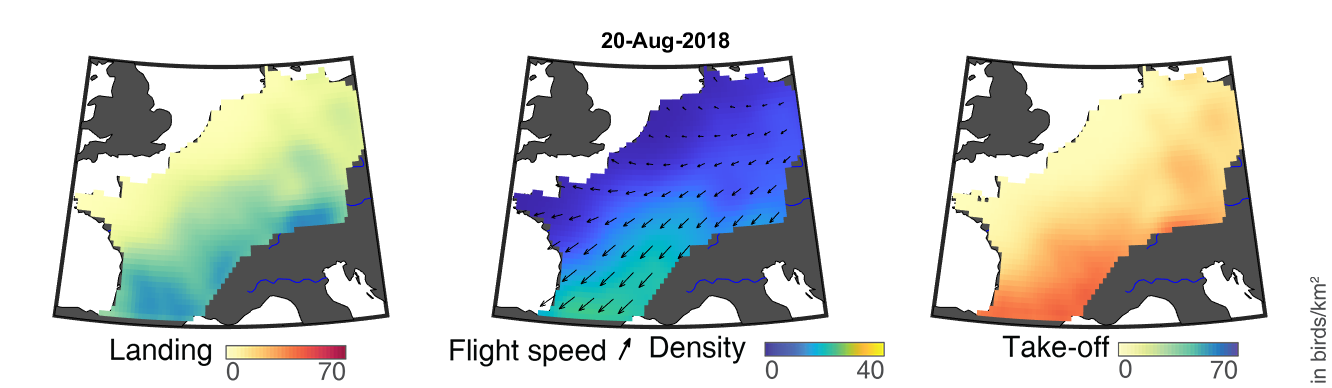

### day_189.png

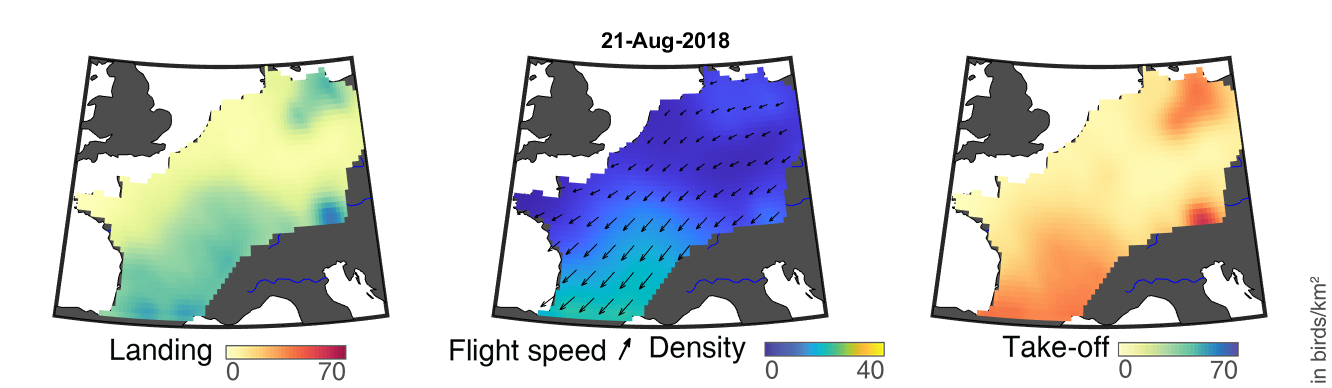
