## Supplementary Material for "Quantifying year-round nocturnal bird migration with a fluid dynamics model"

### Supplementary Material:
Modelling the flow of nocturnal bird migration with year-round European weather radar network.

Raphaël Nussbaumer, Lionel Benoit, Grégoire Mariethoz, Felix Liechti, Silke Bauer & Baptiste Schmid

#### 1 Data Pre-processing

We presents here the processing of the raw weather radar data, starting with the data selection (1.1), then explaining the steps of the data cleaning (1.2) and finally the method to transform the vertical profiles to time series (1.3). This procedure closely follows the processing performed in Nussbaumer et al. (2019) (Appendix A). Yet, we improved the removal of non-bird signal (1.2.3) and the vertical integration (1.3).

The MATLAB code corresponding to these steps can be found at www.rafnuss-postdoc.github.io/BMM/2018/#data-pre-processing. The resulting pre-process dataset is available at https://doi.org/10.5281/zenodo.3610184 (Nussbaumer, 2020).

##### 1.1 Data selection

We downloaded all available data from 1 January 2018 to 1 January 2019 at the finest resolution available of 5 minutes from the ENRAM repository (github.com/enram/data-repository) ENRAM (2020). After a visual quality check, only the data from radars in France, Germany, Belgium and the Netherlands were retained, totaling 37 radars. We limit the analysis to night-time as defined by the local sunrise and sunset at each radar location. We used the bird reflectivity ($\eta$) output from vol2bird rather than bird density because an automatic filtering is applied to the later. Bird reflectivity is later converted into bird density assuming a radar cross section of 11 cm$^2$ (Dokter et al., 2011). The East-West and North-South components of flight speed vectors are derived from the original flight speed and direction information.

#### 1.2 Data cleaning

##### 1.2.1 Bird density

Individual vertical bird density profiles are manually cleaned to remove unreliable observations (e.g. ground scattering, rain echoes) using a MATLAB dedicated GUI program which applies the method detailed in Nussbaumer et al. (2019) (Appendix A, points 2 and 3). In addition, a post-manual cleaning is performed by removing vertical profile data isolated by at least 30 minutes and removing noise by interpolating (log-linearly) the grid cells with a value greater than twice the mean of its eight direct neighbours.

##### 1.2.2 Flight speed

The vertical profiles of flight speed are cleaned as follow. Flight speed is only kept where bird density values are available, thus removing erroneous flight speeds due to rain and ground scattering. Gaps in flight speed are linearly interpolated when sufficient data is available around the gaps. This is defined with threshold of having at least 4 observations within a 20 min x 800 m neighborhood. Finally, the full vertical profile of flight speed at a single time step is only kept if the altitude bins where measurements are present covers at least the equivalent of 50% of the total number of birds over this profile.

##### 1.2.3 Low-reflectivity contamination

While non-bird signal with high reflectivity (e.g. rain) has been removed in the previous step, low-reflectivity signal contamination caused for instance by insects or snow echoes are still presents as it is difficult to identify manually. To remove these spurious signals, we use the difference in air speed and standard deviation in radial velocity to estimate the proportion of birds and non-bird reflectivity, correcting accordingly the bird density and flight speed (Nussbaumer, Schmid, Bauer, & Liechti, 2021).

#### 1.3 Vertical integration

After the data cleaning, the corrected vertical profiles are integrated vertically in order to convert the volumetric bird density [bird/km$^3$] into surface density [bird/km$^2$].

##### 1.3.1 Extrapolation to the ground

The bird density measured in the first altitude bins are often unknown or unreliable due to (1) ground scattering and (2) they are not (or partially) covered by the lowest scan of the weather radar. Yet they cover altitude where many birds might be migrating (e.g., Bruderer, Peter, & Korner-Nievergelt, 2018). In

Nussbaumer et al. (2019), we copied the bird density value of the first non-contaminated bins all the way to the ground (i.e. equivalent to vertical nearest-neighbour interpolation). However, since bird density usually increases near the ground (Figure 1.1) and thus represents a significant proportion of the total birds migrating, this process is prone to underestimate bird density.

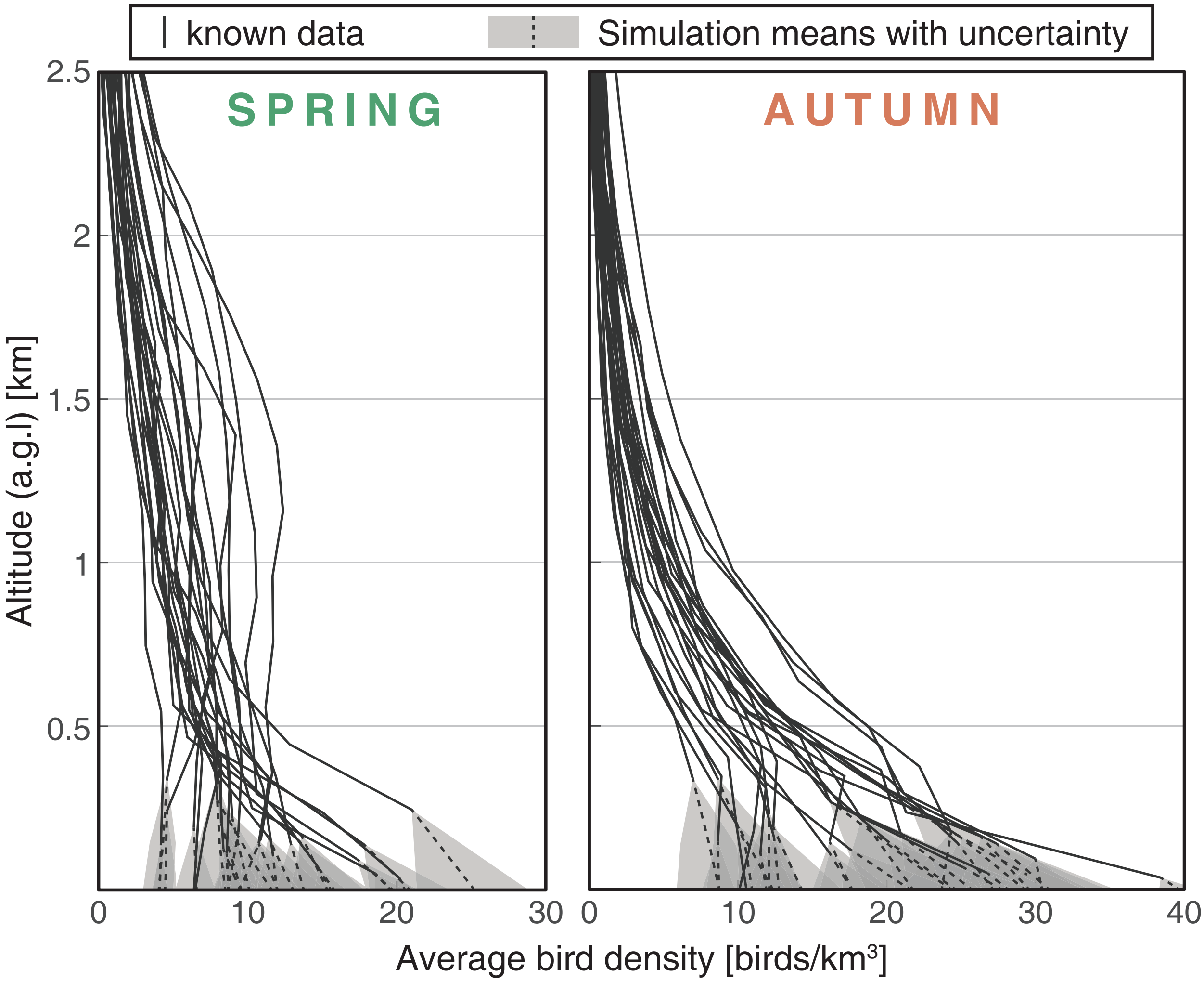


**Figure 1.1.** Average bird density vertical profile in spring (left) and autumn (right). Altitude is above ground level thus, the altitude of the data bins is different for each radar. The radars are unable to measure bird density in the lower elevation, resulting in gaps in the dataset. These gaps are filled with a simulation method based on "image patching" (see below for details). The resulting simulations are shown with a dotted line for the mean and as transparent area for the quantile 10 and 90.

To provide a more reliable estimate, we propose to resort to Multi-Point Statistic (MPS) (e.g., Mariethoz & Caers, 2014) to extrapolate low-altitude bins of bird density profiles. In short, MPS is a simulation method based on a pattern matching approach (see Figure 1.2). For each missing value of bird density, MPS searches for a similar pattern in the (space-time) neighborhood amongst the other radars and pastes the central value of one of the most similar patches in the simulation. The computation of the MPS simulation is performed with Quantile Sampling (QS) (Gravey & Mariethoz, 2019), using the following parameters (see www.gaia-unil.github.io/G2S/#QS for more details):

- Transformation: bird density data are log-transformed prior to MPS simulation..
- Simulation path: starts with the highest altitude, and simulates sequentially from high to low altitude.
- Training set: vertical profiles of all other radars within one week of the night of simulation. Only the first 1000m above radar elevation are used as training data.
- Neighbourhood size: 15 pixels with a uniform kernel of 49 x 7 cells (+/-2h, +/-500m).
- Number of best candidates to consider: 1.5.
- Number of realizations: 30.

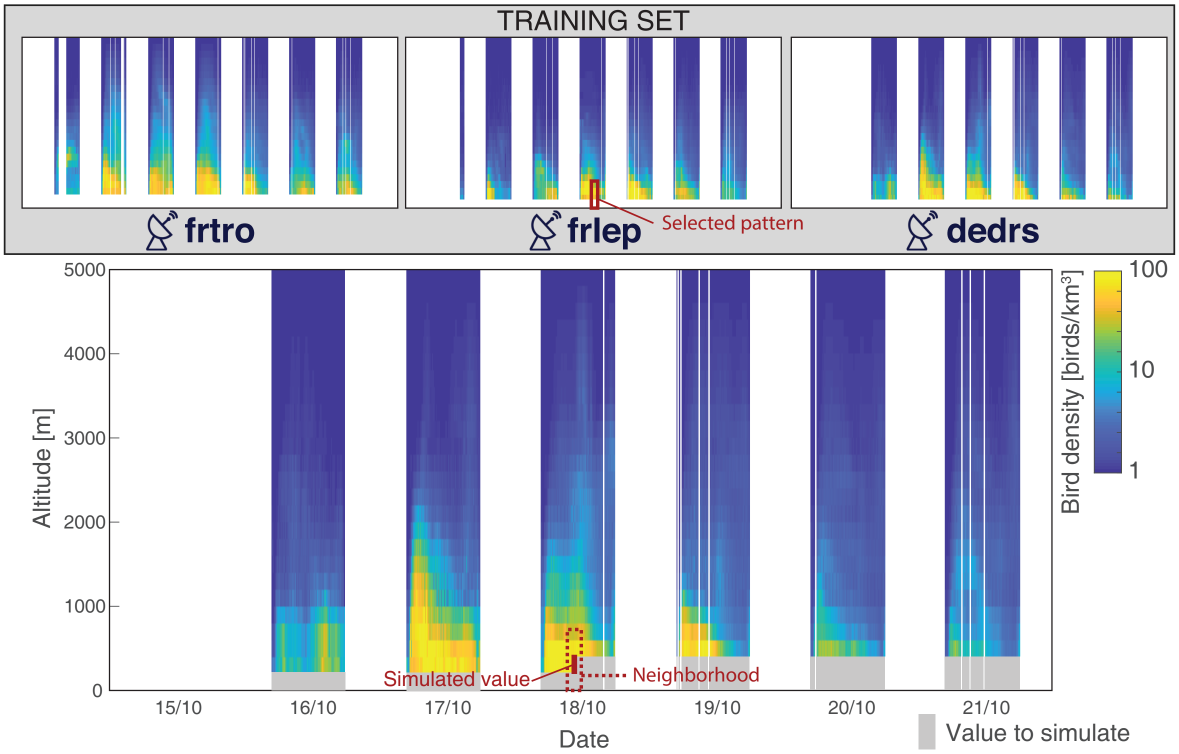


**Figure 1.2.** Illustration of the use of Multiple-Point simulation (MPS) for bird density extrapolation. For each value to simulate (in grey), a similar neighborhood is searched among the training set. When a matching pattern is selected, the corresponding pixel is pasted at the simulated value.

In addition to the extrapolation to the ground, we also performed an extrapolation of bird density up to 5000m a.s.l. This was performed by assuming a density of 0 bird/km$^3$ at 5000m and using a vertical log-linear interpolation.

##### 1.3.2 Local topography

Previously, bird density was integrated over elevations starting at the radar elevation up to 5000m asl. However, this approach is essentially assuming that no birds are flying in the volume below the radar elevation and within a 25km radius. Figure 1.4a illustrates for each weather radar, the altitude of the ground, the radar antenna, the first bins with acceptable data and the digital elevation profile surrounding the radar.

Here, we propose to refine this approach to take into account the local topography surrounding each radar (Figure 1.3). By taking advantage of the extrapolation performed in 1.2, we can estimate the density of birds below the radar elevation. In practice, we compute the volume of air according to a digital elevation model (i.e. the volume above ground ) for each elevation bin (cylinder of 200m height and 25km radius).

In addition, we also attempt to account for the effect that topography has on bird flight altitude. We assume that birds compensate their flight altitude based on the topography that they encounter against their flight direction. Therefore, we use the elevation profile encountered by birds (i.e. the projection of the topography on the plan perpendicular to their flight direction) to estimate the volume of air available for flight (Figure 1.3-2). This operation results in a correction of volume for each bin and each orientation of flight direction (Figure 1.3-3).

Flight speed is integrated vertically by weighting the average with the corresponding bird density which follows by the procedure mentioned above.

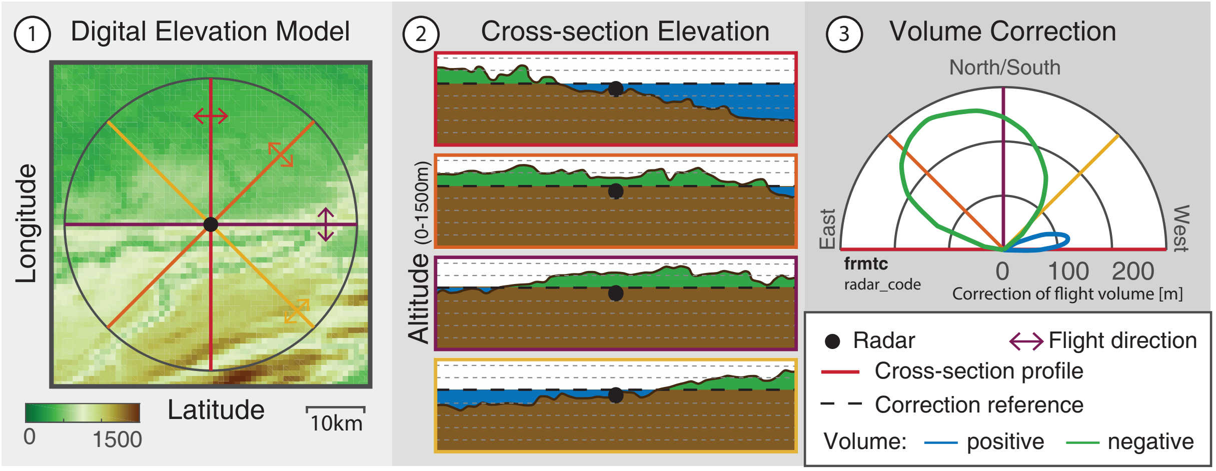


**Figure 1.3.** Difference of volume available for flight depending on the flight direction (0°is north and 90°is west.) Blue indicates a positive volume, and green a negative, compared to the previous method which used a flat topography located at radar altitude.

We illustrates in Figure 1.4 the general influence of the entire procedure. In Belgium, Netherlands and Germany, radars are generally located higher above ground (Figure 1.4a) which can create a small bias when integrating bird density from radar elevation rather than the ground. For most radar, the first bind with acceptable data (i.e. available and not contaminated) is the first bin above the ground elevation. In Figure 1.4b, we compare the resulting migratory flows (quantified by the migratory traffic rate (MTR)) to the previous method employed in (Nussbaumer et al., 2019). Generally, the difference of mean MTR is withing +/- 10% with a few exception when the data of several bins where missing above the ground level (e.g. nlhrw and nldhl) or when the radar was located at a high point compared to its surroudning topography (e.g. frniz). The difference between autumn and spring is mostly linked to the difference in flight altitude (Figure 1.1), but also due the difference orientation of flight (i.e. not only a difference in sens but also direction). The range of values generated by the simulations are relatively small compared to the variation among radars.

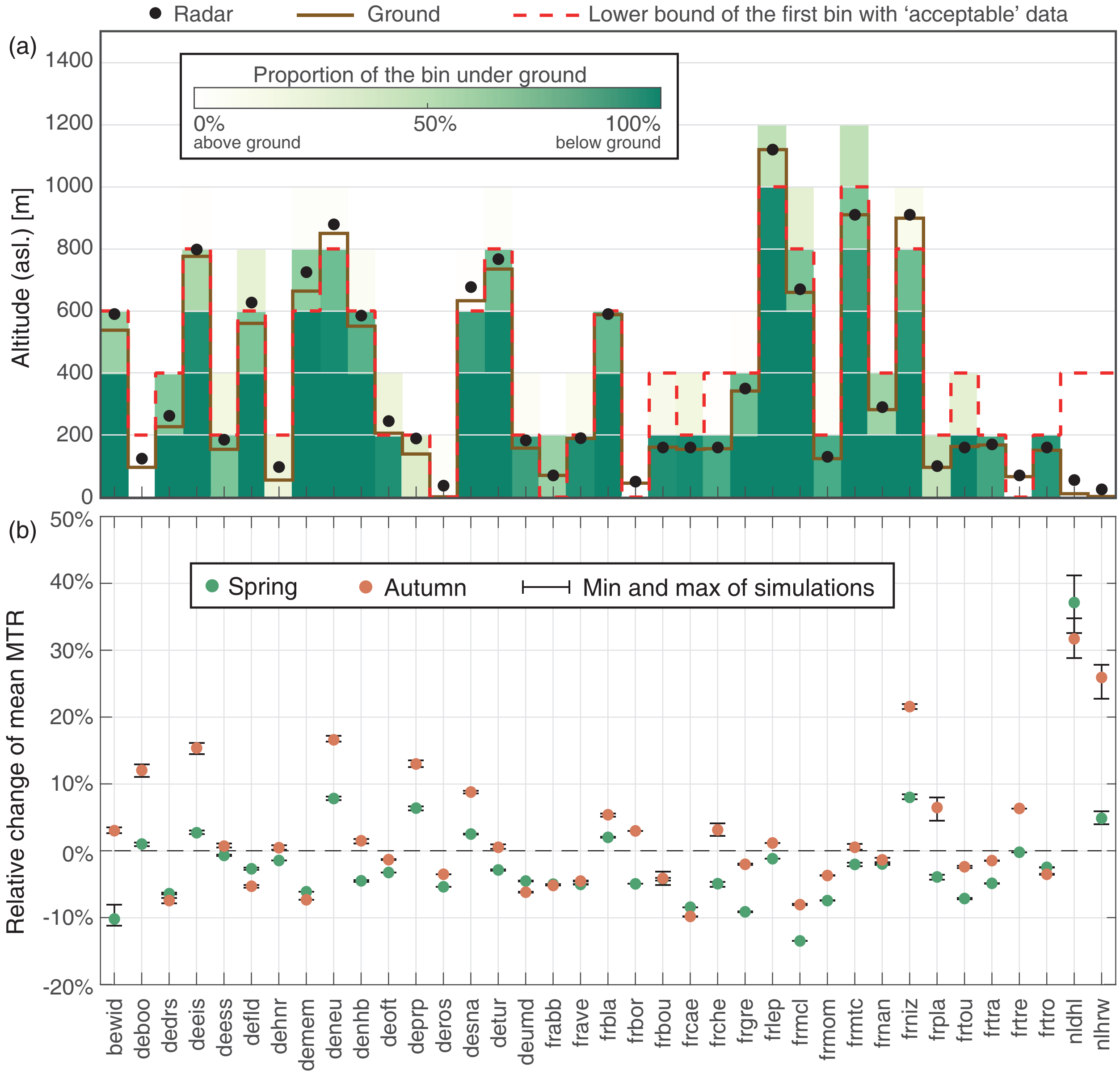


**Figure 1.4.** (a) Altitudinal coverage of the data, proportion of ground and elevation of the radar and ground. (b) Difference in the seasonal migratory traffic rate (MTR) between the new approach and the approached used in (Nussbaumer et al., 2019) (integration starting at radar elevation).

#### 2 Geostatistical interpolation of year-round bird migration data

The interpolation of bird density follows the methodology presented in Nussbaumer et al. (2019) with the following changes to handle year-round data:

- The power transformation of the density data is replaced by a quantile-quantile transformation.
- The non-stationarity of bird migration is accounted for by performing inference and simulation on four independent seasonal blocks (Figure 2.1).
- The (areal) bird density data produced by the pre-processing (1) consists of 30 simulated time series (see MPS simulation in section 1.3). The geostatistical interpolation takes advantage of knowing this range of possible values by using the corresponding uncertainty when fitting the trends of the geostatistical model (both temporal and spatial) with a generalized least square.
- In addition, a nugget effect is fitted to the covariance function of the kriging system to account for residual uncertainties of the data.

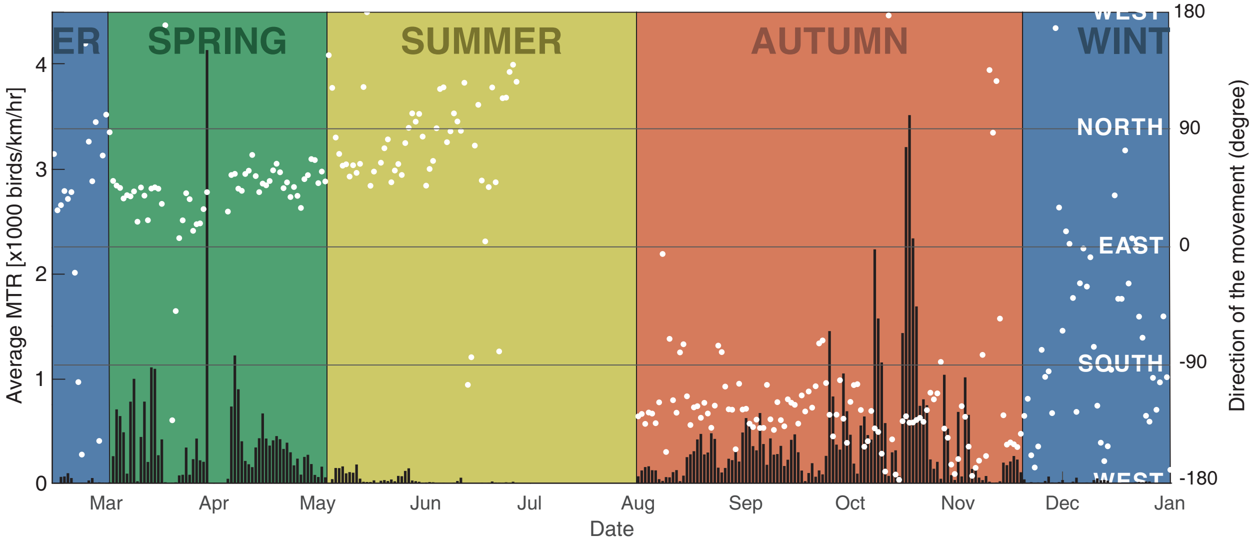


**Figure 2.1.** Summary of bird movement both in quantity and direction, based on the processed radar data. The average mean traffic rate (MTR) per night is shown as bar and the average weighted flight direction per night is illustrated with dots. This figure is used to separate the geostatistical model in four periods (winter, spring, summer and autumn), where the fit and estimation are done separately..

The interpolation of flight speed (both east-west and north-south components) is performed with a similar methodology than bird density with the following exceptions:

- No transformation is performed as the flight speed is close to normally distributed
- No trend was found in flight speed along the night (i.e. NNT). However, there was higher variability of flight speed at the beginning and end of the night, thus a variance trend was still performed.

More technical details of the interpolations performed are available on the github repository www.rafnuss-postdoc.github.io/BMM/2018/#interpolation.

#### 3 Complementary Figures

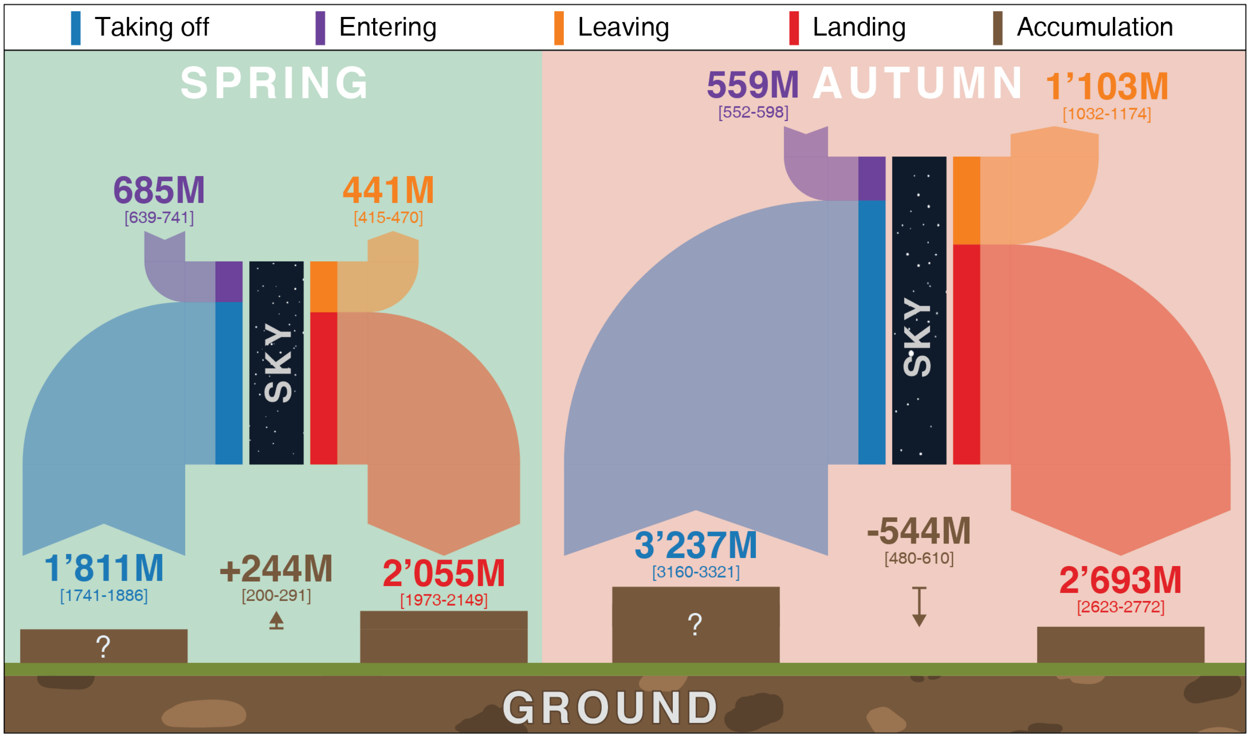


**Figure 3.1.** Seasonal fluxes of birds transiting through the sky in millions (M) with Q5-Q95 uncertainty ranges. The mass balance approach ensures that the number of birds entering and taking-off is equal to those leaving and landing. The taking-off and landing number are summed over the entire domain and season, in other words the same bird can be counted multiple time. While the change in number of the ground can be estimated, the initial number (denoted with a question mark) is unknown for both spring and autumn. Note that the number of birds entering and leaving slightly differ from Figure 4 because here, we show the cumulative numbers per grid cell, whereas Figure 4 sums the fluxes over each entire transects flux thus, neutralising cells with fluxes of opposite sign within a transect.

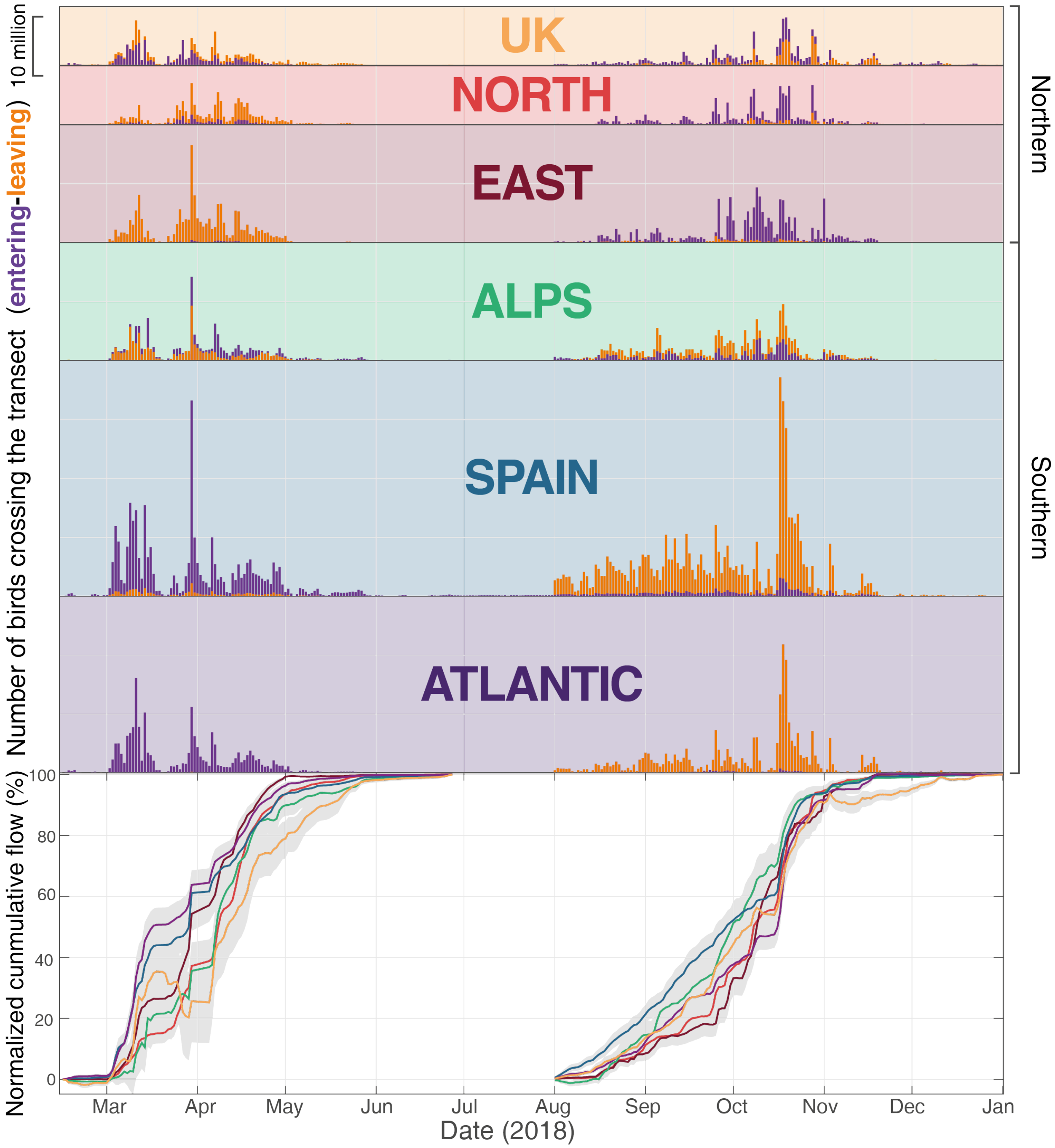


**Figure 3.2.** Nightly fluxes of birds entering (purple) and leaving (orange) the study area through the different transects. (bottom) The normalized cumulative flow is the cumulative sum of the daily flows (entering + leaving) normalized by the total seasonal flow (thus adding to 1) for spring and autumn separately.
